# Genomic exaptation and regulatory landscape shifts as key mechanisms enabling flatworm terrestrialization

**DOI:** 10.1101/2025.01.09.632089

**Authors:** Lisandra Benítez-Álvarez, Raquel García-Vernet, Klara Eleftheriadi, Carlos Vargas-Chávez, Nuria Escudero, Judit Salces-Ortiz, Iñaki Rojo, Fernando Ángel Fernández-Álvarez, Cristina Chiva, Eduard Sabidó, Eduardo Mateos, Fernando Carbayo, Rosa Fernández

## Abstract

Understanding the genomic toolkit that facilitated animal terrestrialization—the transition from aquatic to terrestrial environments—is crucial for unravelling the evolutionary processes behind the origin and diversification of terrestrial biodiversity. Despite its significance, the genomic foundations driving the physiological and metabolic adaptations required for life on land remain largely unexplored across most terrestrial animal phyla. Planarians (phylum Platyhelminthes) represent an ideal model for studying terrestrialization, as only one terrestrial lineage, the family Geoplanidae (order Tricladida) is known to exist. Here, we used an integrative approach combining genomics, transcriptomics, and proteomics to investigate the genetic underpinnings potentially facilitating adaptation to terrestrial environments. Our analysis revealed a significant burst of gene gain preceding the diversification of terrestrial planarians and their split from freshwater relatives, in the branch leading to Tricladida. Upon exposure to abiotic stress, terrestrial and freshwater planarians exhibited distinct genetic toolkits: most differentially expressed genes emerged in orthologous groups gained specifically in the branch leading to Tricladida, over half of which showed signs of strong directional selection in terrestrial flatworms, indicating their adaptive importance for land colonization. Transcriptomic analyses further revealed contrasting stress responses: terrestrial planarians upregulated ancient genes whose origin predates the Geoplanidae lineage to cope with abiotic stress, while freshwater planarians downregulated a separate set of ancestral genes. Our genomic, transcriptomic, and proteomic data consistently show that the genetic toolkit for abiotic stress response in terrestrial planarians is highly differentiated from that of their freshwater counterparts, with significant regulatory shifts as well. Overall, our findings suggest a burst of gene gain in the Tricladida lineage, with co-option of these genes, rather than clade-specific innovations, playing a critical role in the origin and diversification of terrestrial flatworms. This underscores genomic exaptation and regulatory landscape shifts as key mechanisms enabling terrestrialization within Platyhelminthes. This study offers the first genome-wide insight into the genetic toolkit underlying flatworm terrestrialization and contributes broadly to understanding the genomic basis of animal terrestrialization.

## Introduction

The colonization of land by animals is without doubt one of the most important episodes in the animal evolutionary chronicle. The ancestors of extant terrestrial arthropods, nematodes, nemerteans, velvet worms and mollusks, among other phyla, managed to transition from marine to terrestrial environments and successfully adapt to life on land. For that, deep physiological and metabolic changes were needed, as abiotic conditions significantly differ on land and in the sea. This probably came coupled with a profound genomic reshaping to adapt to the new abiotic challenges, including osmoregulation, oxygen homeostasis, locomotion, reproduction, new pathogens to defend from or different sources of food, among others. While some studies have dealt with understanding the genomic changes facilitating this transition in some animal phyla, such as mollusks (Sun et al. 2019; Liu et al. 2021; Krug et al. 2022; Aristide and Fernández 2023), vertebrates (Bi et al. 2021; Meyer et al. 2021) or arthropods (Sharma 2017; Asano et al. 2019; Veldsman et al. 2021), little attention has been paid to some of the animal phyla with terrestrial forms. Such is the case of the land planarians, within the phylum Platyhelminthes. They constitute an excellent model to explore the genomic basis of animal terrestrialization as all described terrestrial forms are within a single clade, the family Geoplanidae. Being sister to families and superfamilies compromising freshwater forms, it is clear that the transition from marine to terrestrial environments in platyhelminthes occurred through the freshwater route: within the order Tricladida, we find a suborder composed mostly by marine species (suborder Maricola), a clade composed of freshwater and terrestrial species (suborder Continenticola) (Sluys et al. 2009) and a third clade comprising freshwater species inhabiting in epigean and hypogean habitats (suborder Cavernicola), with the interrelationships between the three clades being dubious due to incongruences between ribosomal markers (Benítez-Álvarez et al. 2020; Stocchino et al. 2021) and transcriptomic data (Vila-Farré et al. 2023). Land planarians of the family Geoplanidae and its sister group of freshwater planarians, the Dugesiidae, are classified with the superfamily Geoplanoidea in the suborder Continenticola (Sluys et al. 2009).

The reshaping of the gene repertoire through time via gene gain, duplication and loss is a well-known mechanism facilitating adaptation and innovation, as it can provide the raw material for adaptation by introducing new or modified functions that help organisms cope with new challenges. As new coding genetic pieces arise through gene gain or duplication, new functions may arise as well associated with them (Fernández and Gabaldón 2020; Ocaña-Pallarès et al. 2022; Merényi et al. 2023). Furthermore, evolutionary changes in gene repertoires can lead to the diversification of gene functions. This diversification allows organisms to exploit new ecological niches, utilise different resources, or develop resistance to new pathogens or environmental stressors (Wen et al. 2023; Pal and Andersson 2024). As for gene loss, it has been proven a powerful mechanism for genomic innovation as well, as gene interactions and regulatory networks may be strongly impacted, resulting in the reshaping of the functioning gene circuits (Jiménez-Marín et al. 2023; Domazet-Lošo et al. 2024).

Gene gain and duplication has been shown to facilitate adaptation to new environments through several mechanisms, including neofunctionalization (i.e., it creates gene redundancy, allowing one gene copy to maintain its original function while the other evolves new functions), or subfunctionalization (i.e., the original function is divided between the copies) (Braasch et al. 2018; Sandve et al. 2018; Birchler and Yang 2022). This genetic redundancy increases genetic variation, providing a broader substrate for natural selection and enhancing the potential for beneficial mutations while buffering against harmful mutations, as the extra gene copy can often compensate for deleterious changes (Robinson et al. 2023; Xie et al. 2023). Gene duplication can also lead to increased gene expression levels, which may be advantageous in certain environmental conditions, and supports the evolution of complex regulatory networks and novel developmental pathways, contributing potentially to significant morphological innovations (Li et al. 2020; Van de Peer et al. 2021; Veitia and Birchler 2022). These processes can drive rapid speciation and diversification, enabling organisms to explore and thrive in new and changing environments. However, gene repertoire evolutionary dynamics within animal phyla and its role in animal terrestrialization is still largely unexplored.

Here, we aim at unveiling the genomic toolkit that facilitated the colonization of land in platyhelminthes through a combination of phylogenomics, transcriptomics and proteomics, with emphasis on revealing gene repertoire evolutionary dynamics that may have been instrumental for adaptation to terrestrial environments. For that, we sequenced the first high quality transcriptomes to date of multiple terrestrial flatworms (15 species) and explored the evolutionary dynamics of the encoded genes to understand how gene gain, duplication and loss may have contributed to their evolutionary success in living on land. We further tested which genes may be putatively involved in response to abiotic stress related to living on land, confirmed their presence at the protein level with proteomics, and explored their evolutionary origin within the Platyhelminthes phylogeny. Our results revealed a burst of genomic innovation prior to the origin and diversification of terrestrial planarians, confirmed by transcriptomics and proteomics analyses, therefore pointing to genomic exaptation as a key facilitator of the colonisation of land environments in platyhelminthes. Our results provide the first genome-wide evidence of the toolkit involved in flatworm terrestrialization, and more broadly contribute to delineating the big picture of the genomic basis of animal terrestrialization.

## Materials and Methods

An overall representation of the analytical workflow used in this study is shown in Fig. 1.

**Fig. 1.**
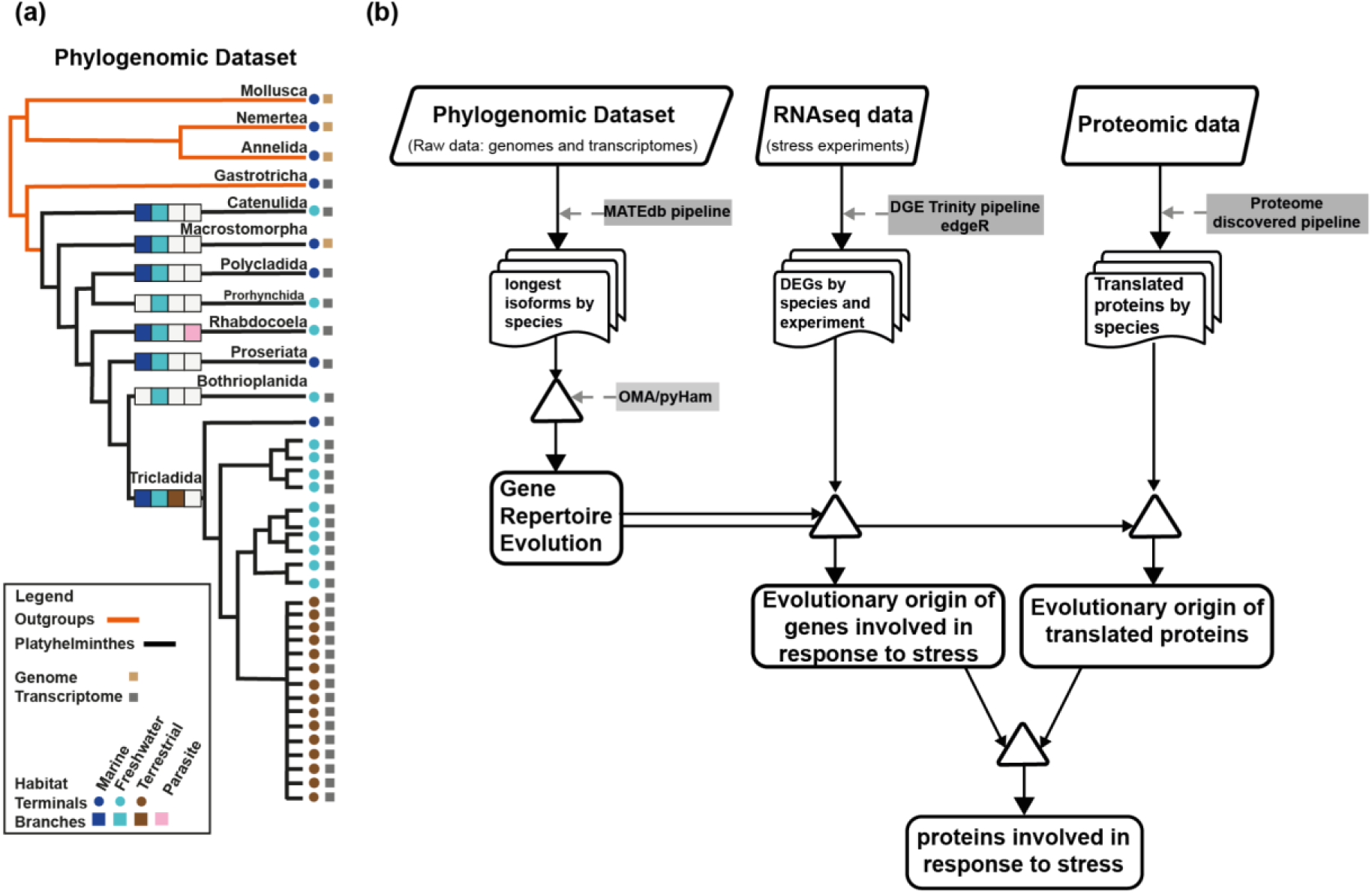
Multiomics analytical workflow of this study. a) Schematic species tree showing the phylogenomic dataset. Outgroups are represented with orange branches and Platyhelminthes with black ones. Habitats reported for each flatworm lineage are represented by squares in the branches, and the habitat of each species is indicated by circles in the terminals (dark blue: marine, light blue: freshwater, dark brown: terrestrial, light pink: parasite). Tree topology and habitats extracted from Laumer et al. (2015). The type of data is represented by squares in the terminals (transcriptome, grey, or genome, yellow). b) Workflow followed in this study integrating genomic, transcriptomic and proteomics data.

### Sampling and sequencing of terrestrial planarians

We hand-collected 15 species of terrestrial planarians from Spain and Brazil. Sampled species, coordinates and localities are available at Supplementary Table S1. Samples were collected and stored in RNALater at <20°C until processed. RNA extraction was performed using the TRIzol® reagent (Invitrogen, USA) method following the manufacturer’s instructions and adding 5 µg of RNase-free glycogen as a carrier to the aqueous phase for low input samples. Concentration of all samples was assessed by Qubit RNA BR Assay kit (Thermo Fisher Scientific). Samples were subjected to Illumina’s TruSeq Stranded mRNA library preparation kit and sequenced on a NovaSeq 6000 (Illumina, 2 × 150 bp) for a 6Gb coverage.

### Gene repertoire evolutionary dynamics

In order to infer patterns of gene gain, duplication and loss that may have facilitated flatworm terrestrialization, we retrieved the proteomes (specifically the longest isoform files) of a set of species representing the main lineages of the phylum Platyhelminthes from MATEdb2 (Martínez-Redondo et al. 2024a; Vargas-Chávez et al. 2024), with particular emphasis on having a good representation of species of the order Tricladida, as this order comprises both freshwater and terrestrial species (Supplementary Table S1; Fig. 1). To polarise the comparative genomics results, we used a topology reflecting our current understanding of platyhelminth interrelationships (Benítez-Álvarez et al. 2025; Laumer et al. 2015). All species were subject to orthology inference in OMA v2.6 (Altenhoff et al. 2019) to infer hierarchical orthologous groups (HOGs), defined as set of genes that have descended from a single common ancestor within a taxonomic range of interest (Altenhoff et al. 2013). pyHam (Train et al. 2019) was used to infer gene repertoire evolutionary dynamics across the Platyhelminthes phylogeny based on the previously-inferred HOGs following the analytical strategy described in (Vargas-Chavez et al. 2024). In the case of the ancestor of terrestrial planarians, the duplicated and expanded HOGs were calculated in two ways. First, we used a cut-off of two copies in at least 20% in at least 20% of the species within Geoplanidae (‘unbalanced’ duplications and expansions hereafter) to ensure that expansions we considered were consistently present across multiple taxa, minimizing the impact of rare or lineage-specific duplications. Second, since we included four subfamilies within Geoplanidae, we recalculated duplicated and expanded genes considering only duplications or expansions that occurred in these four subfamilies, with the goal of recovering HOGs that duplicated or expanded at the MRCA of land planarians (Geoplanidae) with higher confidence (‘balanced’ duplications and expansions hereafter).

### Proteomics data and protein identification

Proteomic raw data of *O. nungara* and *S. mediterranea* was obtained from García-Vernet et al. (2025). Briefly, proteins were extracted by using TRIzol® reagent (Invitrogen, USA) following (Simões et al. 2013). The final protein pellets were resuspended in 25 µl of resuspension buffer (1%SDS and 8M Urea in Tris-HCl 1M, pH 8). Samples were digested and desalted for the LC-MS/MS analysis and analysed using a Orbitrap Fusion Lumos mass spectrometer (Thermo Fisher Scientific, San Jose, CA, USA) coupled to an EASY-nLC 1200 (Thermo Fisher Scientific (Proxeon), Odense, Denmark). The Proteome Discoverer software suite (v.2.5, Thermo Fisher Scientific) and the Mascot search engine (v2.6.2) were used for preprocessing the data and identifying the peptides. To perform the peptide identification, we used as reference databases the longest isoforms of *O. nungara* and *S. mediterranea*, as previously described for the gene repertoire dynamics analysis analyses. For constructing the final proteomic dataset, we only included proteins identified with high confidence levels and designated as the master protein within a protein group. We defined a protein group as a set of proteins containing at least two unique peptides. Detailed experimental and statistical methods can be found in García-Vernet et al. (2025).

### Differential gene expression involved in response to abiotic stress

With the goal of pinpointing the gene repertoire used in response to abiotic stress and explore its evolutionary origin, specimens of *Obama nungara* (terrestrial planarian, family Geoplanidae, n=30) and *Schmidtea mediterranea* (freshwater planarian, family Dugesiidae, n=36) were subjected to several abiotic conditions related to their terrestrial/freshwater ecological niches, respectively: exposure to visible and UV-B light, hyperoxia, hypoxia, osmotic stress and exposure to chemical cues (food and the proximity of a dead specimen of the same species). Specimens under visible light were kept under natural light (i.e., close to a window in the laboratory) for 15 minutes in an empty tray with no space to hide. For exposure to UV-B light, specimens were exposed under a UV-B lamp (302 nm) for 2 minutes. Animals were allowed to recover in darkness 24 hours in a dark chamber at a constant temperature of 16°C. Hyperoxia experiments were carried out by adding pure oxygen into a controlled chamber until reaching 38-44% oxygen for 20 minutes. Oxygen concentration was continuously monitored by an oxygen sensor located at the interior of the hyperoxia chambers (Presens OXY-1 ST Fiber). Hypoxia experiments were done in a HypoxyLab™ (Oxford Optronix) at 8% oxygen concentration for the aquatic species and 15% oxygen concentration for the terrestrial species for 20 minutes. For the osmoregulation experiment, *S. mediterranea* was immersed in sea water for 30 seconds to mimic osmotic stress, making sure that the exposure time was causing stress but not mortality and *O. nungara* was left inside a 9 cm petri dish covered in filter paper for 15 minutes to dry out, considering it as a low humidity condition for the animals. For the attraction to chemical cues associated with food, specimens were starved for a minimum of three days and then placed in 9 cm petri dishes. Fresh beef liver was used to attract the animals, whilst for the attraction to the scent of conspecific dead specimens, 100 ul of freshly smashed flesh was used in each of the species, and the experiments were stopped when a clear change of behaviour was observed in the specimens (in the food experiments, the specimens were touching the food and ejecting the pharynx to eat, and in the death experiments the specimens started to move faster going first towards and then opposite to the smashed flesh of a conspecific individual). Control samples were directly processed without any additional treatment. All samples were left in starvation for at least 24h before any experiment to reduce gut content. Three to five biological replicates per species and experiment were included. After the experiments, the specimens were flash frozen in liquid nitrogen and kept <-70°C until further processing. mRNA libraries were built and sequenced as indicated above for the terrestrial planarians (see number of specimens per experiment in Supplementary Table S7).

Quality assessment of the raw Illumina RNA-seq reads of *O. nungara* and *S. mediterranea* of all the experiments was performed using FastQC v0.11.9 (https://www.bioinformatics.babraham.ac.uk/projects/fastqc). Adapters and low-quality bases were trimmed using Trimmomatic v0.39 (MINLEN: 75, SLIDINGWINDOW: 4:15, LEADING: 10, TRAILING: 10, AVGQUAL: 30) (Bolger et al. 2014) and the trimmed RNA-seq reads were also assessed with FastQC before further analysis. For both species, a de novo reference transcriptome assembly generated with Trinity v2.11.0 (Haas et al. 2013) using as input a combination of the RNA-seq reads of the stress experiments and used further for differential gene expression analysis. The indexing and quantification of the transcripts was performed using Salmon v1.5.2 (Patro et al. 2017) for each sample. For each gene, the transcripts per million (TPM) value was calculated and a perl script (align_and_estimate_abundance.pl) included in the Trinity package was used to generate the counts and trimmed mean of M-values (TMM) scaling normalized expression matrices. Differential gene expression analysis was conducted using the perl script run_DE_analysis.pl also included in Trinity toolkit and the edgeR (Robinson et al. 2010) package. Finally, the Benjamini-Hochberg correction method (Benjamini and Hochberg 1995) was applied with the script analyze_diff_expr.pl with cut-offs of adjusted p-value at 0.001 (FDR) and a four-fold change.

### Evolutionary origin of the gene repertoire involved in response to abiotic stress

We assigned the differentially expressed genes in the stress experiments and the proteins detected by proteomics to HOGs. We then reconciled the HOGs with the Platyhelminthes species tree to infer the evolutionary origin of these genes following a phylostratigraphic reconstruction (i.e., in which node each HOG originated) based on pyHAM (Train et al. 2019) results.

### Detection of shifts in selection regimes in genes involved in response to abiotic stress

To detect selective pressure changes during transition to terrestrial habitat in HOGs including DEGs in *O. nungara* that arose earlier to the diversification of terrestrial planarians (i.e., in Tricladida, Continenticola, and Geoplanoidea), we used Pelican (Duchemin et al. 2023). This profile method detects directional selection changes across the amino acid sequences. We coded the non-terrestrial habitat as background and terrestrial as foreground character. Because the Pelican’s output includes a set of p-values for each site in the alignment, we used these site-level p-values to obtain gene-level prediction using the Gene-wise Truncated Fisher’s method (Duchemin 2023) considering the best k=10 p-values in each alignment. Additionally, we applied the adjustment method Benjamini & Hochberg (1995) (Benjamini and Hochberg 1995) (BH) based on the false discovery rate.

### Functional annotation and enrichment analyses

The putative function of genes identified as relevant in each of the analyses (i.e., gene repertoire evolution, differential gene and protein expression, etc) was explored through functional annotation and enrichment. For functional annotation, we used both homology-based methods eggNOG-mapper v2 (Cantalapiedra et al. 2021) and Natural Language Processing (NLP)-based ones (FANTASIA) (Martínez-Redondo et al. 2024b), since homology-based methods did not recover GO terms for a high percentage of the genes of interest (Supplementary Table S2). On average, 98.96% of sequences were annotated with FANTASIA at the gene ontology (GO) level, in contrast to a mean number of 67.16% sequences annotated with eggNOG-mapper at GO level and 48.68% at gene name level. Using these annotations, several enrichment analyses were performed related to the analyses of gene repertoire evolution, and differential gene and protein expression (Supplementary Table S3) using the Fisher test and the elim algorithm implemented in the topGO package (Rahnenfuhrer 2024). As a background for the enrichments, we used the HOGs generated previously to the event under exploration as we were interested in analysing what functions were enriched with respect to the genomic information arising prior to that event (i.e., this allows for a more fine-grain understanding of which functions changed during an internode). We analysed different sets of HOGs: 1) HOGs arising in different nodes, 2) HOGs including proteins detected by proteomics, 3) HOGs including DEGs, and 4) HOGs including DEGs detected by proteomics (Supplementary Table S3). Furthermore, in order to identify the enriched pathways including genes inferred as relevant for these analyses, we used Reactome (Milacic et al. 2024) with the annotations inferred by eggNOG-mapper v2. Finally, for a better understanding of the function of coding genes related to response to abiotic stress, we characterised the DEGs detected by proteomics in both species using the best hits obtained with BLAST (Camacho et al. 2009) these sequences against the no-redundant database (Available from: https://www.ncbi.nlm.nih.gov/gene/ last available update on May 1^st^ 2024).

## Results

### A burst of gene gain at the origin of Tricladida as a key driver of flatworm terrestrialization

Gene repertoire evolution inferred across Platyhelminthes including a representative data set of the phylum showed a high number of gained HOGs in the internodes leading to Tricladida, Continenticola, and Geoplanoidea (Fig. 2a). This pattern was confirmed at the proteomic level, since a similar pattern was found when we mapped the number of HOGs including translated proteins in *O. nungara* and *S. mediterranea* in the phylogeny (Fig. 2b; see below). On the contrary, the number of gained HOGs in the internode leading to the terrestrial planarians was low, indicating that gene gain was not particularly high at the onset of Geoplanidae and, thus, that the evolutionary appearance of new genes at the internode leading to terrestrial planarians was not a main driving force behind flatworm terrestrialization (Supplementary Table S4).

**Fig. 2.**
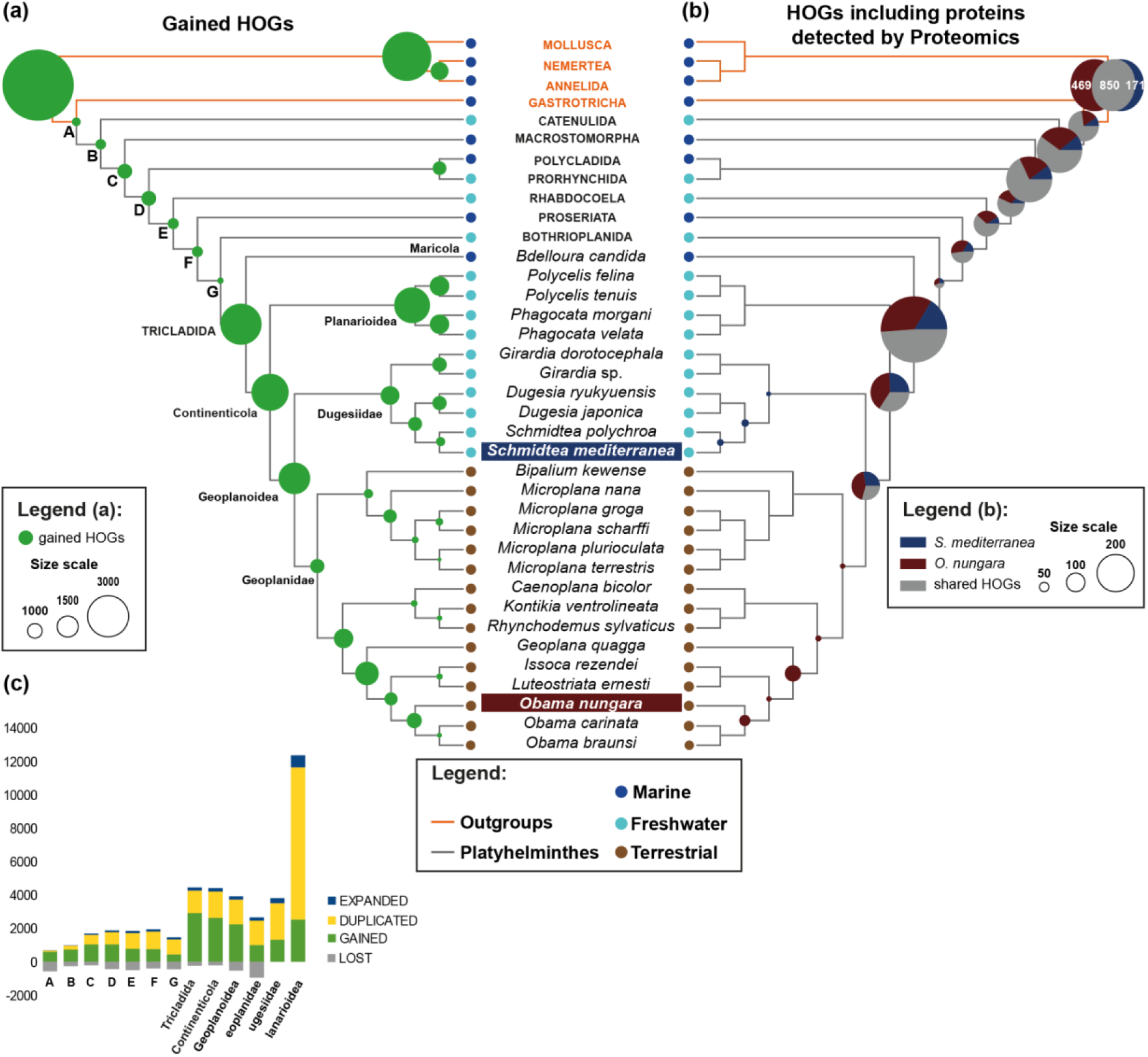
Gene repertoire evolution across Platyhelminthes. The outgroups are highlighted in orange and the members of Platyhelminthes are in black. The habitat is indicated in dark blue (marine), light blue (freshwater) and brown (terrestrial). a) Gained HOGs are represented in green circles on the nodes. The size is proportional to the number of gains in the lineage. Nodes belonging to Tricladida lineage are indicated with the taxonomic ranking. Other lineages are indicated with letters from A to G. b) Proportion of HOGs including proteins detected by proteomics. The pie charts represent the number of HOGs with assignment of sequences belonging to *Obama nungara* (dark brown), *Schmidtea mediterranea* (dark blue), or both species (grey). The size of the pie charts is proportional to the number of HOGs except for the first node where the information is shown in a Venn diagram and the number of HOGs is indicated. c) Stacked bar chart indicates the proportion of lost (grey), gained (green), duplicated (yellow), and expanded (blue) HOGs by node where the gene repertoire evolution was calculated. The notation of the nodes is the same as in the tree.

Regarding the putative function of the genes that arose in the branches leading to Tricladida, Continenticola, and Geoplanoidea, they were highly enriched in regulatory functions. Terms related to regulation of gene expression, transcription, and protein translation and maturation were enriched in the three lineages, as well as cellular transport and cellular cycle (Fig. 3a, Supplementary Material 1).

**Fig. 3.**
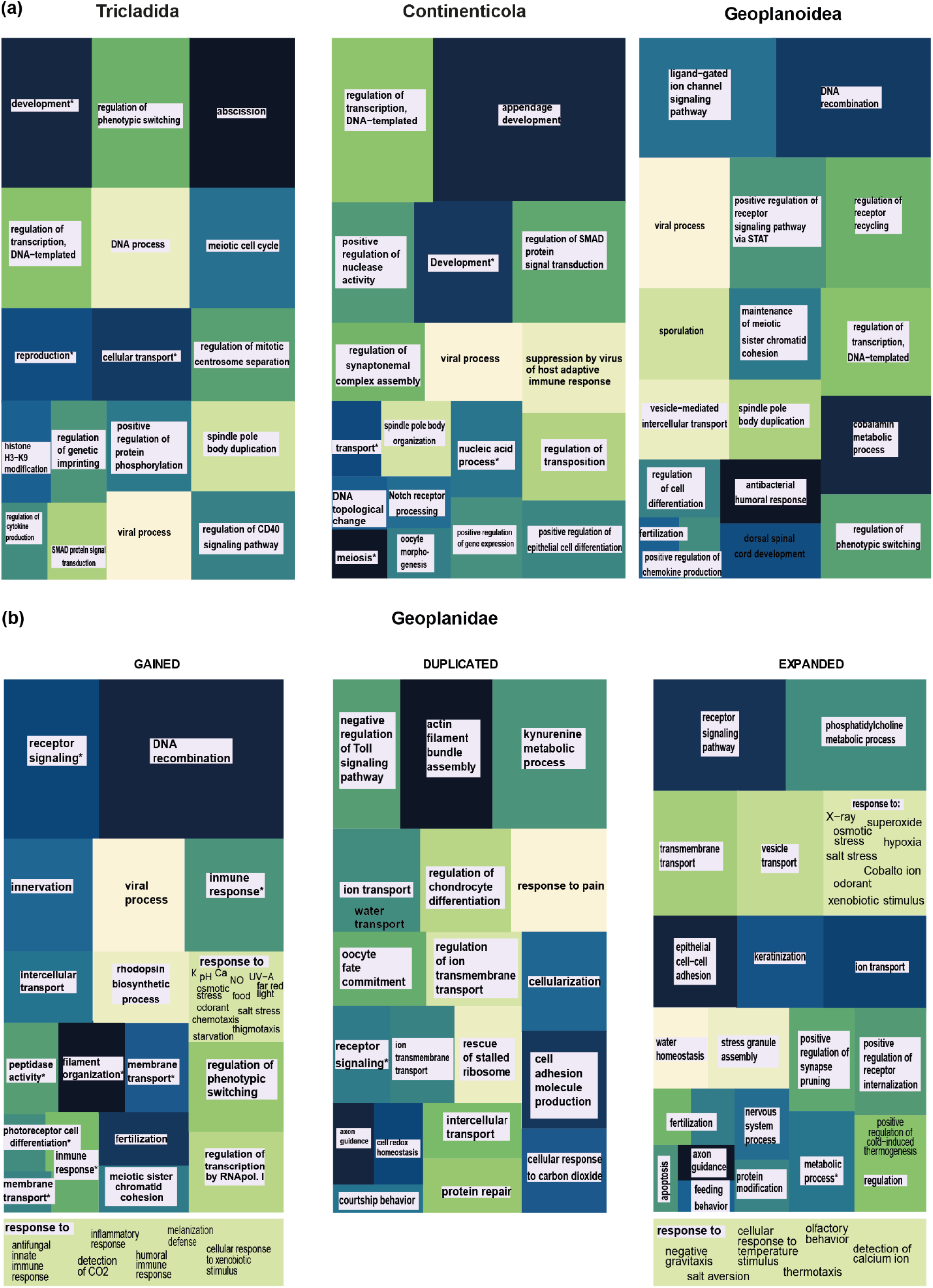
Simplified treemaps indicating the most enriched biological process in the gene repertoire across Tricladida. Only the most general GOs in the squares are indicated. a) Gained HOGs in Tricladida, Continenticola, and Geoplanoidea. b) Gained, duplicated and expanded HOGs in the ancestor of terrestrial planarians, the family Geoplanidae. The GO enrichment analyses were performed in duplicated and expanded HOGs, balanced between the major Geoplanidae lineages. Fractions representing “response to” functions are expanded under the figure.

### Ancient origin for most proteins in freshwater and terrestrial planarians

Translated genes were specifically investigated using proteomics data obtained through a bottom-up approach in *S. mediterranea* and *O. nungara*. A total of 2663 and 3704 proteins were confidently identified, respectively. For both species, HOGs containing these proteins originated before the splitting of terrestrial planarians and dugesiids (Supplementary Table S6), being specifically assigned to HOGs that arose during ancient diversification events: in the root branch, in the nodes leading to Catenulida and Macrostomorpha, as well as in Tricladida and Continenticola (Fig. 2b, Supplementary Table S6). This general pattern, similar to the one obtained in the gene repertoire evolution analyses, further supports the burst of gene gain observed within the gene repertoire dynamics at the Tricladida and Continenticola nodes. Different functions were enriched in proteins detected in both species, potentially indicating a different biological background regarding regulation, behaviour, cell differentiation, and cellular transport. Besides, most functions enriched in HOGs including proteins detected in both species (i.e., shared HOGs) were related to regulation, cellular transport, metabolic processes, and translation (Supplementary Material 2).

### Characterization of the gene repertoire in the ancestor of terrestrial planarians

Although the number of gains in the ancestor of terrestrial planarians was much lower compared to older nodes (Fig. 2a, 2c), enriched functions in gained HOGs during the splitting of this lineage were related to functions potentially related to adaptive response to life on land, such as processes related to nucleic acid stability (DNA recombination, meiotic sister chromatid cohesion, and regulation of transcription), regulation of cellular damage (regulation of neuron apoptotic processing, positive regulation of epithelial cell proliferation, positive regulation of oxidative stress-induced cell death), membrane transport, melanization defence response, negative regulation of photoreceptor cell differentiation, and response to several abiotic stimuli (ions concentration, pH, UV, osmotic stress, etc). Furthermore, functions related to these biological processes were also enriched in duplicated and expanded HOGs (Fig. 3b).

### Different genetic toolkit involved in the response to abiotic stress in freshwater and terrestrial planarians, yet arising at the same evolutionary time

We conducted a series of experiments aimed at identifying genes potentially involved in response to abiotic conditions related to freshwater or terrestrial environments through differential expression analysis, namely exposure to hypoxia and hyperoxia, chemosensory cues (food and dead specimens of the same species), UV and visible light, and osmotic stress (see Methods; summary statistics, density plots, volcano plots, expression heatmaps and matrices of expression values for all differentially expressed genes are available in the GitHub repository associated to this manuscript and as Supplementary Table S7). The number of differentially expressed genes (DEGs) per experiment is shown in Table 1.

**Table 1.** Assignment of differentially expressed genes (DEGs) to hierarchical orthogroups (HOGs) by experiment and species. UP: upregulated, Down: downregulated, Chemo_Death: chemoreception of dead conspecific animals, Chemo_Food: chemoreception of food, VL: visible light, UV24D: ultraviolet light with 24 hours of recovery in darkness.

| Experiment | *Obama nungara* |  |  | *Schmidtea mediterranea* |  |  |
| --- | --- | --- | --- | --- | --- | --- |
|  | DEGs | HOGs | DEGs assigned to HOGs | DEGs | HOGs | DEGs assigned to HOGs |
| Chemo_Death_UP | 140 | 49 | 59 | 69 | 26 | 29 |
| Chemo_Food_UP | 134 | 54 | 62 | 49 | 12 | 13 |
| Hyperoxia_UP | 881 | 617 | 641 | 75 | 3 | 3 |
| Hypoxia_UP | 846 | 536 | 562 | 99 | 5 | 6 |
| Osmoregulation_UP | 245 | 115 | 141 | 179 | 95 | 96 |
| UV24D_UP | 86 | 41 | 42 | 57 | 17 | 20 |
| VL_UP | 533 | 356 | 380 | 57 | 5 | 5 |
| **Unique UP** | **1677** | **891** | **995** | **468** | **140** | **148** |
| Chemo_Death_Down | 43 | 14 | 15 | 1861 | 172 | 194 |
| Chemo_Food_Down | 27 | 7 | 7 | 781 | 68 | 72 |
| Hyperoxia_Down | 565 | 315 | 330 | 9 | 0 | 0 |
| Hypoxia_Down | 226 | 97 | 98 | 42 | 5 | 6 |
| Osmoregulation_Down | 37 | 10 | 11 | 869 | 450 | 471 |
| UV24D_Down | 28 | 8 | 8 | 2672 | 208 | 236 |
| VL_Down | 74 | 34 | 41 | 2255 | 171 | 198 |
| **Unique Down** | **782** | **383** | **401** | **5580** | **756** | **861** |
| **Unique all experiments** | **2459** | **1270** | **1396** | **6045** | **895** | **1008** |
| **Exclusive HOGs *O. nungara*: 1270** |  |  |  |  |  |  |
| **Exclusive HOGs *S. mediterranea*: 738** |  |  |  |  |  |  |
| **Shared HOGs between species: 157** |  |  |  |  |  |  |

Differentially expressed genes were assigned to HOGs in order to understand their evolutionary origin in the planarian evolutionary chronicle. Although a high number of DEGs were not assigned to HOGs (due to DEGs either being species-specific or noncoding), the number of assigned genes allowed us to unveil important differences in the mechanisms of response to stress between freshwater and terrestrial planarians. A total of 1270 HOGs were identified including DEGs in *O. nungara* and 895 in *S. mediterranea*. From them, 891 included sequences under positive regulation and 383 under negative regulation in the terrestrial planaria. On the other hand, 148 and 861 were under positive and negative regulation, respectively, in the freshwater species.

The assignment of these genes to HOGs (Table 1, Supplementary Table S8) allowed us to infer their evolutionary origin through phylostratigraphy. Although most of the HOGs including DEGs arose in the same nodes for both species (those corresponding to Tricladida, Continenticola, and Geoplanoidea) (Fig. 4, Supplementary Table S8), the number of shared HOGs was very low (n=157, Table 1), indicating that most of the genes involved in response to stress in freshwater and terrestrial planarians had a different evolutionary origin, despite arising in the same branches of the planarian phylogeny. In agreement with this, enrichment analyses of HOGs including DEGs of each species showed different enriched functions, indicating different genetic mechanisms of response to stress in freshwater and terrestrial planarians (Supplementary Material 3).

**Fig. 4.**
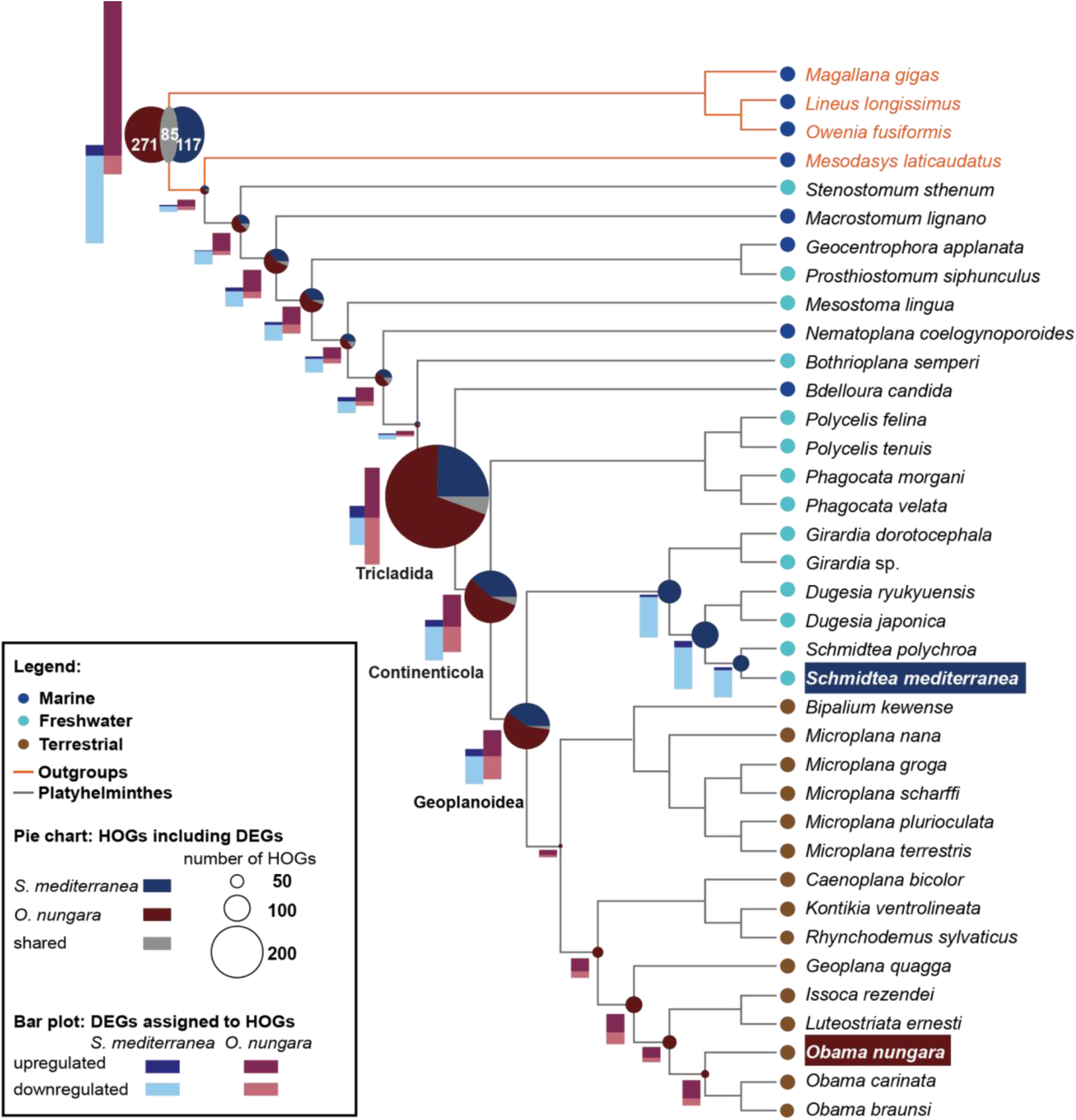
Evolutionary origin of differentially expressed genes (DEGs) assigned to HOGs. The bars represent the proportional number of upregulated and downregulated DEGs assigned to HOGs from *Schmidtea mediterranea* and *Obama nungara*. The pie charts represent the number of HOGs with the assignment of DEGs by species and the shared HOGs (HOGs with DEGs assigned from both species). The size of the pie charts is proportional to the number of HOGs except for the most basal node where the information is shown in a Venn diagram and the number of HOGs is indicated.

Remarkably, DEGs in both species also showed a different polarization in the direction of regulation. While *S. mediterranea* showed that most DEGs were down-regulatated, most DEGs in *O. nungara* were up-regulated (Fig. 4), indicating distinct evolutionary strategies in response to environmental stress. This suggests that *O. nungara* may have developed more active stress-response mechanisms, likely reflecting the greater challenges posed by terrestrial habitats, whereas *S. mediterranea* relies more on down-regulation pathways, which may be better suited to the relatively more stable conditions of freshwater environments.

In the case of *O. nungara,* most of DEGs assigned to HOGs were detected as upregulated in hyperoxia and hypoxia experiments. On the other hand, in the case of *S. mediterreanea* those DEGs were detected as downregulated in the osmoregulation experiment (Fig. 5a, and b).

**Fig. 5.**
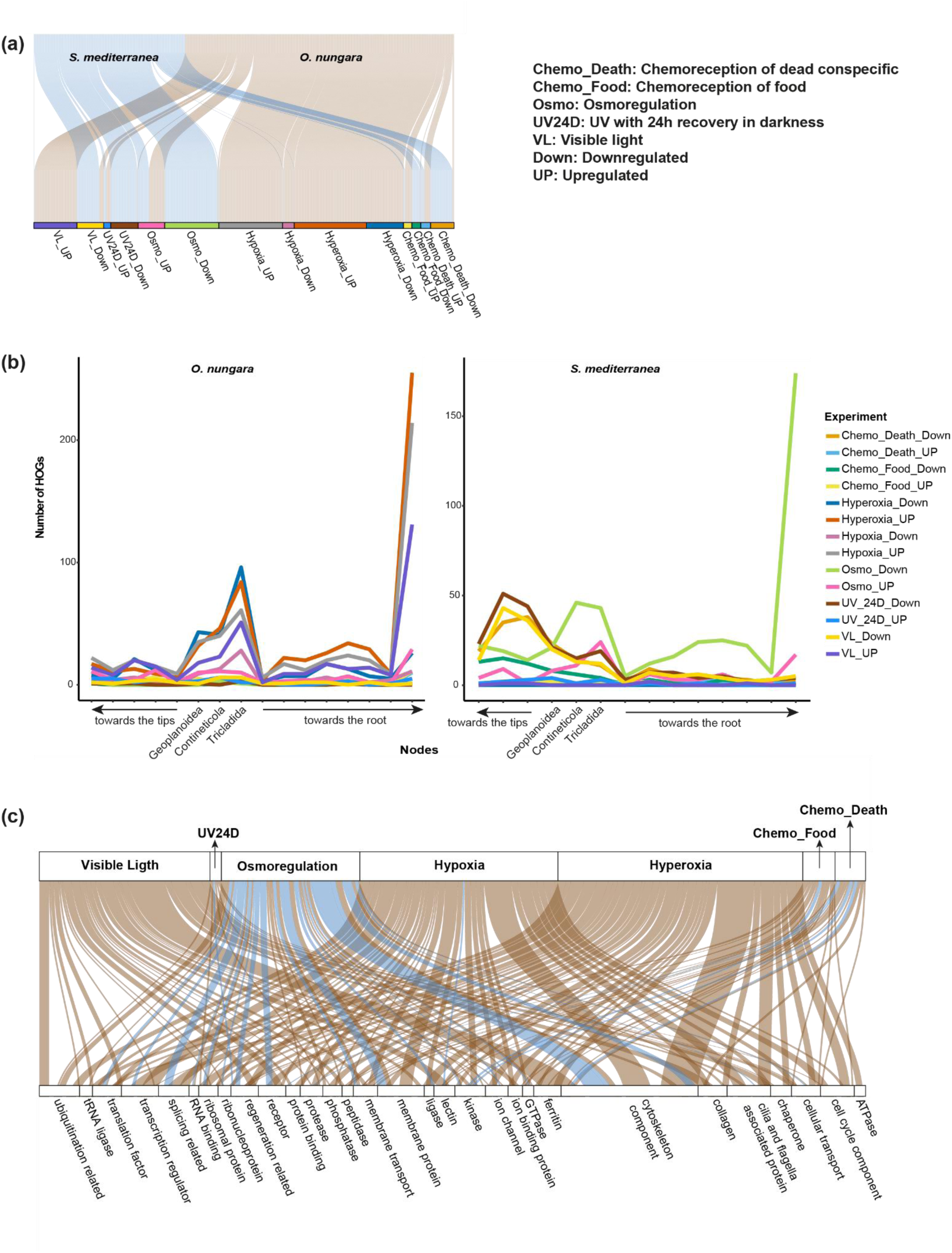
Functional characterization of DEGs. a) alluvial diagram representing the DEGs assigned to HOGs by experiments. b) graphs representing the number of HOGs with DEG assigned by node by experiment for *O. nungara* and *S. mediterranea* independently c) alluvial diagram representing the most frequent functionalities in which are implicated the BLAST hits of DEGs in stress experiments detected also by proteomics (DEGs-prot) in both species *O. nungara* (brown) and *S. mediterranea* (blue).

Regarding the common DEG-containing HOGs, no shared HOGs were found when the DEGs in the same experiment and in the same direction of regulation were compared for both species. However, DEGs detected in different experiments and with a different direction of regulation between species (i.e., upregulated in one species and downregulated in the other) were included in the same HOGs. Most of these genes were downregulated in *S. mediterranea* when exposed to osmotic stress, while upregulated in *O. nungara* when exposed to hypoxia, hyperoxia, visible light or osmotic stress (Table 1; Supplementary Material 4, Supplementary Table S9). This pattern suggests that these genes are involved in fundamental stress-response pathways but have diverged in their regulation between terrestrial and freshwater planarians, reflecting the different environmental pressures.

To gain deeper insights into the functional implications of differential gene expression, we examined whether the identified DEGs were translated into proteins. For that, we identified the DEGs that were detected by proteomics (DEGs-prot hereafter). Among the DEGs assigned to HOGs, 357 out of 1396 were detected through proteomics in *O. nungara*, and 129 out of 1008 in *S. mediterranea* (Table 2). Phylostratigraphy analysis, which identified the evolutionary origin of each DEG-containing HOG, revealed that the majority originated in Tricladida, Continenticola, and Geoplanoidea (Supplementary Table S10). These findings validate the expression of these genes and support their involvement in the response to stressful environmental conditions.

**Table 2.** Assignment of differentially expressed genes (DEGs) and proteins to HOGs. DEG-prot: DEGs that were detected by proteomics. Shared HOGs: HOGs including genes from *O. nungara* and *S. mediterranea*.

|  | Transcriptomics (DEGs) | Proteomics | DEGs-prot |
| --- | --- | --- | --- |
| **Assigned genes to HOGs** |  |  |  |
| *O. nungara* | 1396 | 3510 | 357 |
| *S. mediterranea* | 1008 | 2605 | 129 |
| **Identified HOGs** |  |  |  |
| *O. nungara* | 1270 | 1228 | 173 |
| *S. mediterranea* | 895 | 540 | 42 |
| Shared HOGs | 157 | 1768 | 35 |

The functional characterization of these genes through GO enrichment, pathways identification, and functional annotation with BLAST showed a differential functional background in these genes between the freshwater and the terrestrial species, with a wider diversity of functions identified in *O. nungara* (Fig. 5c, Supplementary Fig. 1, Supplementary Material S5, Supplementary Table S11). Proteins involved in response to stress in *O. nungara* were enriched in functions related to development, regulation, cytoskeleton organisation, gene expression and transport (Supplementary Fig. 1, Fig. 5c, Supplementary Table S11). On the other hand, although *S. mediterranea* share many general functions (Supplementary Fig. 1), the number of genes and the enriched functions are different to the terrestrial species (Supplementary Fig. 1). Regarding the proteins identified by BLAST hits, the most frequent functions detected in both species are related to cytoskeleton components and membrane proteins (Fig. 5c).

Regarding the gene repertoire involved in response to the different abiotic stressors tested in the experiments, both species showed contrasting patterns. In *O. nungara*, most DEGs-prot were detected when specimens were exposed to hyperoxia, hypoxia and visible light experiments. On the other hand, in *S. mediterranea* most DEGs-prot were detected when individuals were exposed to osmoregulatory stress (Fig. 5c). The evolutionary origin of most DEG-containing HOGs in response to hypoxia, hyperoxia and exposure to visible light in *O. nungara*, and in response to osmotic stress in *S. mediterranea*, coincide with the burst of gene gain at Tricladida, and then in Continenticola and Geoplanoidea (Supplementary Table S10), in line with the burst of gene gain detected genome-wide and discussed above. This suggests that the bursts of gene gain at these phylogenetic nodes contributed significantly to the development of adaptive mechanisms in flatworms, allowing them to cope with specific environmental stressors encountered in their respective habitats.

### Half of DEGs gained before the diversification of terrestrial planarians are under directional selection

To detect changes in selection pressures potentially related to habitat shift, we selected DEG-containing HOGs with a balanced representation of both terrestrial and non-terrestrial planarians (i.e., including at least 20% of the species in each group), resulting in a dataset comprising 444 HOGs (see Methods). A statistically significant shift in directional selection in terrestrial planarians was detected in more than 50% of these HOGs (Fig. 6, Supplementary Table S12-13).

**Fig. 6.**
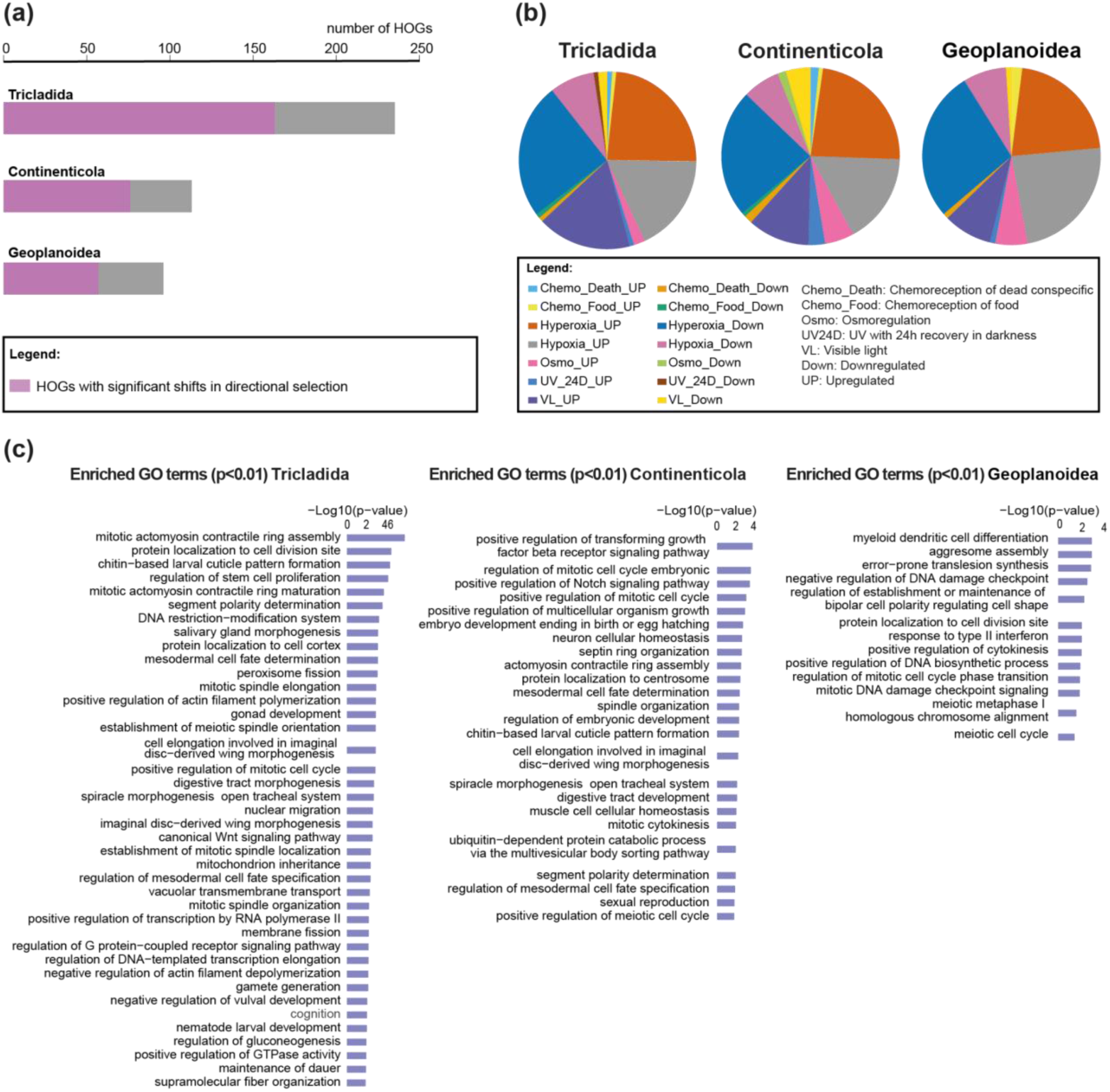
DEGs gained in Tricladida, Continenticola, and Geoplanoidea under directional selection. a) Proportion of HOGs under directional selection in the three lineages. The full bar represents the total number of DEG-containing HOGs with a balanced representation of both terrestrial and non-terrestrial planarians (see Methods). In pink, the proportion of HOGs under directional selection. b) Proportion of HOGs under directional selection by experiment and regulation. c) Enriched GO terms (p<0.01) of DEGs under significant shifts of selection in the experiments of hypoxia, hyperoxia and osmoregulation. The longest bar represents the lowest p-values.

Enriched functions of DEGs included in HOGs under selection arising in Tricladida are principally related to morphogenesis, cellular organisation, regulation of DNA template transcription, and transmembrane transport. Likewise, DEGs arising in Continenticola showed enriched functions related to morphogenesis also, cellular cycle regulation, DNA damage response, cellular transport, response to stimulus, and regulation of translation. Finally, DEGs arising in Geoplanoidea are enriched in functions related to DNA protection such as regulation of transcription and DNA repair, and cell cycle regulation (Supplementary Material 6). In summary, most of DEGs detected in *O. nungara*, which arise in Tricladida, Continenticola, and Geoplanoidea showed shift selection signal and are enriched in functions related to morphogenesis, cellular organisation, cell cycle regulation, DNA protection, and transmembrane transport.

Notably, most genes displaying selection shift signals are associated with responses to hypoxia, hyperoxia, and visible light (Fig. 6b, Supplementary Table S12). In Tricladida, the newly emerged genes are enriched in functions such as amino acid transmembrane transport, barbed-end actin filament capping, cell cycle DNA replication, macromolecule localization, reproductive processes in multicellular organisms, embryo development, filamentous growth, microtubule-organizing center organization, cell cycle regulation, and transcriptional control. For Continenticola, the key enriched functions include the regulation of translation, vacuolar transport, cellular stress responses, exocrine system development, positive regulation of cell cycle processes, microtubule organization, protein stabilization, responses to mechanical stimuli, and reproductive functions (e.g., embryonic cleavage and sexual reproduction). Finally, in Geoplanoidea, the genes that arose are related to diapause initiation, karyosome formation, host transcription modulation by symbionts, negative regulation of biosynthetic processes, positive regulation of intracellular signaling and sister chromatid cohesion, primary cell septum biogenesis, response to various stimuli (including interferon-gamma), protein phosphorylation, RNA processing, and vesicle coating (Fig. 6c, Supplementary Material 7).

## Discussion

### Gene gain and genomic exaptation as key drivers of flatworm terrestrialization

In this study, we unveiled a burst of gene gain detected much earlier than the origin of terrestrial planarians (i.e., in the internodes leading to Tricladida, Continenticola, and Geoplanoidea). The newly gained and duplicated genes were significantly enriched in functions related to gene regulation, transcription, and protein maturation—processes essential for coordinating complex developmental and adaptive pathways. This may indicate that the observed burst of gene gain may have provided the raw material for adaptive innovations that were key to the successful colonisation of land, as tested and discussed below. In this study, we detected a burst of gene gain well before the emergence of terrestrial planarians (i.e., in the internodes leading to Tricladida, Continenticola, and Geoplanoidea). These newly gained and duplicated genes were enriched in functions related to gene regulation, transcription, and protein maturation—key processes for coordinating complex developmental and adaptive pathways. Such a genomic expansion could have provided the raw material needed for the adaptive innovations that ultimately enabled planarians to colonize land.

Although whole-genome duplication (WGD) is a strong candidate for driving this burst of gene gain, other mechanisms—such as chromosomal rearrangements, tandem and segmental duplications, or even de novo gene birth—may also have contributed to the observed expansions. Recent work in clitelates (Lewin et al. 2024; Vargas-Chavez et al. 2024) demonstrates how chromosomal rearrangements can radically reshape genomes, creating novel genes and modifying existing ones to meet adaptive demands. In planarians, the high levels of genomic rearrangements reported within the genus *Schmidtea* and in comparisons with *Macrostomum* (Ivanković et al. 2024) suggest that similar structural changes might also be at play, reshuffling regulatory regions, altering gene expression, or producing entirely new genetic material through fusion events. The burst of novel HOGs and the expansion of regulatory genes in our dataset lend strong support to WGD as the most likely primary mechanism driving gene gains in Tricladida and related lineages. Still, chromosomal rearrangements cannot be ruled out and merit further study, given their potential to complement WGD by fine-tuning genomic architectures for adaptive purposes—an effect also noted in annelids (Vargas-Chavez et al. 2024). Ultimately, the absence of high-quality reference genomes for terrestrial planarians constrains our ability to determine the relative contributions of these mechanisms, underscoring the need for further research in this area.

Gene gain is a crucial driver of adaptation in animals, enabling them to survive and thrive in new environments by introducing novel functions and expanding genetic diversity. In various animal species, gene duplications and de novo gene gains have facilitated adaptations to environmental challenges. For example, in invasive species such as the sea lamprey, genetic bottlenecks followed by the emergence of outlier genes associated with key life history traits demonstrate how gene gain can contribute to rapid adaptation in novel habitats (Yin et al. 2021). Similarly, in parasitic trematodes like *Atriophallophorus winterbourni*, gene duplications and novel genes have been shown to be essential for the evolution of parasitic mechanisms, showcasing the adaptive potential of new genes in a host-specific environment (Zajac et al. 2021).

Similarly to our findings, (Aristide and Fernández 2023) showed that multiple independent shifts from marine to freshwater and terrestrial environments in mollusks were facilitated by parallel expansions of ancient gene families. These expansions were predominantly involved in critical processes such as metabolism, osmoregulation, and defense mechanisms, showing the relevance of pre-existing genetic toolkits in adapting to new, non-marine environments. (Aristide and Fernández 2023) also identified convergent evolution at the orthogroup level, revealing that while parallel expansions were key to the molluscan transitions towards freshwater and land, lineage-specific gene gains further contributed to the development of unique adaptations in different lineages. Their study also showed that while lineage-specific gene gains played a role in providing unique adaptations, much of the adaptive toolkit stemmed from pre-existing genomic features. In contrast, our results suggest that for terrestrial planarians, older gene gains were especially important, with little evidence of more recent, lineage-specific acquisitions contributing to land colonization. This highlights how planarians may have relied more heavily on repurposing ancient genetic innovations, underscoring the central role of pre-existing gene repertoires in meeting the challenges of a terrestrial lifestyle.

Overall, these findings align with broader patterns seen in animal evolution, where gene gain has been repeatedly linked to adaptive innovations. By generating genetic variation and providing the raw material for selection, gene gain enables organisms to develop specialised traits, such as enhanced stress tolerance, new metabolic pathways, or unique behavioural adaptations (Hull et al. 2017; Balart-García et al. 2023). In terrestrial planarians, this process likely played a central role in their evolutionary success, driving the genomic changes necessary for coping with the challenges of terrestrial life.

Moreover, our results showed that ca. half of DEGs that arose prior to the origin and diversification of terrestrial planarians (i.e., in Tricladida, Continenticola, and Geoplanoidea) were under directional selection. This suggests that gene gain not only contributed to their initial adaptation to land but also continued to be refined through evolutionary pressures, enhancing their fitness in terrestrial environments. The evidence of directional selection acting on these gained genes indicates their critical role in stress-response mechanisms and other adaptive traits needed for survival on land.

### Divergent stress response mechanisms in terrestrial and freshwater planarians

Our comparative transcriptomic analysis between the terrestrial *O. nungara* and the freshwater *S. mediterranea* revealed striking differences in how these species respond to environmental stressors. In general, *O. nungara* seems to possess a broad genetic repertoire that enables it to respond effectively to various environmental stressors, particularly changes in oxygen concentrations (hypoxia and hyperoxia) and exposure to visible light. These stressors are among the most significant challenges in the adaptation to terrestrial life, where fluctuations in oxygen availability and increased exposure to light and UV radiation require robust protective mechanisms. The extensive set of DEGs-prot in *O. nungara* under these conditions suggests that this species has repurposed existing genes to develop new functions suited to terrestrial environments. On the other hand, *S. mediterranea* displays a stress response machinery predominantly focused on osmoregulatory functions, which is crucial for survival in freshwater habitats. The evolutionary origin of most DEG-containing HOGs responding to hypoxia, hyperoxia, and visible light in *O. nungara*, and to osmotic stress in *S. mediterranea*, coincides with bursts of gene gain at the Tricladida, Continenticola, and Geoplanoidea nodes (Fig. 5b). This suggests that these periods of genomic expansion provided a reservoir of genetic material that could be differentially exapted or co-opted in each lineage to meet specific environmental challenges. In *O. nungara*, genes initially involved in basic cellular processes may have been repurposed to develop enhanced responses to oxygen fluctuations and light exposure, conferring a selective advantage in terrestrial ecosystems. Similarly, in *S. mediterranea*, existing genes may have been co-opted to improve osmoregulatory functions, essential for maintaining homeostasis in freshwater environments. These lineage-specific adaptations highlight the role of genomic exaptation and co-option in the evolutionary history of flatworms. By differentially utilizing the genetic material acquired during key evolutionary events, each species developed specialized stress response systems tailored to their habitats. The ability of *O. nungara* to respond to terrestrial stressors and of *S. mediterranea* to manage osmotic stress exemplifies how genomic plasticity facilitates adaptation to diverse ecological niches.

Not only were DEGs largely non-overlapping between the two species, but they also showed opposite patterns of regulation. While most DEGs in *S. mediterranea* were down-regulated in response to stress, *O. nungara* exhibited a predominance of up-regulated DEGs. This pattern suggests that these genes are involved in fundamental stress-response pathways but have diverged in their regulation between terrestrial and freshwater planarians, reflecting the differing environmental pressures. *S. mediterranea*, inhabiting a more stable freshwater environment, may suppress these genes to conserve energy under stress, whereas *O. nungara*, facing more variable terrestrial conditions, may potentially activate these pathways to trigger a more active response to daily environmental fluctuations. Interestingly, common DEG-containing HOGs in both species were found to be upregulated in *O. nungara* and downregulated in *S. mediterranea* in response to different stressors, such as hypoxia, hyperoxia, and osmotic stress. This pattern implies that these genes are involved in core stress-response pathways but have diverged in their regulatory roles, reflecting the distinct evolutionary pressures faced by freshwater and terrestrial environments. The enrichment of different functional categories in the DEGs of each species further supports this divergence, with *O. nungara* showing an enrichment of functions related to active cellular processes such as transport and metabolism, while *S. mediterranea* exhibits functions tied to homeostasis and stability (Supplementary Material 4).

Such divergent adaptive responses to environmental pressures have been observed across a variety of taxa. For example, (Zhang et al. 2024) found that GR-RBPa proteins in plants exhibited differential expression under the same stress conditions across species, indicating that regulatory roles are shaped by specific environmental factors. In the same vein, (Rahi et al. 2019) discovered that freshwater prawns from the genus *Macrobrachium* had different expression patterns of candidate genes for freshwater adaptation, which likely evolved due to the unique pressures of freshwater environments. These studies align with the divergent gene expression patterns observed between terrestrial and freshwater planarians, where distinct stressors in aquatic versus terrestrial habitats may have driven these differences. Additionally, Lacasse and Aubin-Horth (2019) demonstrated how neuropeptide receptor expression diverged between stickleback populations, revealing adaptations to different ecological niches. Similarly, in the planarians, genes involved in stress-response pathways likely underwent functional divergence as the species adapted to their specific environments. This divergence is supported by the observation that genes within the same HOGs were upregulated in *O. nungara* in response to hypoxia and visible light stress, but downregulated in *S. mediterranea* when exposed to osmotic stress. The regulation of these shared HOGs suggests they play fundamental roles in stress response, but their divergent expression patterns reflect adaptations to different ecological pressures. Overall, these studies underscore that the evolutionary divergence in stress-response mechanisms between terrestrial and freshwater planarians reflects broader trends observed in other taxa. The distinct upregulation and downregulation strategies between *O. nungara* and *S. mediterranea* highlight how habitat-specific pressures—such as oxygen variability, light exposure, and osmotic stress—have likely driven the evolution of different regulatory responses. These differences not only reveal the unique challenges faced by these species in their respective environments but also provide insight into the broader patterns of evolutionary adaptation across species.

### Key genes in stress response and adaptation to terrestrial life in planarians

To further dissect the functions related to stress response in freshwater and terrestrial planarians we analyzed the functional annotation of DEGs detected by proteomics (DEGs-prot; Supplementary Fig. 1 and Fig. 5c). The functional annotations inferred by BLAST against the nucleotide database unveiled the importance of cellular process and regeneration in response to stress in these animals. Components of the cytoskeleton were the most frequent elements in both species. Among them, one of the most repeated in both species were the Ankyrin domains. Ankyrins are a family of adaptor proteins that link integral membrane proteins with the submembranous actin/β-spectrin cytoskeleton. Ankyrins are able to interact with a high variety of proteins, organising and stabilising protein networks (Cunha and Mohler 2009). The *TRPA1,* an ion channel key in the transduction of nociceptive signals in *S. mediterranea* has 14 N-terminal ankyrin repeats. TRPA1 in this freshwater species is activated in response to stressful conditions involving also the production of H_2_O_2_/ROS in response to noxious heat, contributing to channel activation to mediate defensive responses (Arenas et al. 2017). Additionally, Ankyrins are related to the regeneration and cellular turnover in planarians and are part of the group of orthologous genes conserved across whole body regenerative animals (Chereddy and Makino 2024).

The differential expression of the basal body component CEP135 in response to stress conditions in both species may highlight the key role of epidermis in stimulus perception and protection in these organisms. CEP135 is an essential protein for the basal body assembly in planarians (Azimzadeh and Basquin 2016). Actin components are also important elements of cytoskeleton. In planarians, Actin filaments are actively involved in regeneration and cellular turnover (Pascolini et al. 1993; Liu et al. 2023). Additionally, other cytoskeleton components such as microtubules associated with proteins are fundamental in the regeneration process (Gambino et al. 2022). On the other hand, dynein, also found differentially expressed in both species, is fundamental for cilia motility, an important function related to locomotion as well as excretory and reproductive systems in planarians (Lesko and Rouhana 2020). Another protein detected here with high frequency in both species is collagen. Collagen is one of the most abundant structural proteins in animals with 28 collagen types described in vertebrates (Shoulders and Raines 2009). In planarians, collagen proteins are an important component of muscular fibres and are the major structural components of the extracellular matrix (Cote et al. 2019) with an important role in regulating proliferation of neoblast during the regeneration process (Chan et al. 2021).

The high number of DEGs-prot related to cytoskeleton components, cell cycle, and regeneration process per se may indicate that the regeneration machinery is key in the response to stress in planarians. Continuous cellular turnover allows the replacement of damaged cells, and this can be crucial in the maintenance of tissue homeostasis through the unique mechanism described in planarians. This process has been extensively described in freshwater planarians belonging to the family Dugesiidae, being *S. mediterranea* the model species used in most studies, since the regenerative capability is highly variable, from whole body regeneration in dugesiids to the total absence of regeneration in several marine planarians (Ivankovic et al. 2019; Vila-Farré et al. 2023). Although only few regeneration studies have included terrestrial planarians, the results until moment indicate that terrestrial planarians, the sister group of Dugesiidae, can also show high regenerative capabilities (Milanese and Carbayo 2024), as is the case of *Luteostriata abundans* and others invasive lands planarians as *Bipalium kewense* and *Dolichoplana striata,* whose high regeneration capacity even allows them to reproduce asexually (Boll et al. (2023) and references therein; Milanese and Carbayo 2024). In planarians, the differentiated cells do not divide mitotically, and the neoblasts are the only dividing cells (Morita and Best 1974; Newmark and Sánchez Alvarado 2000). Thus, the cellular turnover driven by neoblasts differentiation is the only way to maintain tissue homeostasis, cell number and body size in these animals, allowing the regulation of body growth depending on environmental conditions (González-Estévez and Saló 2010; González-Estévez et al. 2012; Felix et al. 2019). The continuous cell turnover implicates the periodic elimination of selected differentiated cells and their replacement by the differentiated descendant of adult stem cells that become functional (Pellettieri and Sánchez Alvarado 2007). The differentiation process implicates a series of cellular changes under strong genetic regulation that guarantees the proper maintenance of the planarian body plan. Our findings suggest that the machinery governing cellular turnover may play a critical role in responding to environmental stress and adapting to new habitats in planarians. The gene repertoire associated with this response could also be linked to the evolution of regeneration capabilities across Platyhelminthes. Specifically, the burst of gene gain identified in Tricladida may be a key factor in the evolutionary success of this group, potentially driving the expansion of regenerative abilities observed in triclads. This hypothesis opens avenues for future research to further explore the interplay between gene repertoire evolution, cellular turnover, and regenerative success in this remarkable lineage.

We also found a high number of lectins in both species. Lectins have been linked to terrestrialization in mollusks due to the expansions of lectin gene families in terrestrial mollusks and their key role in innate immunity (Aristide and Fernández 2023). Additionally, lectin domains have been predicted as toxic in *O. nungara* in a recent study, with a potential protection function in terrestrial planarians (García-Vernet et al. 2025). The existence of specific families of lectins described in free-living flatworms (Shagin et al. 2002) and our results point out the important role that these proteins can play in the evolution of Platyhelminthes, especially in complex processes of diversification such as the conquest of land.

Another protein found differentially expressed in both species and detected by proteomics was ferritin. This protein is the main intracellular iron storage in most eukaryotic cells (Plays et al. 2021), with an important cytoprotective functions against oxidative damage (Arosio et al. 2015) and the innate immunity by similar mechanisms in both vertebrates and invertebrates as well as the regulation of apoptosis process (Ong et al. 2006). Additionally, ferritin has been described to be involved in response to anoxia in the interstitial mollusk *Littorina littorea* (Larade and Storey 2004). We identified ferritin differentially expressed in response to hyperoxia and others experiments in *O. nungara* and in osmoregulation and chemoreception in *S. mediterranea* (Supplementary Table S11). In planarians, ferritin is also related to reproduction. In sexual freshwater planarians, ferritin in association with the Wnt pathway is fundamental for yolk-egg production (Vila-Farré et al. 2023). Due to the importance of ferritin in functions avoiding oxidative stress and immunity, this protein differentially expressed in almost all experiments in *O. nungara* could be related to the response to stress in terrestrial planarians.

Several DEGs-prot were detected only in *O. nungara.* Among them, we can highlight the Cathepsin L, a protein expressed in the intestinal system and probably related to food digestion, and also described as a biomarker in response to heavy metals contamination in *Dugesia japonica* (Ma et al. 2019). Here we found this protein upregulated in the two types of chemoreception experiments as well as in osmoregulation in a terrestrial planaria. Another gene detected only in *O. nungara* was the E3 ubiquitin-protein ligase HUWE1. In general, the HUWE1 ligase is a key regulator of DNA damage response, cell cycle control, apoptosis, autophagy and inflammation (Gong et al. 2020). In *S. mediterranea*, it has been detected to be highly expressed in neoblasts with an important role in regeneration and stem cell regulation (Henderson et al. 2015). However, here we detected this gene upregulated in almost all experiments (chemoreception food, hyperoxia, hypoxia, osmoregulation, and visible light) only in *O. nungara*.

All in all, our study underscores the pivotal role of the regeneration machinery and specific gene repertoires in the stress responses of both freshwater and terrestrial planarians. The abundance of differentially expressed genes related to cytoskeletal components, cell cycle regulation, and regeneration indicates that continuous cellular turnover is crucial for adapting to environmental stress and new habitats. Proteins like lectins and ferritin play significant roles in innate immunity and protection against oxidative damage, particularly in terrestrial species like *O. nungara*. The unique expression of proteins such as Cathepsin L and HUWE1 in *O. nungara* suggests the development of specialized molecular pathways to cope with land-specific challenges. These findings imply that the bursts of gene gain in the Tricladida lineage may have facilitated the expansion of regenerative capabilities and stress response mechanisms, contributing to the evolutionary success of planarians. Future research exploring these molecular adaptations will deepen our understanding of planarian evolution, their remarkable regenerative abilities, and their relationship with ecological adaptation.

### Final remarks

This study unveils the genomic mechanisms underlying planarian terrestrialization, offering significant insights into the evolutionary processes that enabled this transition. We identified a burst of gene gain occurring earlier in flatworm evolutionary history—prior to the emergence of land planarians. This genomic innovation, enriched in regulatory and stress-response functions, likely provided a foundational genetic toolkit that was later co-opted through genomic exaptation to facilitate adaptation to terrestrial environments. Interestingly, we did not observe significant gene gains at the branch leading directly to terrestrial planarians. Instead, our findings suggest that the pre-existing expanded gene repertoire was repurposed to meet the new challenges of life on land. This illustrates how genomic exaptation can be more important than gene gains occurring precisely at the node where phenotypic changes are observed. The ability to utilize and adapt existing genes underscores the significance of gene regulation and functional divergence in evolutionary adaptation. Our study also highlights divergent stress-response mechanisms between the terrestrial *Obama nungara* and the freshwater *Schmidtea mediterranea*, indicating that these species have evolved distinct strategies to cope with environmental stressors. In *O. nungara*, the upregulation of stress-related genes suggests a dynamic response to terrestrial challenges such as fluctuating oxygen levels and increased exposure to visible light—key obstacles in adapting to land environments. Conversely, *S. mediterranea* exhibits stress-response mechanisms primarily focused on osmoregulation, which is advantageous in freshwater habitats where maintaining osmotic balance is essential. This divergence emphasizes the crucial role of gene regulation and exaptation in evolutionary adaptation.

Overall, this study provides the first comprehensive genome-wide evidence elucidating the mechanisms driving flatworm terrestrialization. Our findings not only expand our understanding of land colonization in flatworms but also align with broader evolutionary patterns observed across other taxa. Specifically, we reveal that multiple evolutionary strategies—including early gene gains, expansion of conserved gene families, directional selection, and the repurposing of existing genes—have been instrumental in overcoming the environmental challenges of transitioning from aquatic to terrestrial habitats. Integrating our new insights from planarians with previous phylum-wide, genome-wide studies in other groups such as mollusks underscores the complexity and diversity of the genomic processes underlying terrestrialization across animal lineages. The fact that significant genomic innovations occurred prior to the phenotypic transition highlights the importance of pre-existing genetic diversity and the role of exaptation in evolutionary innovation. Collectively, this research contributes to a deeper understanding of the genomic factors that enable large-scale evolutionary convergence during habitat transitions, elucidating how animals have repeatedly adapted to life on land throughout evolution.

## Available Information

A repository containing the scripts and files (inputs, intermediates, and outputs) needed to reproduce the analyses as well as several supplementary files for a better understanding of results is available here: https://github.com/MetazoaPhylogenomicsLab/Benitez-Alvarez_et_al_2025_Gene_repertoire_evolution_land_planarians

## Acknowledgements

We thank Adrian Altenhoff for guidance and discussions about pyHAM. Gemma I. Martínez-Redondo kindly helped us collect *Microplana* specimens, and shared scripts to improve the GO enrichment treeplots. RG-V acknowledges the support of the Margarita Salas grant funded by the Spanish Ministry of Universities and the European Union Next Generation EU/PRTR. RF acknowledges support from the following sources of funding: Ramón y Cajal fellowship (grant agreement no. RYC2017-22492 funded by MCIN/AEI /10.13039/501100011033 and ESF ‘Investing in your future’), the European Research Council (this project has received funding from the European Research Council (ERC) under the European Union’s Horizon 2020 research and innovation programme (grant agreement no. 948281)), the OSCARS project (funding from the European Commission’s Horizon Europe Research and Innovation programme under grant agreement no. 101129751) and the Secretaria d’Universitats i Recerca del Departament d’Economia i Coneixement de la Generalitat de Catalunya (AGAUR 2021-SGR00420). FÁF-Á was supported by a Beatriu de Pinós fellowship from Secretaria d’Universitats i Recerca del Departament de Recerca i Universitats of the Generalitat de Catalunya (Ref. BP 2021 00035) and a Ramón y Cajal fellowship (Ref. RYC2023-043494-I) funded by MCIN/AEI /10.13039/501100011033 and FSE+. This research was supported by the Spanish government through the ‘Severo Ochoa Centre of Excellence’ accreditation (CEX2019-000928-S). We also thank Centro de Supercomputación de Galicia and CSIC for access to computer resources (CESGA and DRAGO respectively).

