## Supplementary Figure S1 for "Genomic exaptation and regulatory landscape shifts as key mechanisms enabling flatworm terrestrialization"

**Supplementary Figure 1.** Functional characterization of differentially expressed genes detected by proteomics (DGEs-prot). a) Treemaps representing the major clusters of enriched functions from *O. nungara* and *S. mediterranea.* b) Top pathways in which the DGEs-prot are involved.

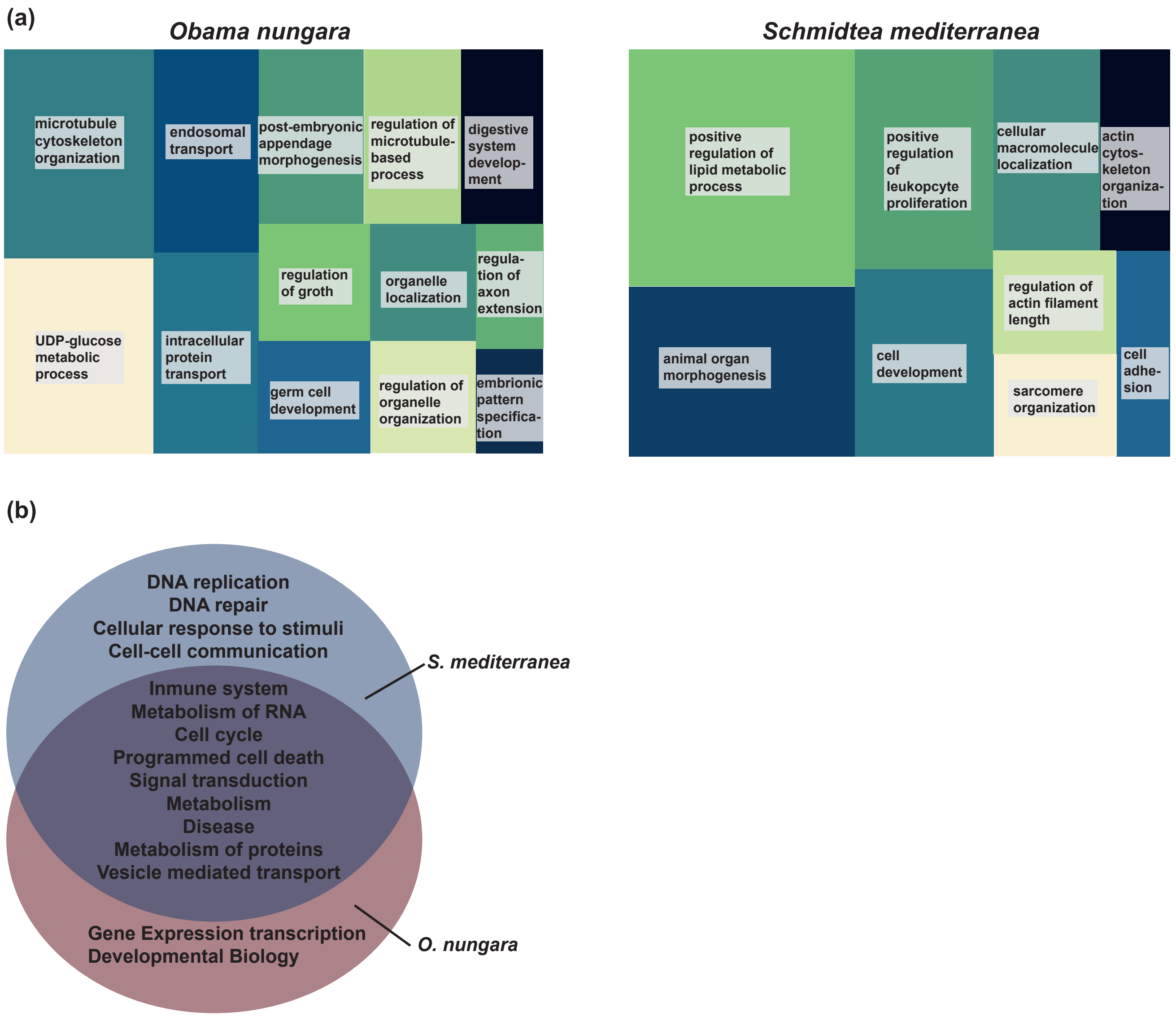
