## Supplementary Material 1 for "Genomic exaptation and regulatory landscape shifts as key mechanisms enabling flatworm terrestrialization"

**Supplementary Material 2.** Enrichments Gene Repertoire Evolution. Treemaps showing enriched GO terms (BP: Biological Processes, MF: Molecular Functions, CC: Cellular Components) in HOGs arisen in Tricladida (node 65), Continenticola (node64), Geoplanoidea (node 63), and Geoplaniidae (node 62)

TreeMap_Enrichm_node65_GAINED_BP

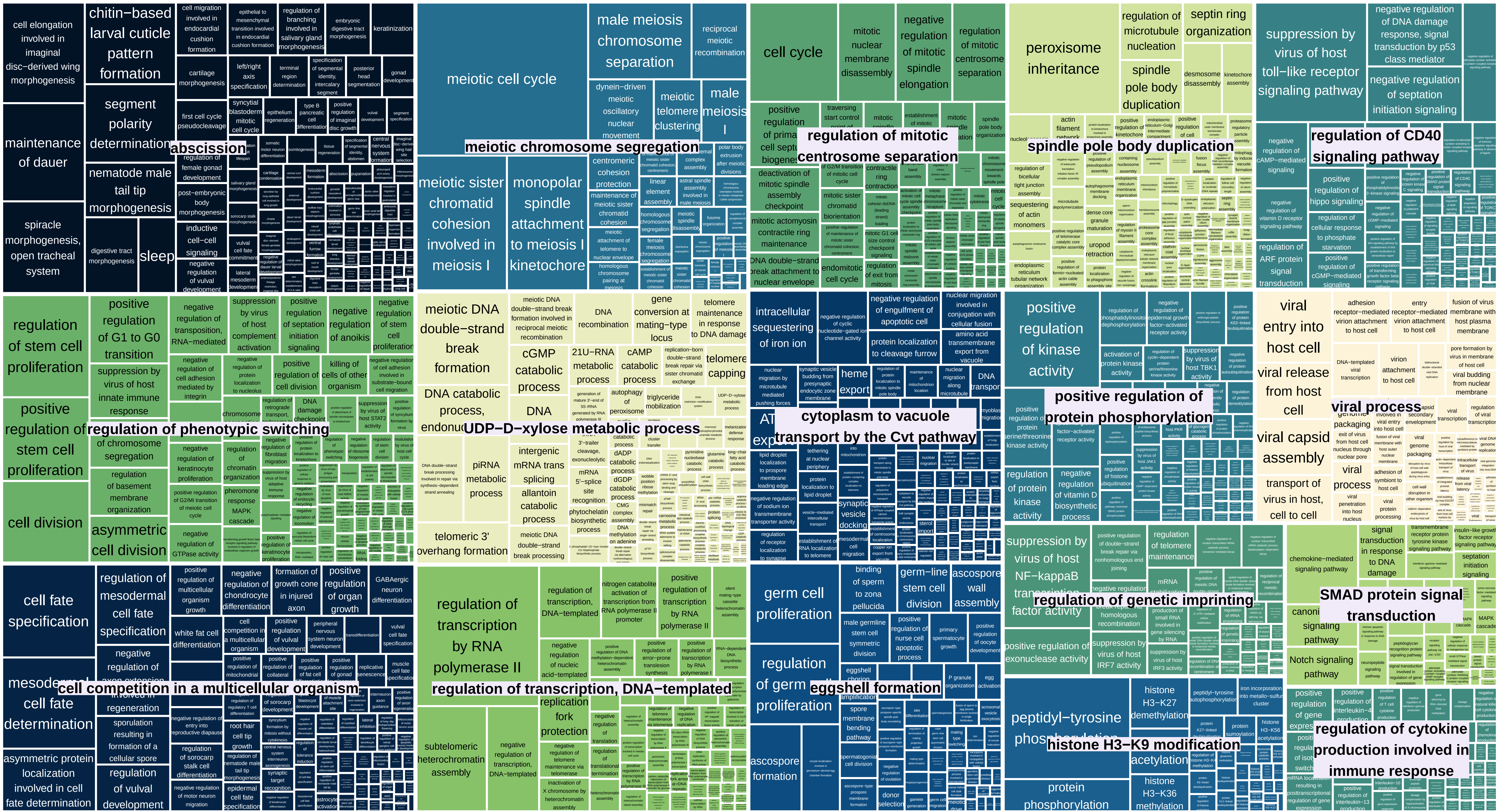

TreeMap_Enrichm_node65_GAINED_MF

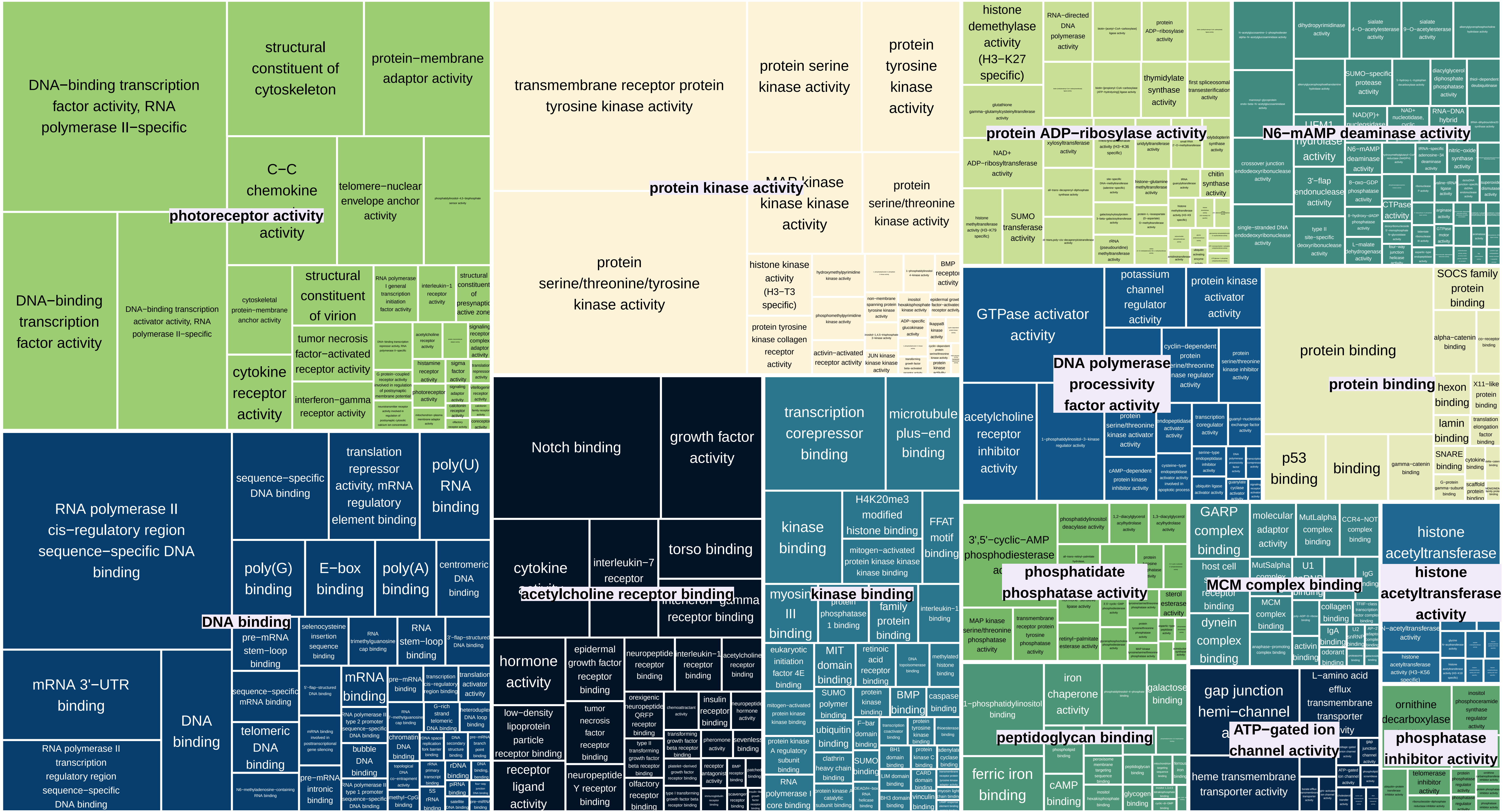

### TreeMap_Enrichm_node65_GAINED_CC

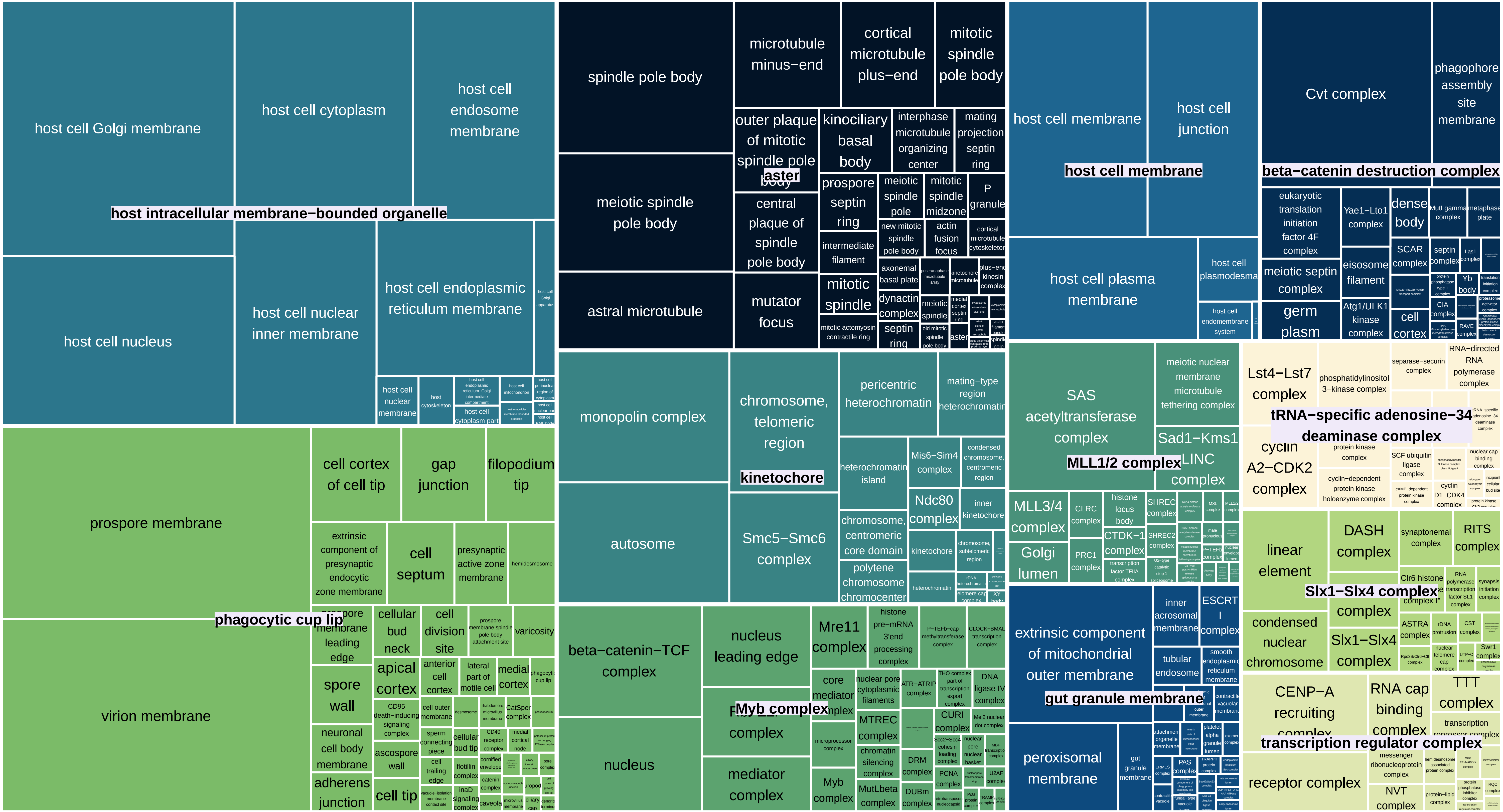

TreeMap_Enrichm_node64_GAINED_BP

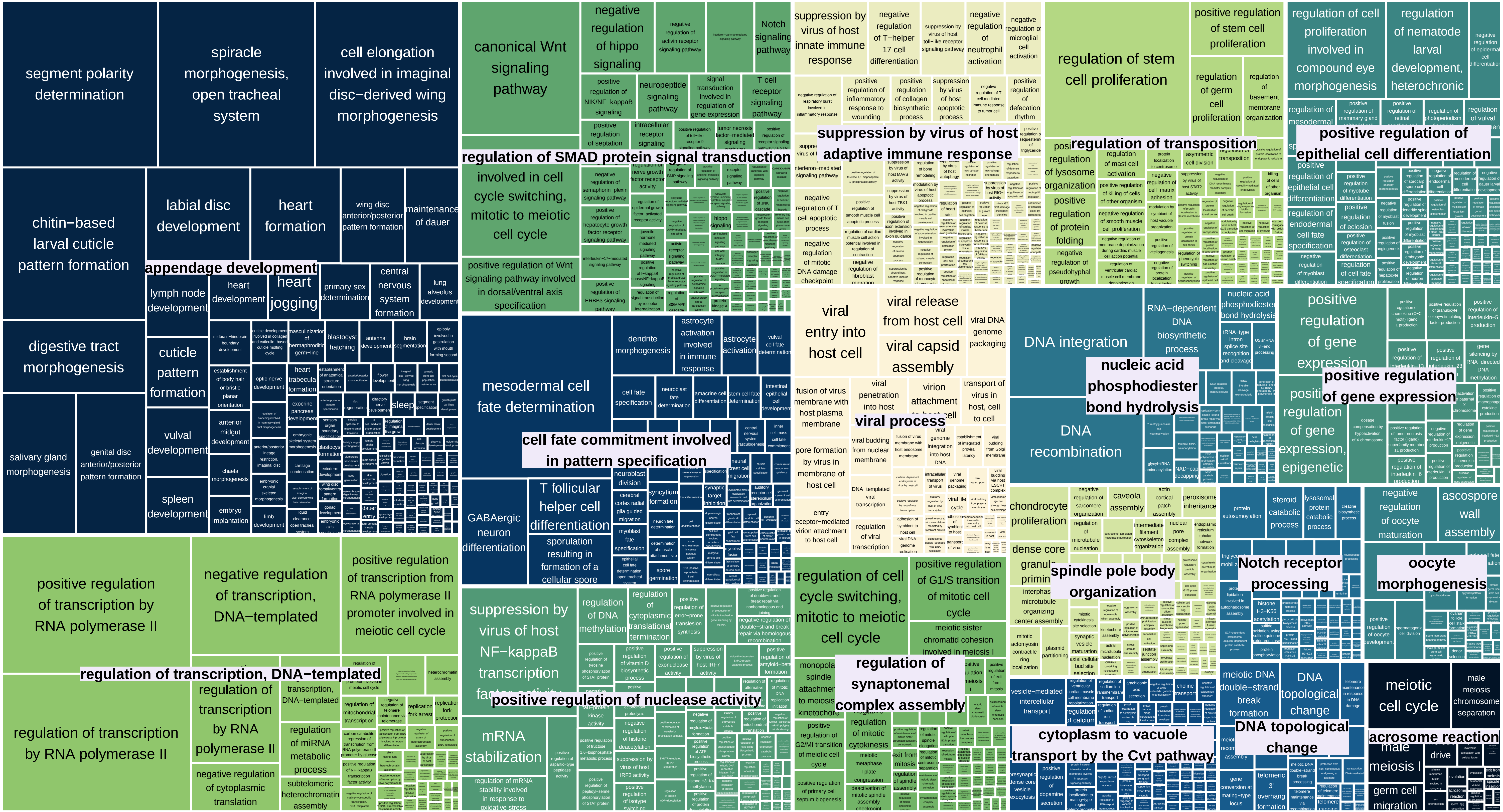

TreeMap_Enrichm_node64_GAINED_MF

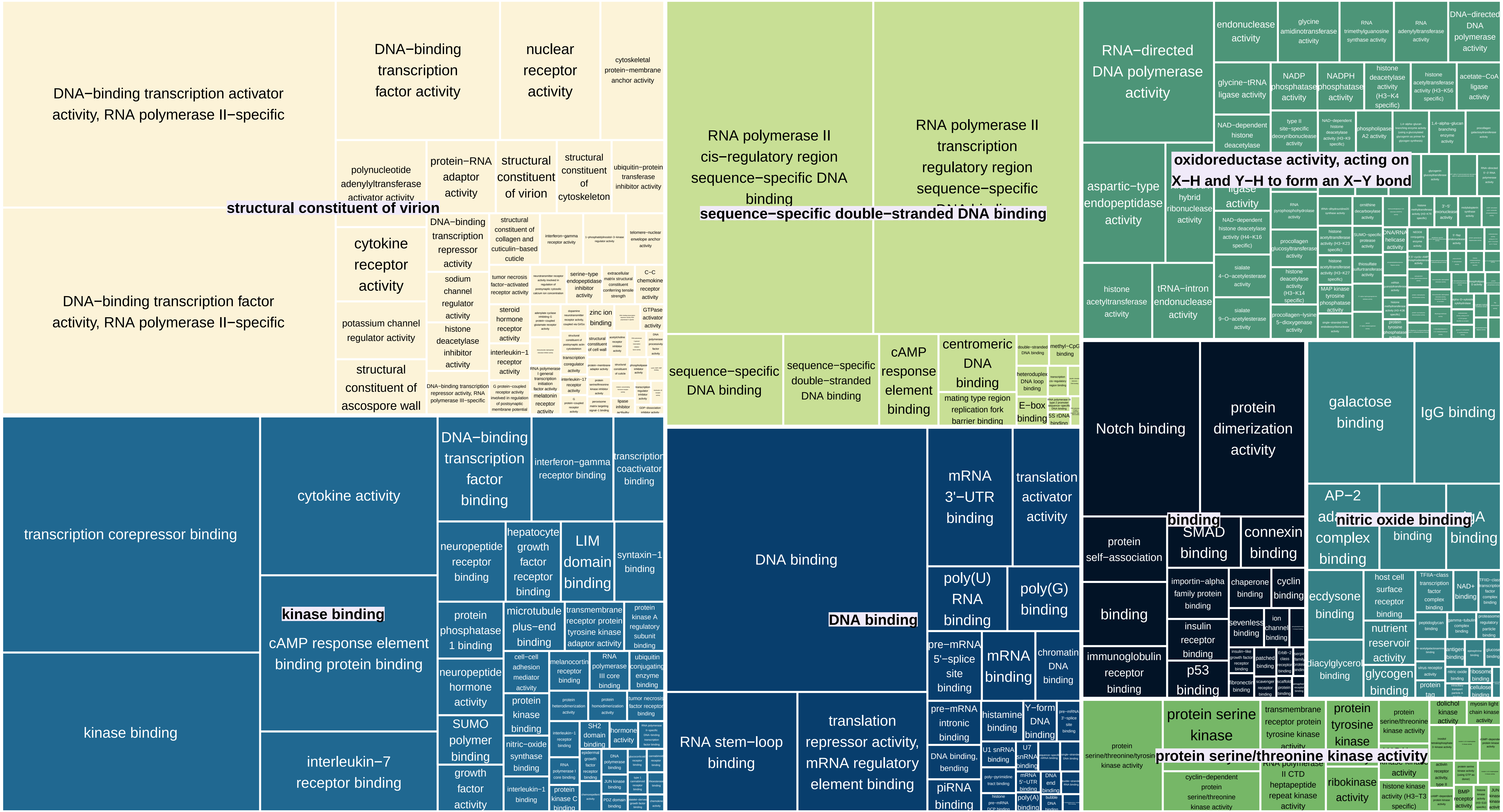

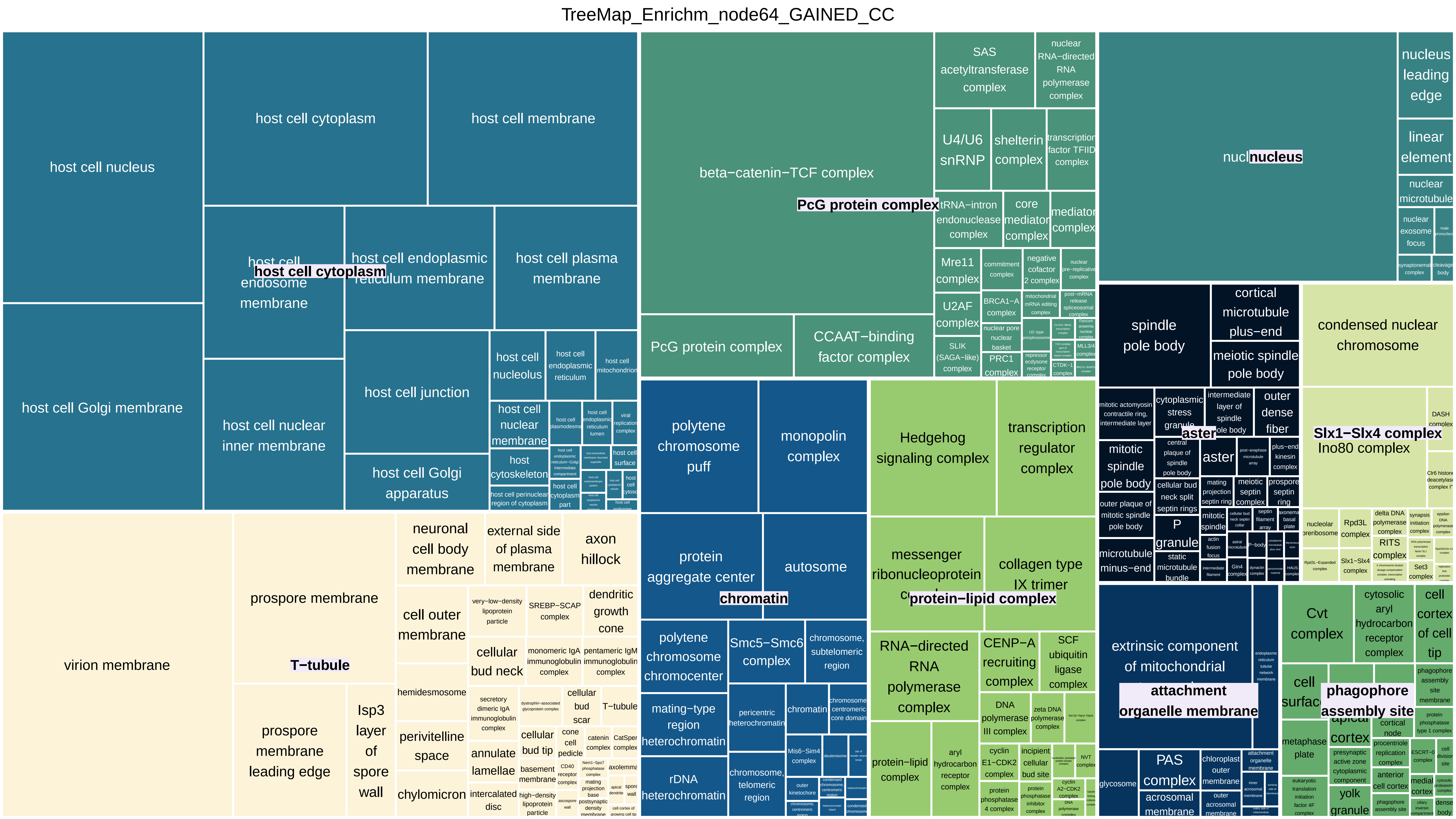
TreeMap_Enrichm_node64_GAINED_CC
host cell nucleus
host cell cytoplasm
host cell membrane
host cell endosome membrane
host cell endoplasmic reticulum membrane
host cell plasma membrane
host cell Golgi membrane
host cell nuclear inner membrane
host cell junction
host cell Golgi apparatus
host cell nucleolus
host cell endoplasmic reticulum
host cell mitochondrion
host cell nuclear membrane
host cytoskeleton
beta−catenin−TCF complex
PcG protein complex
CCAAT−binding factor complex
SAS acetyltransferase complex
nuclear RNA−directed RNA polymerase complex
U4/U6 snRNP
shelterin complex
transcription factor TFIID complex
tRNA−intron endonuclease complex
core mediator complex
mediator complex
Mre11 complex
U2AF complex
SLIK (SAGA−like) complex
PRC1 complex
BRCA1−A complex
nucleus
nucleus leading edge
linear element
nuclear microtubule
nuclear exosome focus
spindle pole body
cortical microtubule plus−end
meiotic spindle pole body
condensed nuclear chromosome
Slx1−Slx4 complex
Ino80 complex
aster
mitotic spindle pole body
outer dense fiber
P granule
microtubule minus−end
virion membrane
prospore membrane
T−tubule
prospore membrane leading edge
Isp3 layer of spore wall
neuronal cell body membrane
external side of plasma membrane
axon hillock
cell outer membrane
cellular bud neck
dendritic growth cone
hemidesmosome
perivitelline space
chylomicron
annulate lamellae
intercalated disc
polytene chromosome puff
monopolin complex
protein aggregate center
autosome
chromatin
polytene chromosome chromocenter
Smc5−Smc6 complex
chromosome, subtelomeric region
mating−type region heterochromatin
rDNA heterochromatin
pericentric heterochromatin
chromosome, telomeric region
Hedgehog signaling complex
transcription regulator complex
messenger ribonucleoprotein complex
collagen type IX trimer
protein−lipid complex
RNA−directed RNA polymerase complex
CENP−A recruiting complex
SCF ubiquitin ligase complex
protein−lipid complex
aryl hydrocarbon receptor complex
extrinsic component of mitochondrial
attachment organelle membrane
PAS complex
acrosomal membrane
glycosome
Cvt complex
cytosolic aryl hydrocarbon receptor complex
cell cortex of cell tip
cell surface
phagophore assembly site
metaphase plate
yolk granule

TreeMap_Enrichm_node63_GAINED_BP

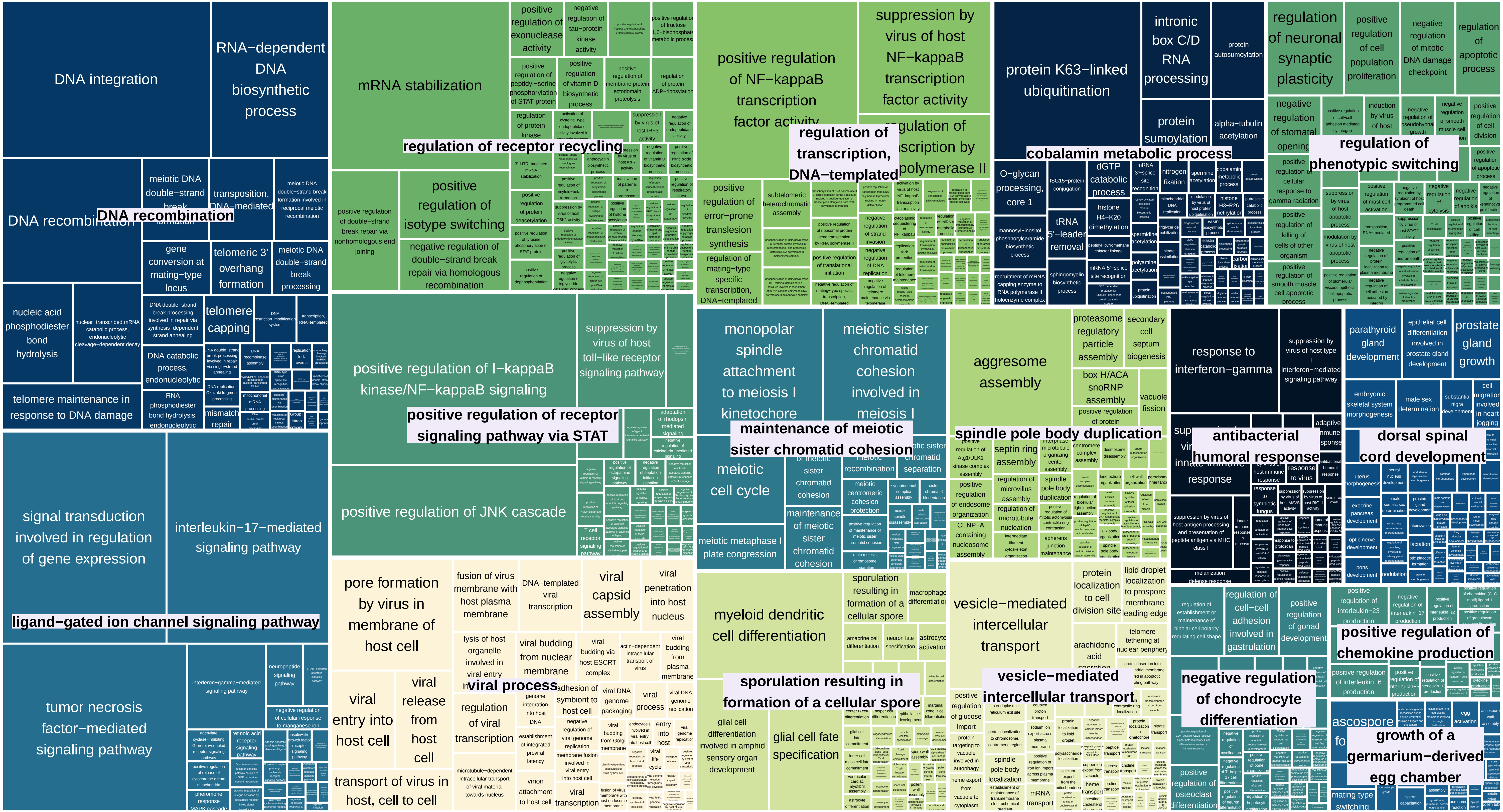

TreeMap_Enrichm_node63_GAINED_MF

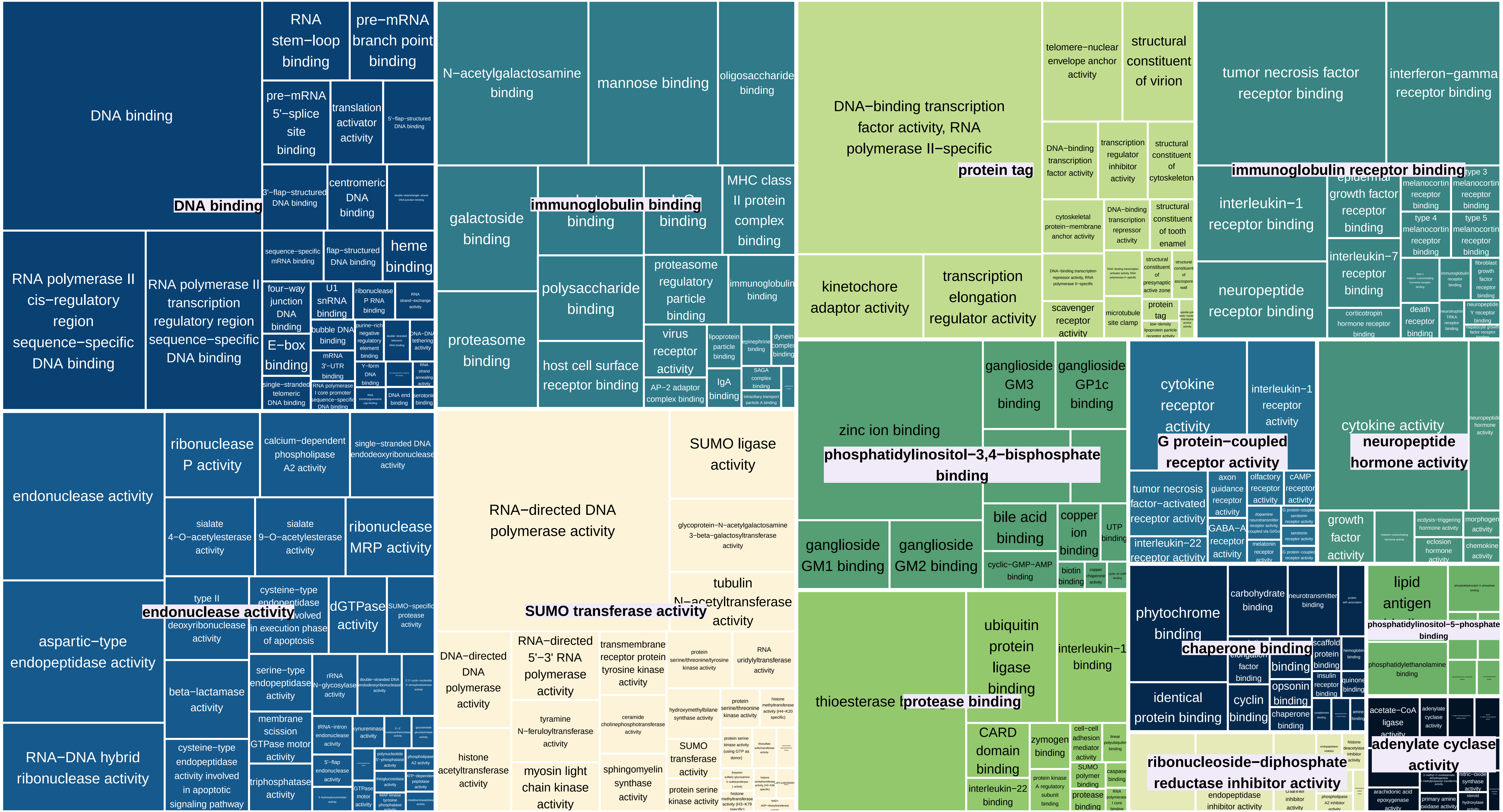

TreeMap_Enrichm_node63_GAINED_CC

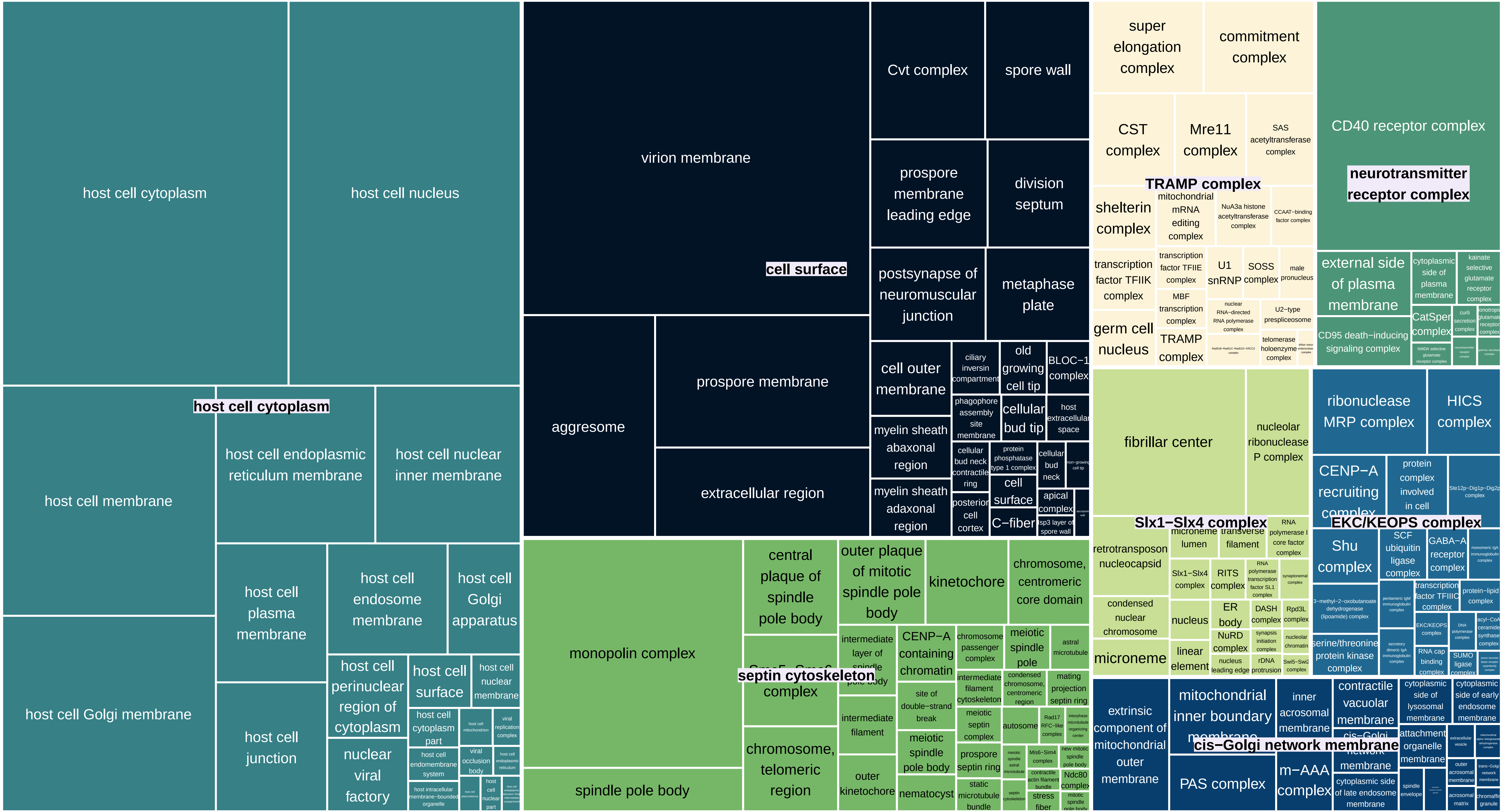

TreeMap_Enrichm_node62_GAINED_BP

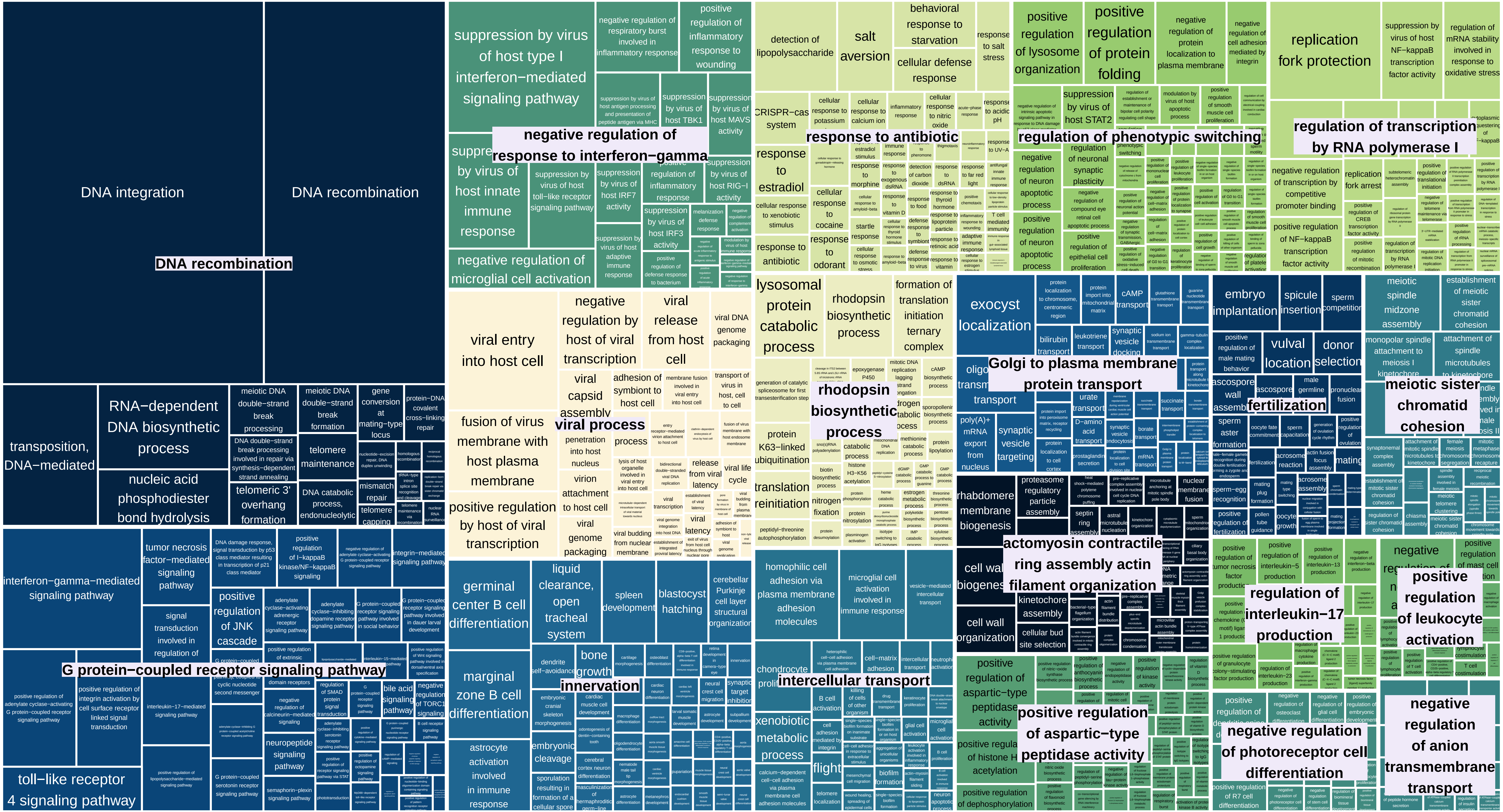

TreeMap_Enrichm_node62_GAINED_MF

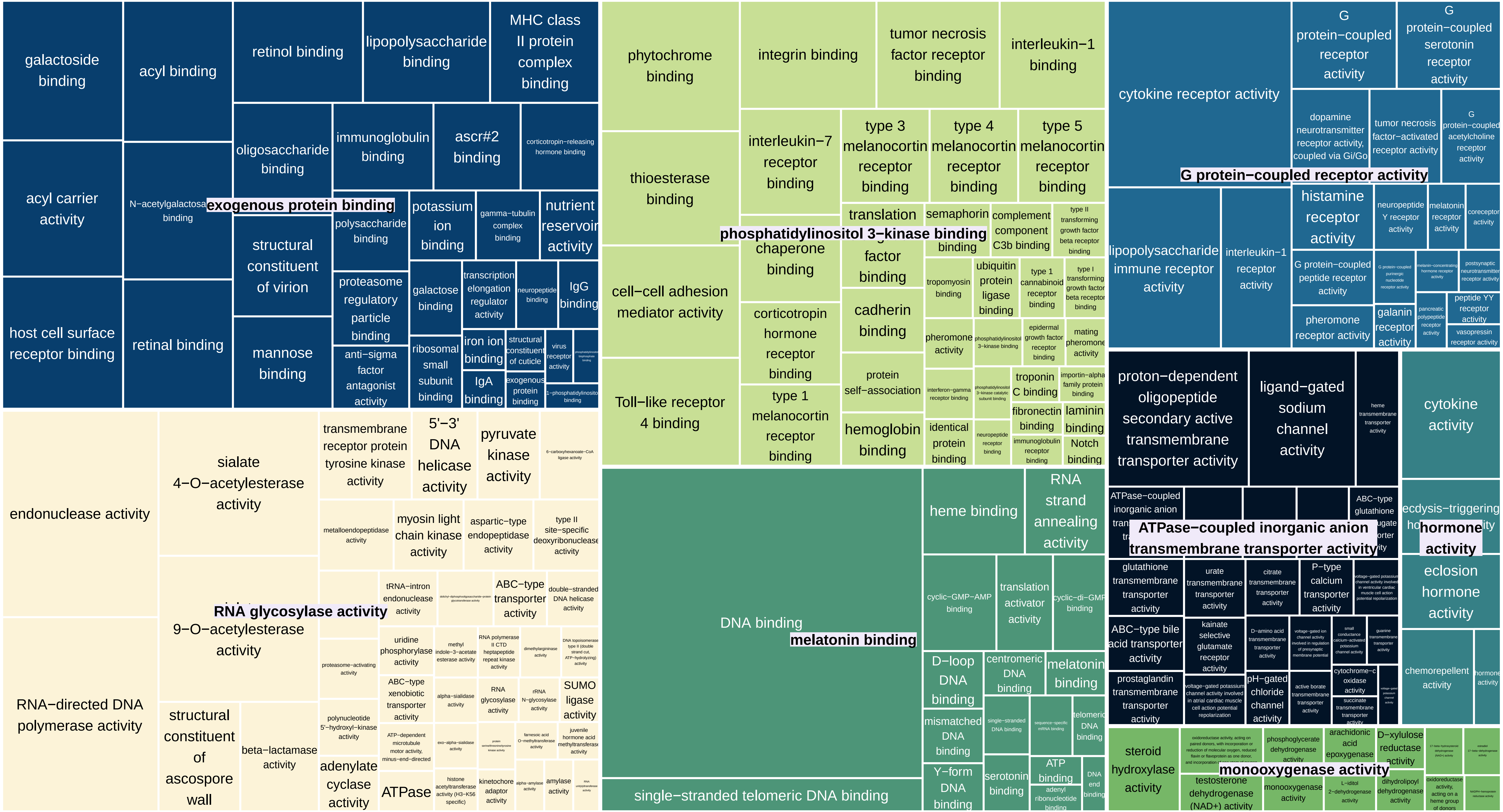

TreeMap_Enrichm_node62_GAINED_CC

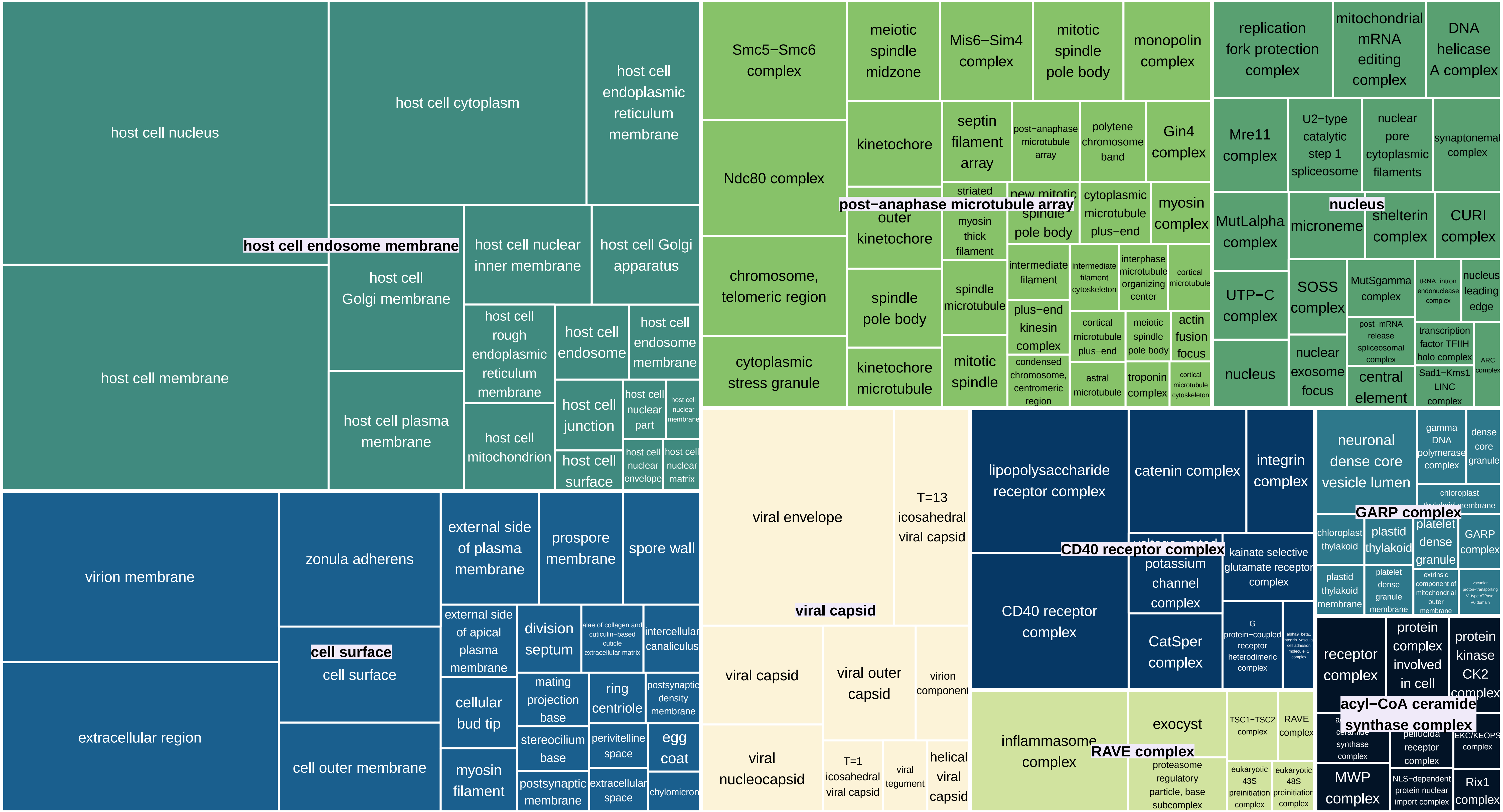

TreeMap_Enrichm_node62_DUPLICATED_balanced_BP

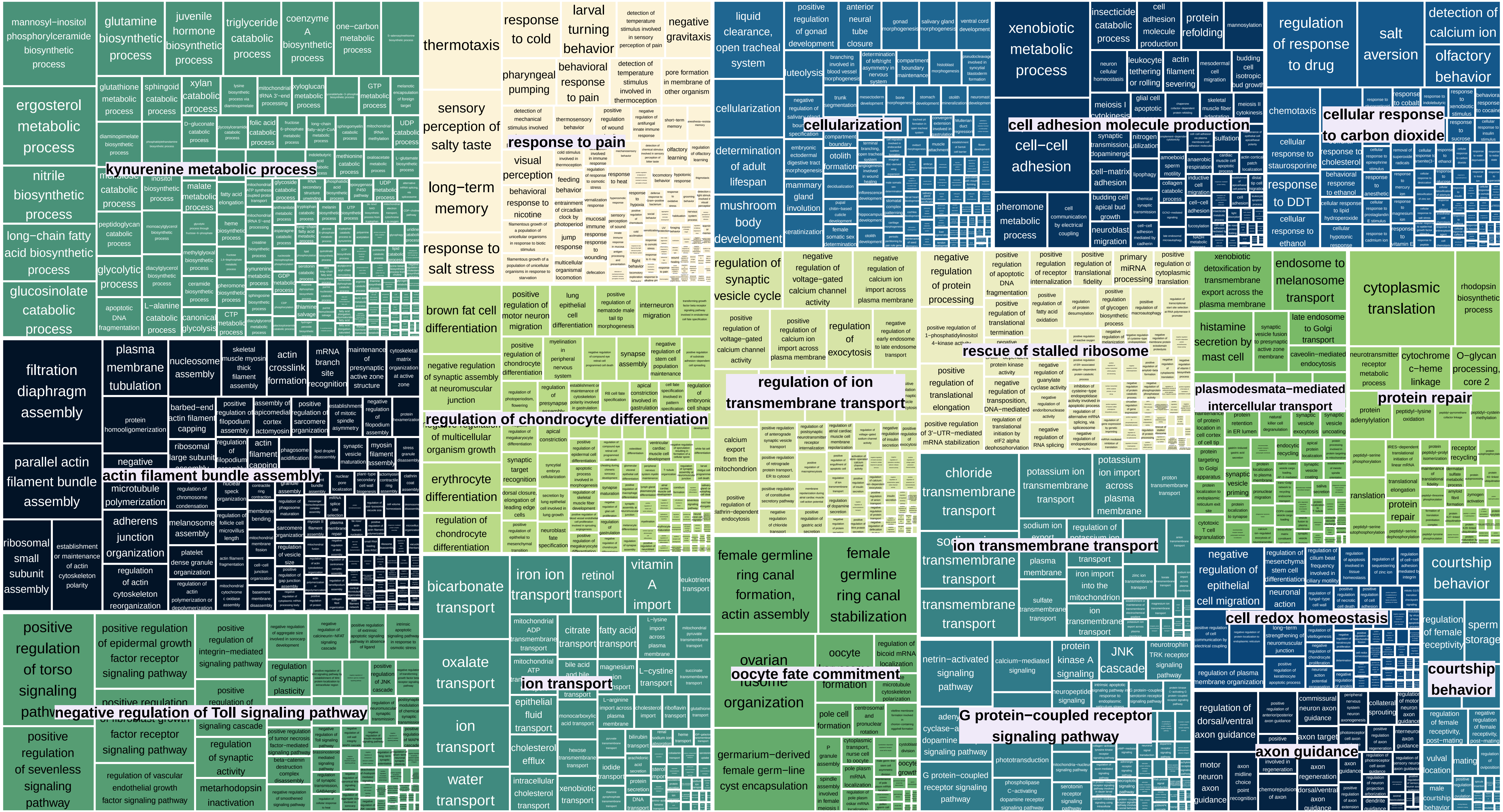

TreeMap_Enrichm_node62_DUPLICATED_balanced_MF

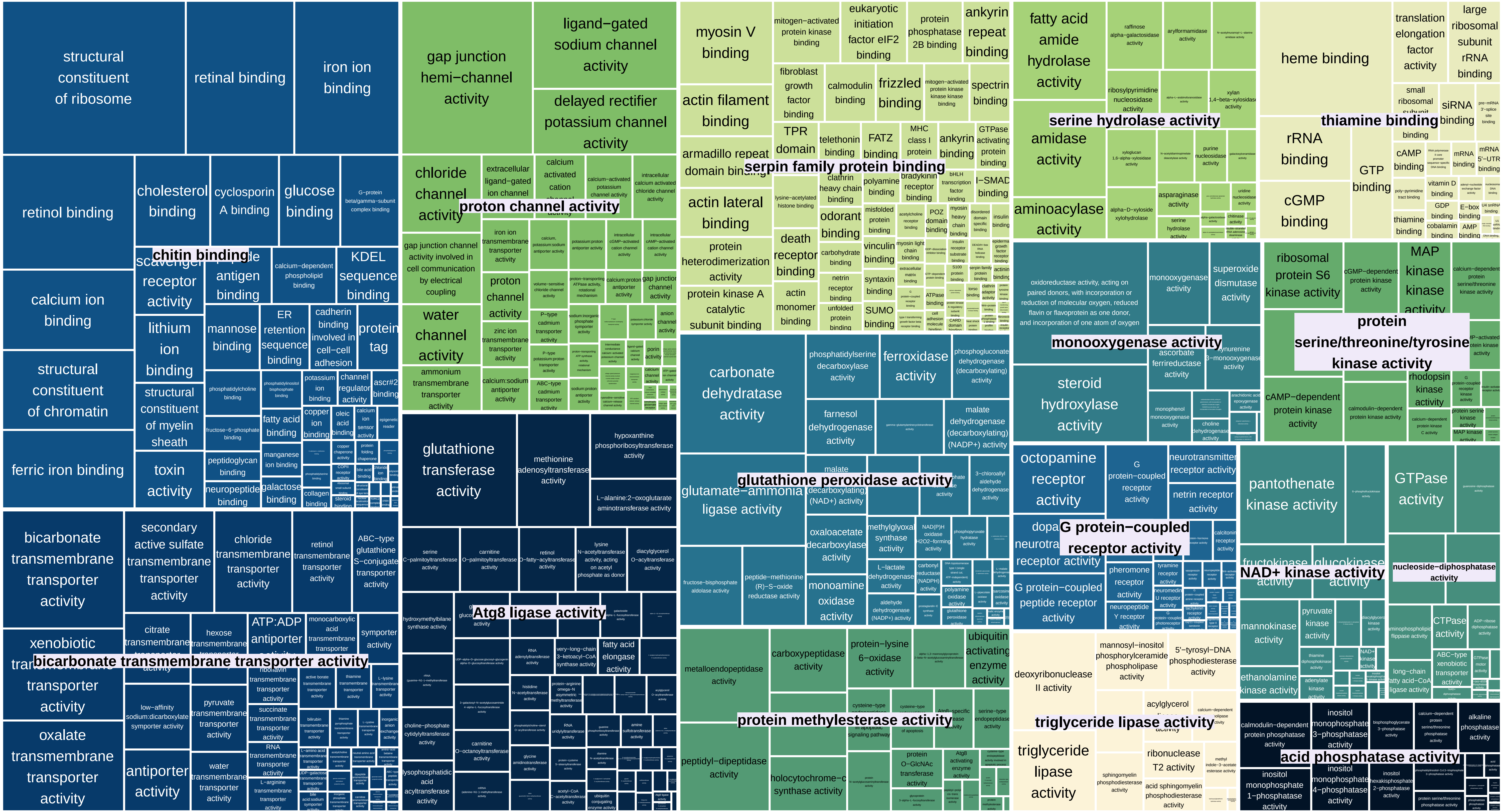

TreeMap_Enrichm_node62_DUPLICATED_balanced_CC

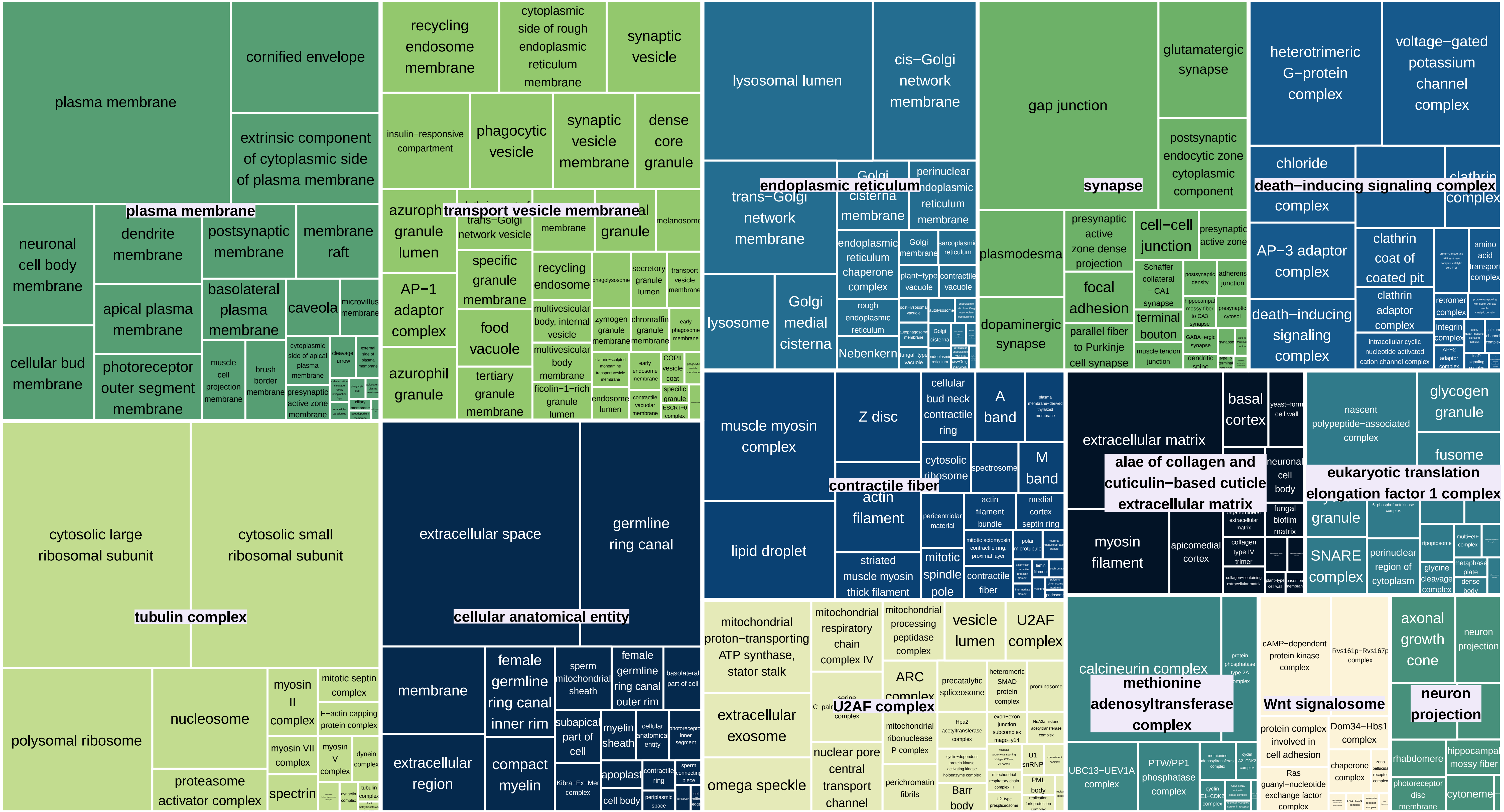

TreeMap_Enrichm_node62_EXPANDED_balanced_BP

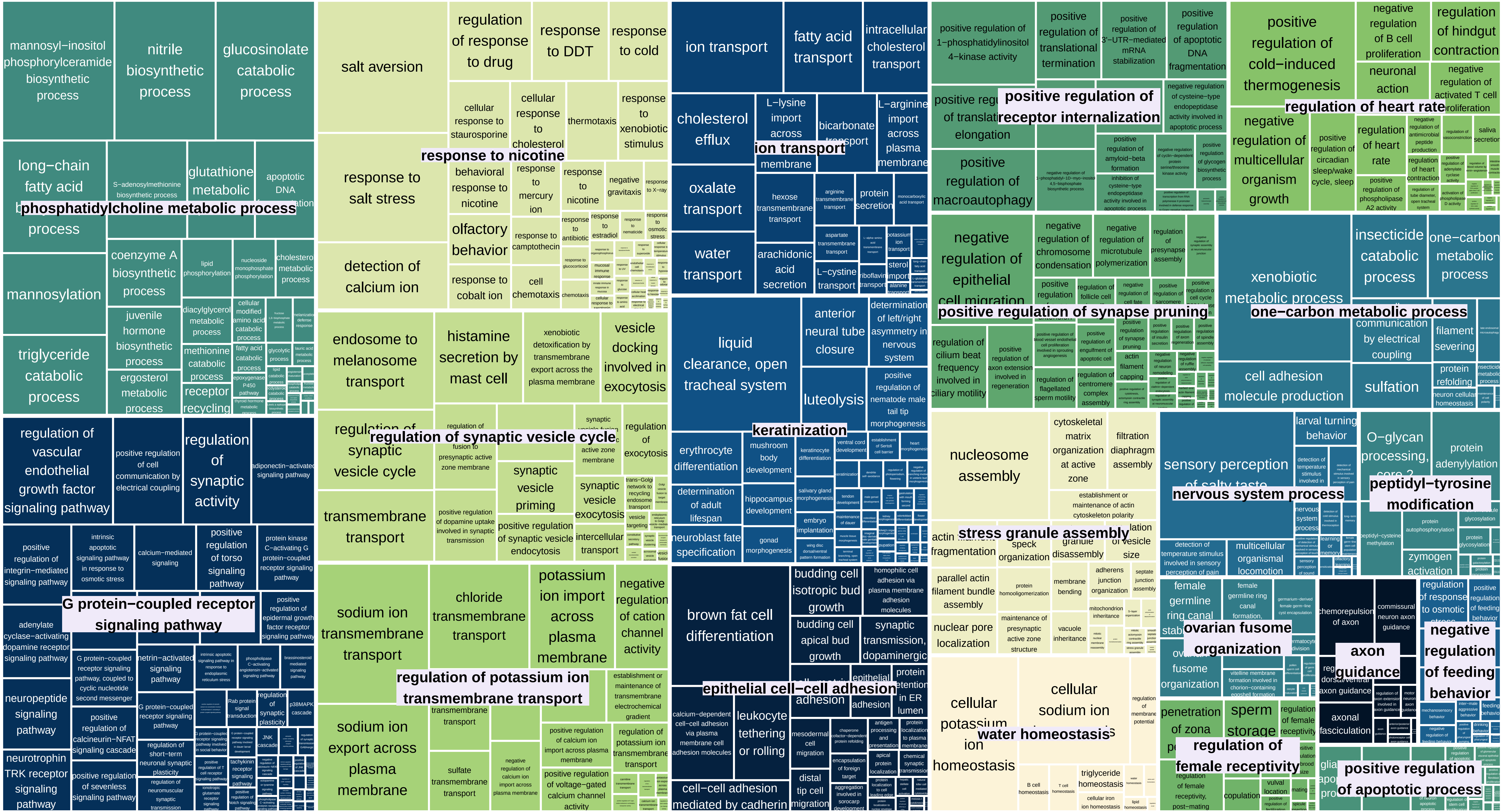

TreeMap_Enrichm_node62_EXPANDED_balanced_MF

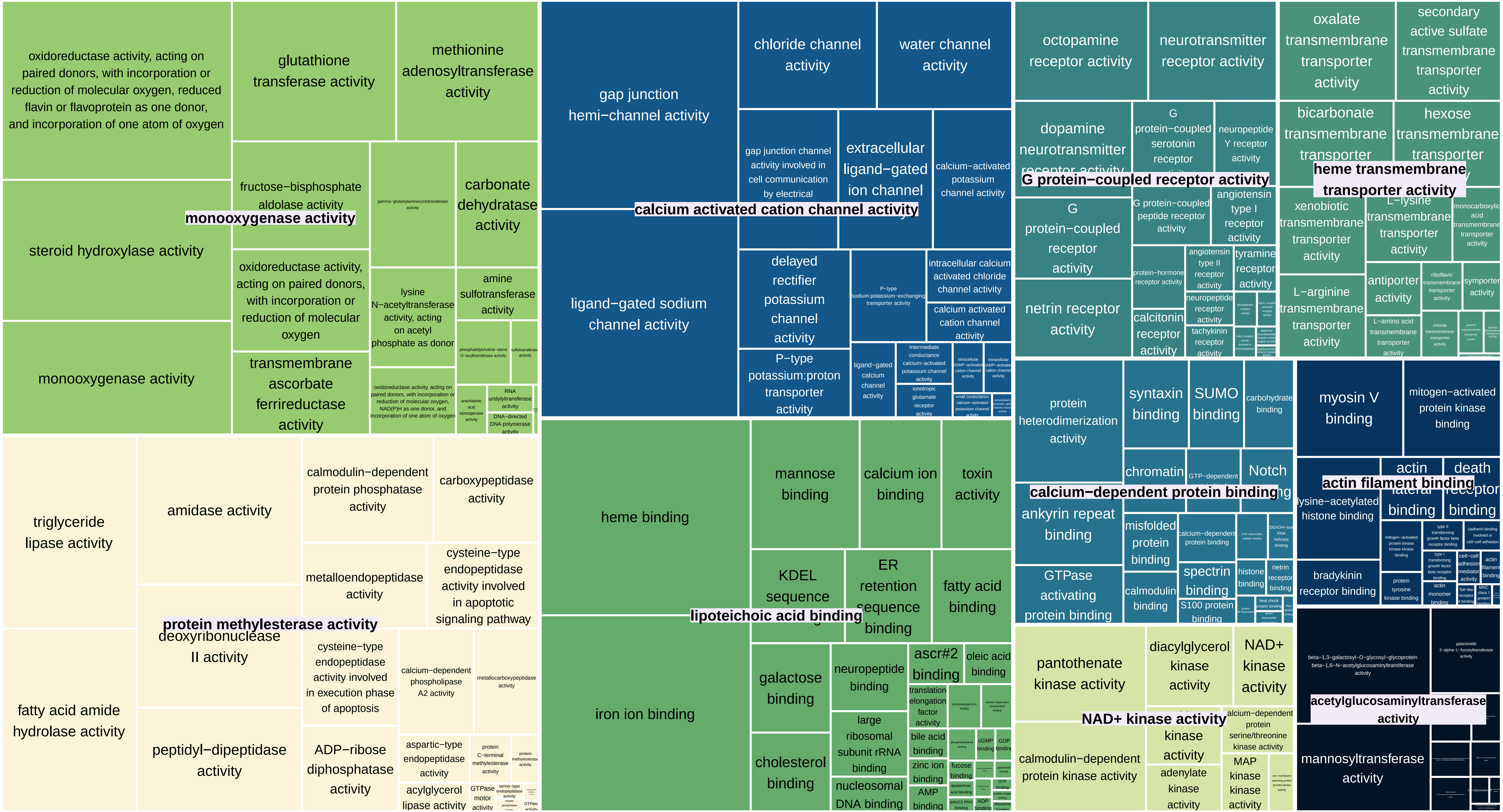

TreeMap_Enrichm_node62_EXPANDED_balanced_CC

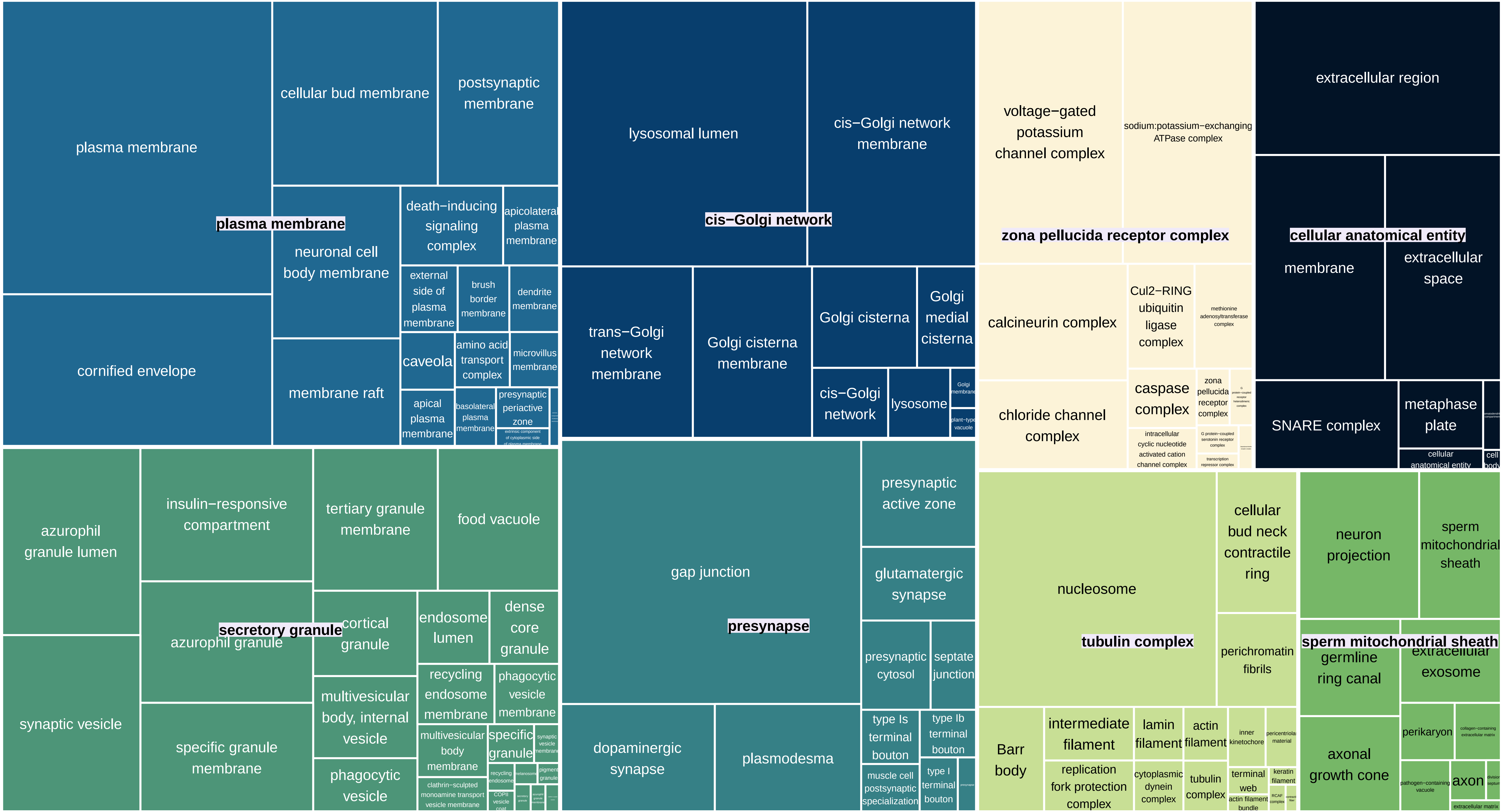
