## Supplementary Material 2 for "Genomic exaptation and regulatory landscape shifts as key mechanisms enabling flatworm terrestrialization"

**Supplementary Material 3.** Enrichments Proteomics. Treemaps showing enriched GO terms (BP: Biological Processes, MF: Molecular Functions, CC: Cellular Components) in proteins detected in ONUN and SMED. Additionally, the enrichment of HOGs including proteins detected in ONUN, proteins detected in SMED, and proteins detencted in both species

TreeMap_Enrichm_ONUN_prot_BP

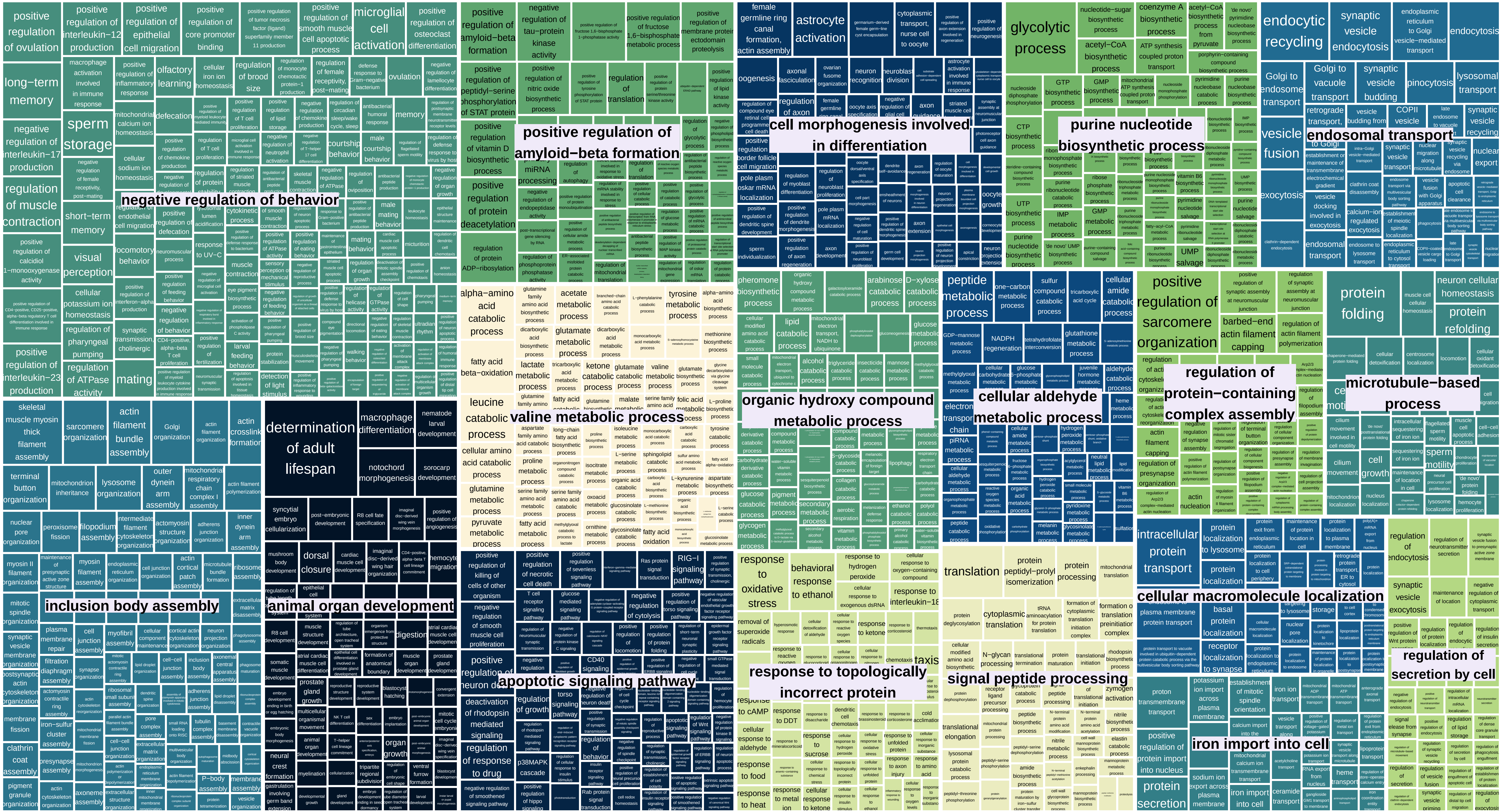

TreeMap_Enrichm_ONUN_prot_MF

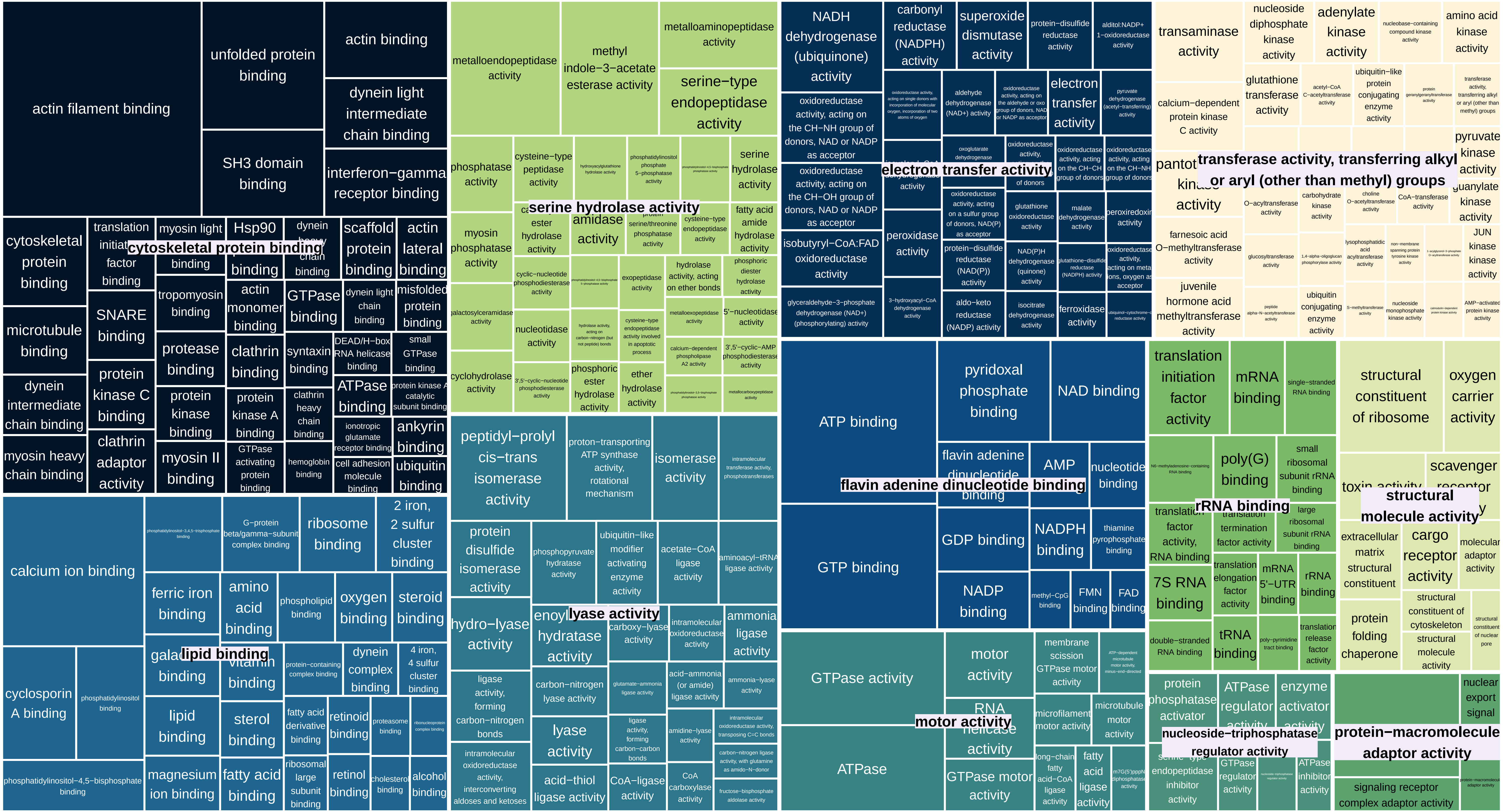

TreeMap_Enrichm_ONUN_prot_CC

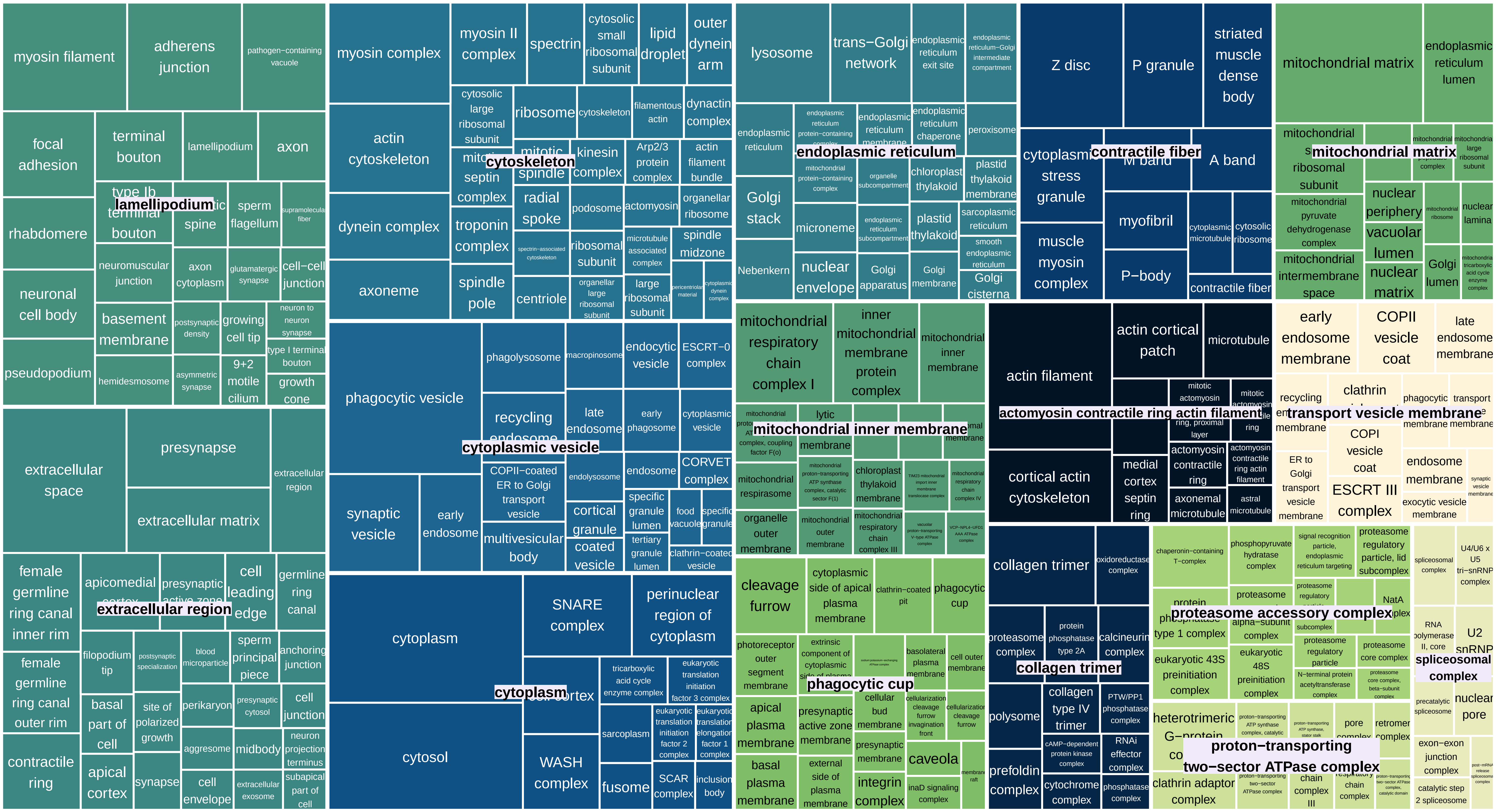

TreeMap_Enrichm_SMED_prot_BP

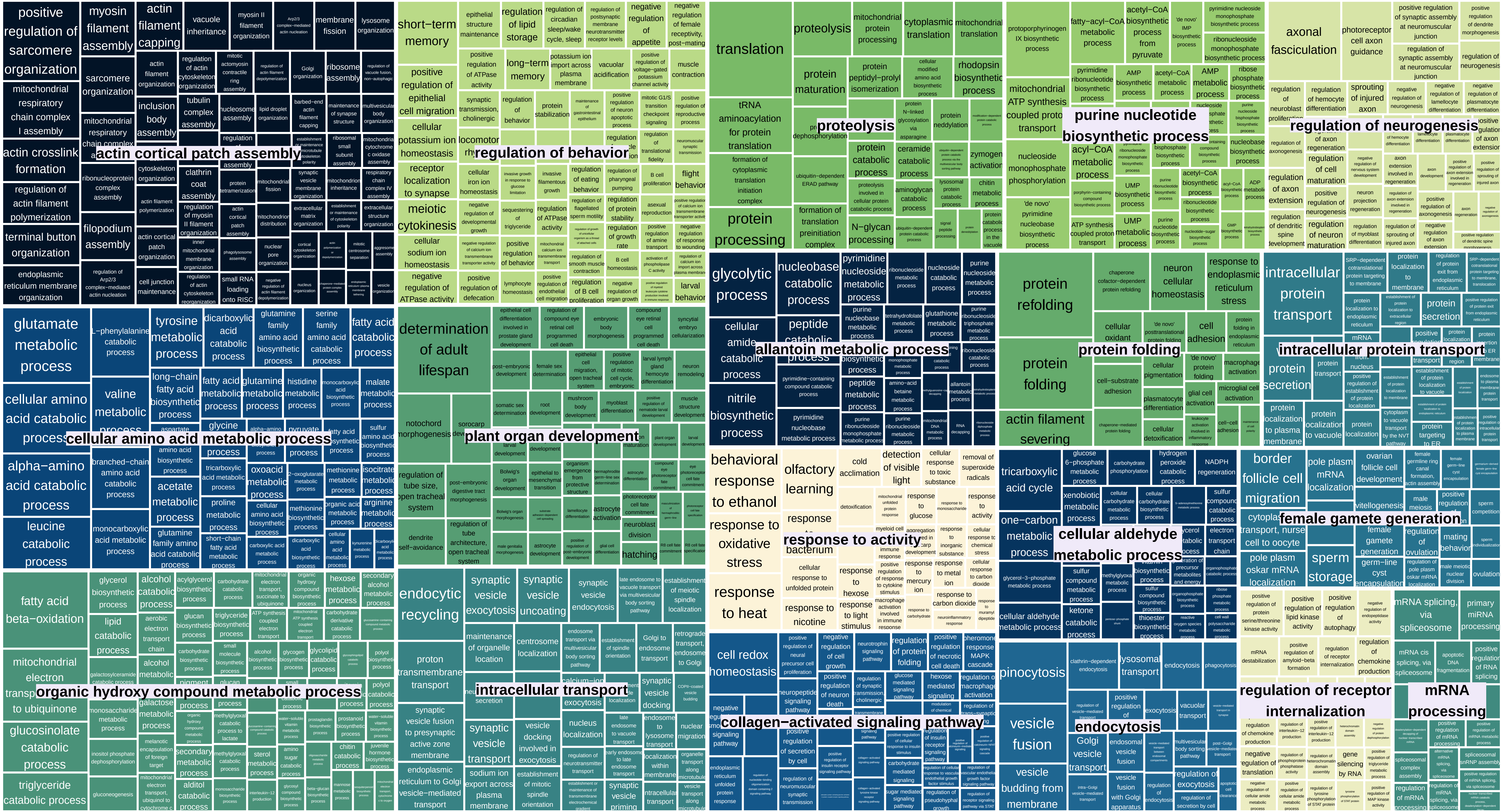

TreeMap_Enrichm_SMED_prot_MF

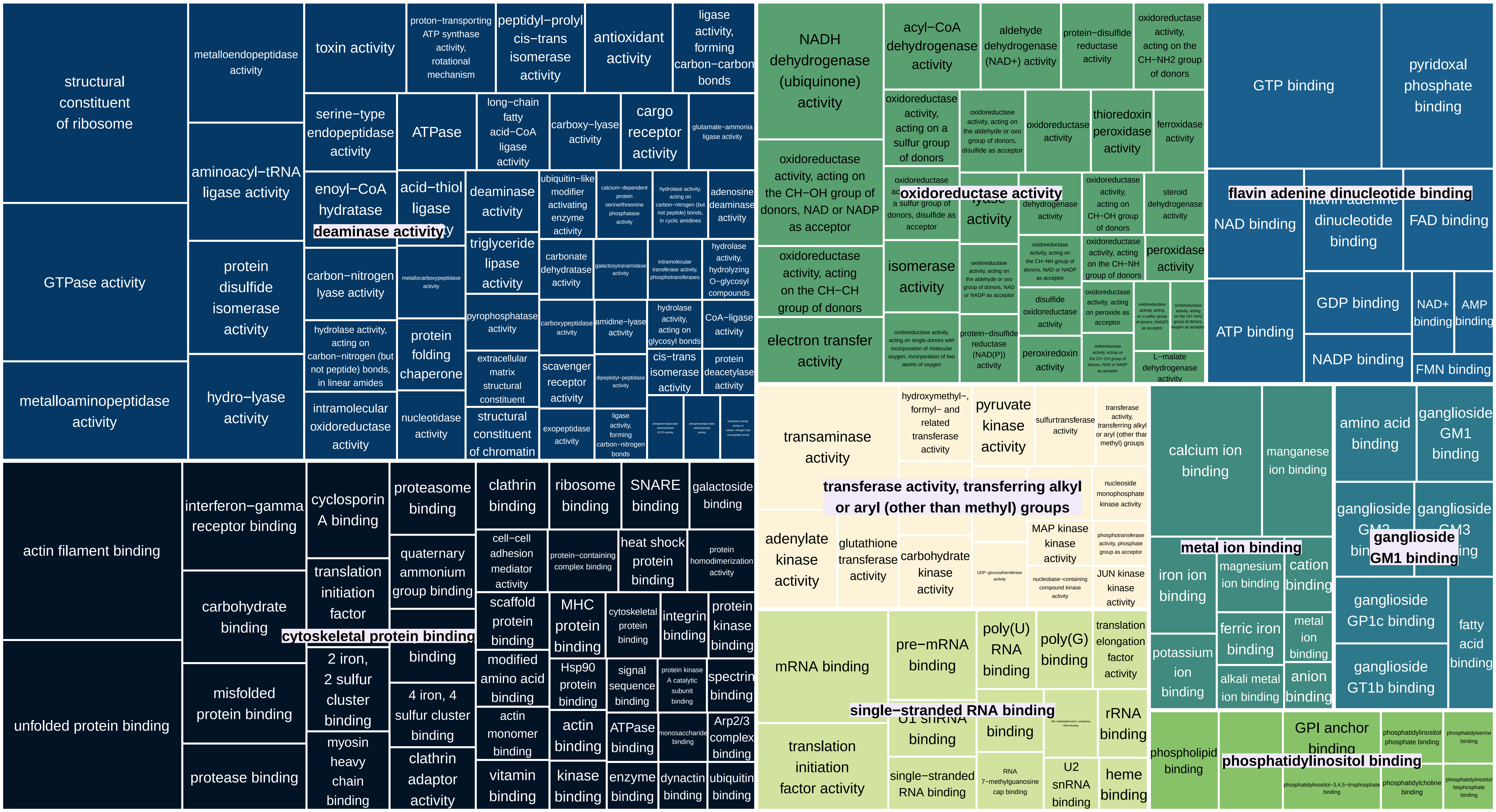

TreeMap_Enrichm_SMED_prot_CC

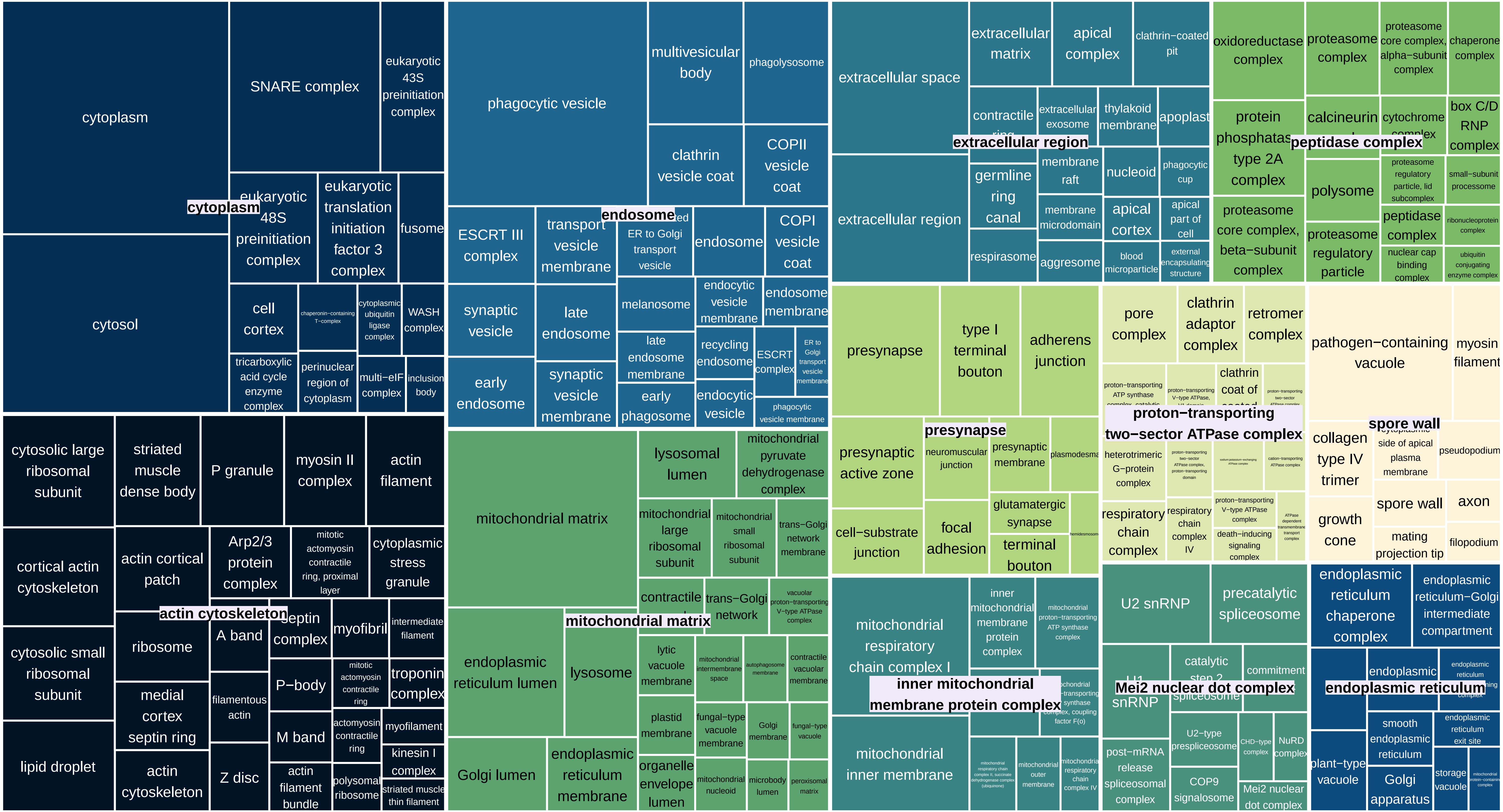

TreeMap_Enrichm_ONUN_HOGs_BP

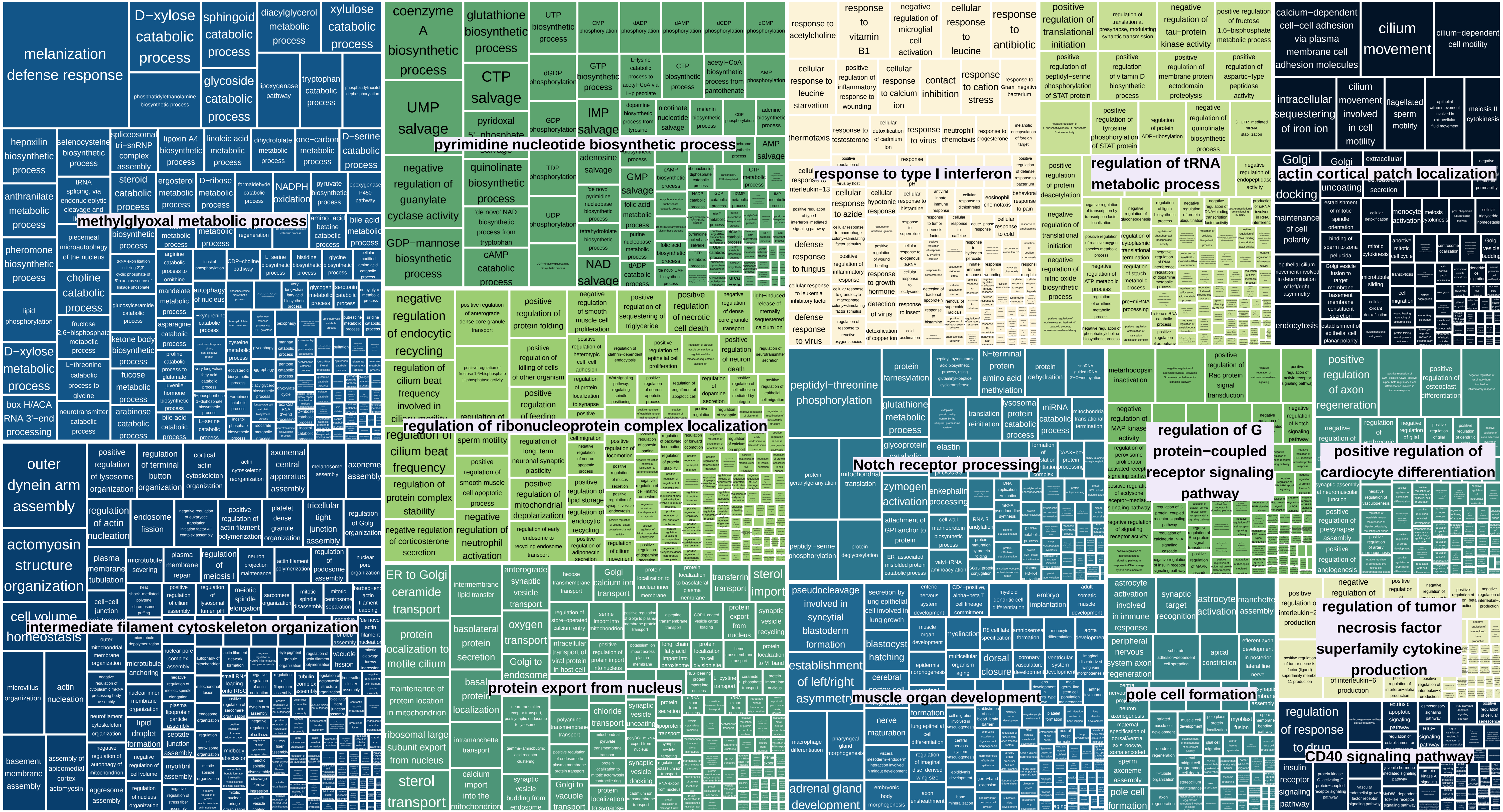

TreeMap_Enrichm_ONUN_HOGs_MF

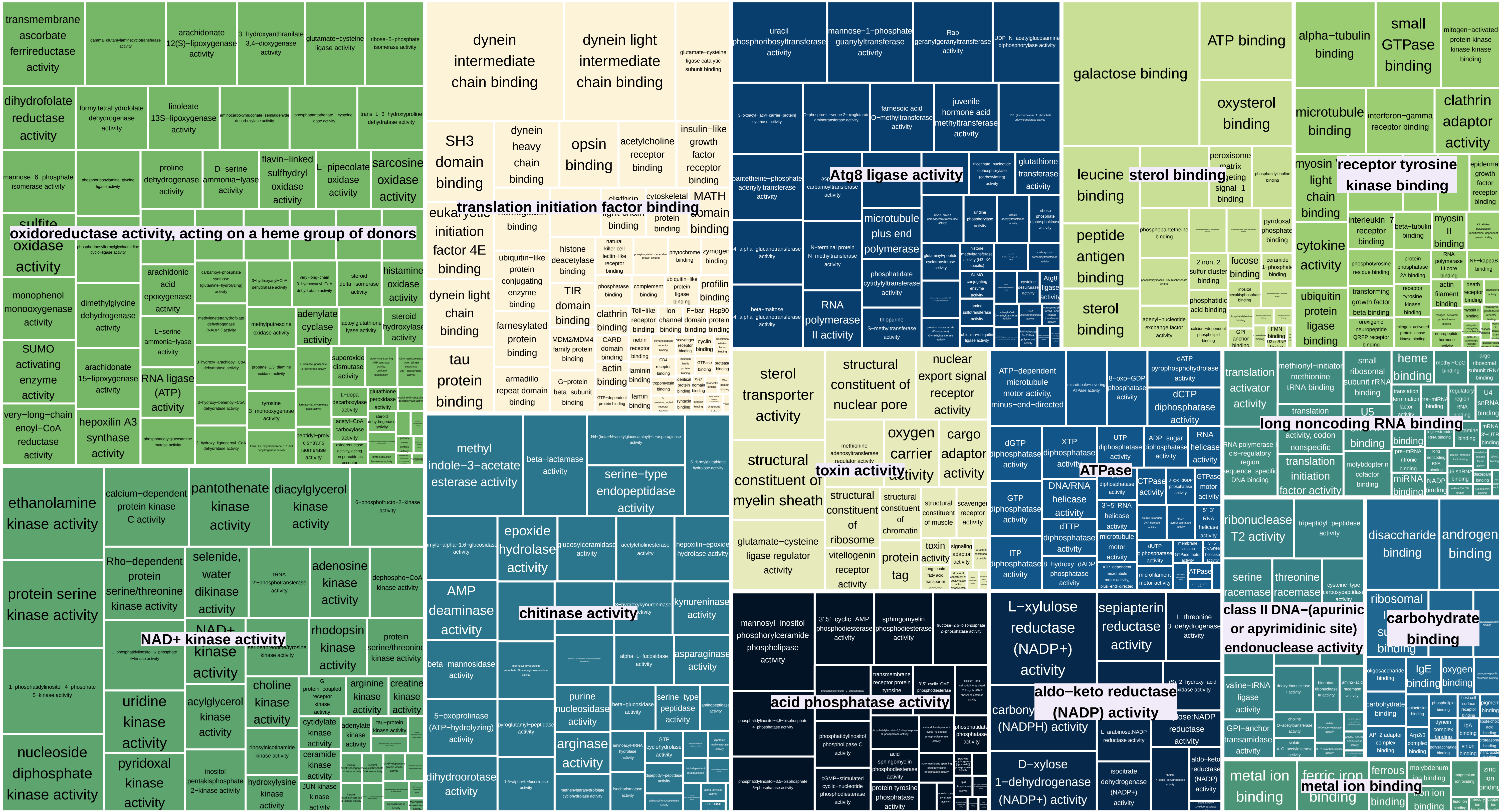

TreeMap_Enrichm_ONUN_HOGs_CC

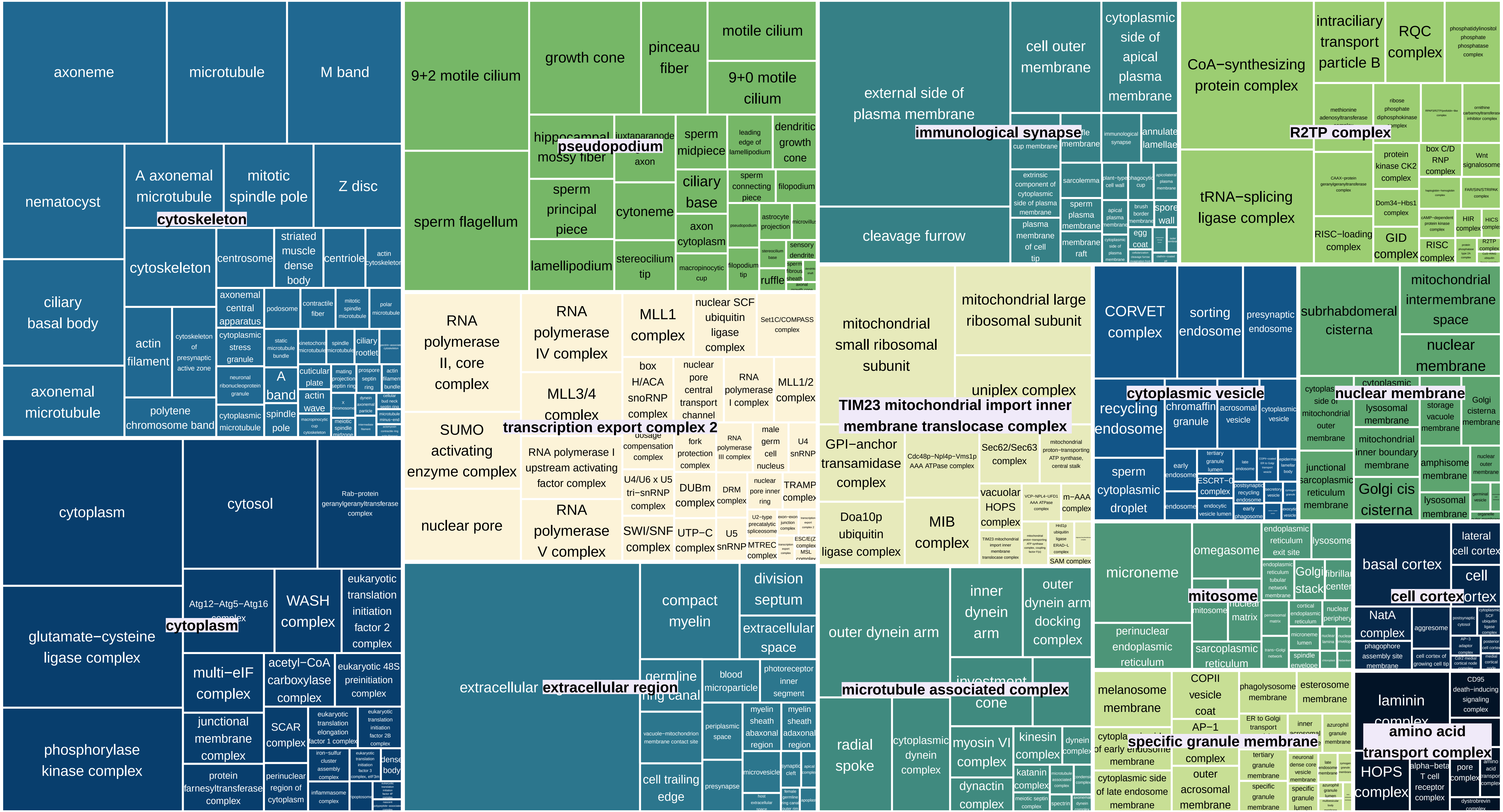

TreeMap_Enrichm_SMED_HOGs_BP

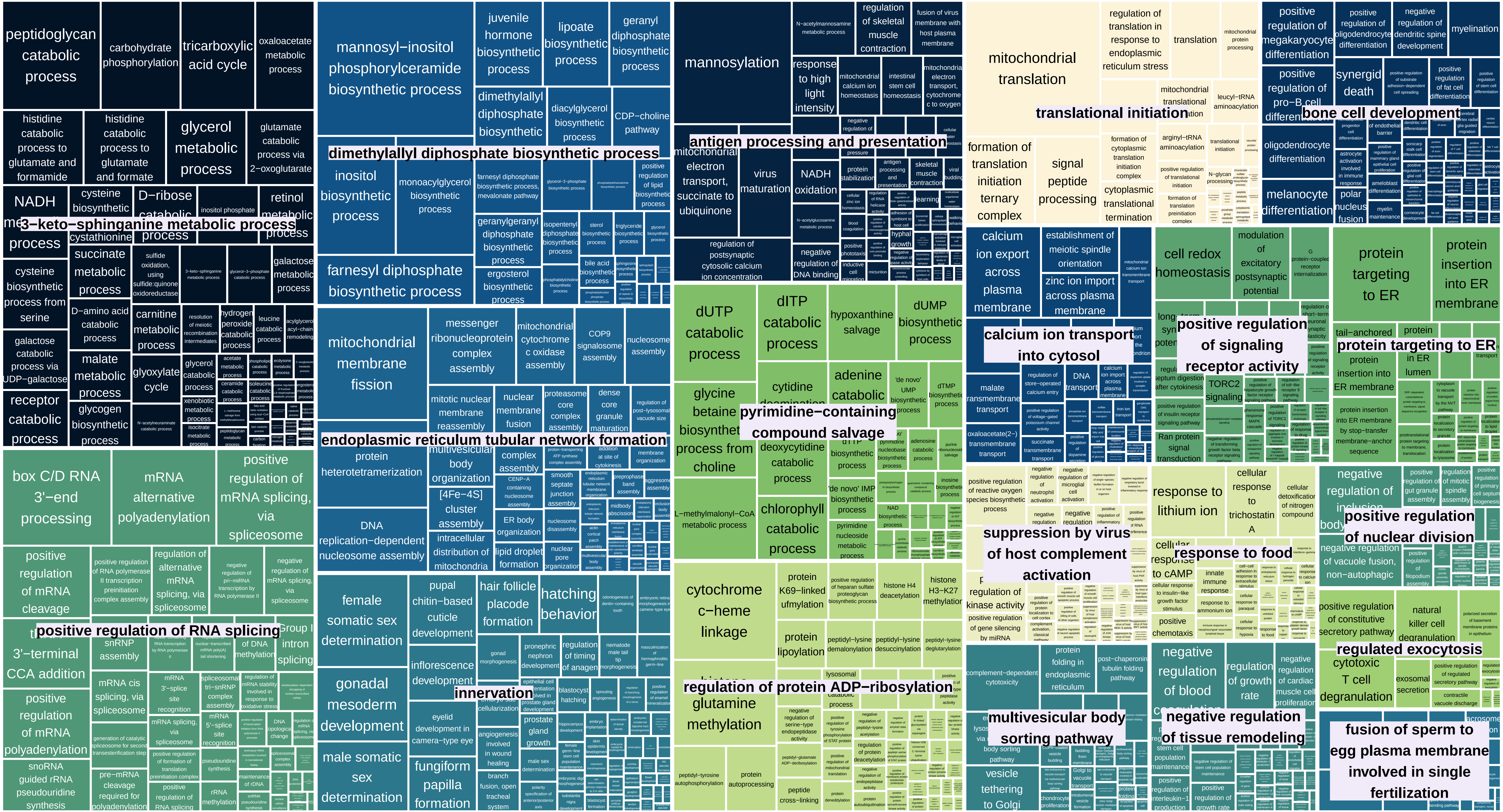

TreeMap_Enrichm_SMED_HOGs_MF

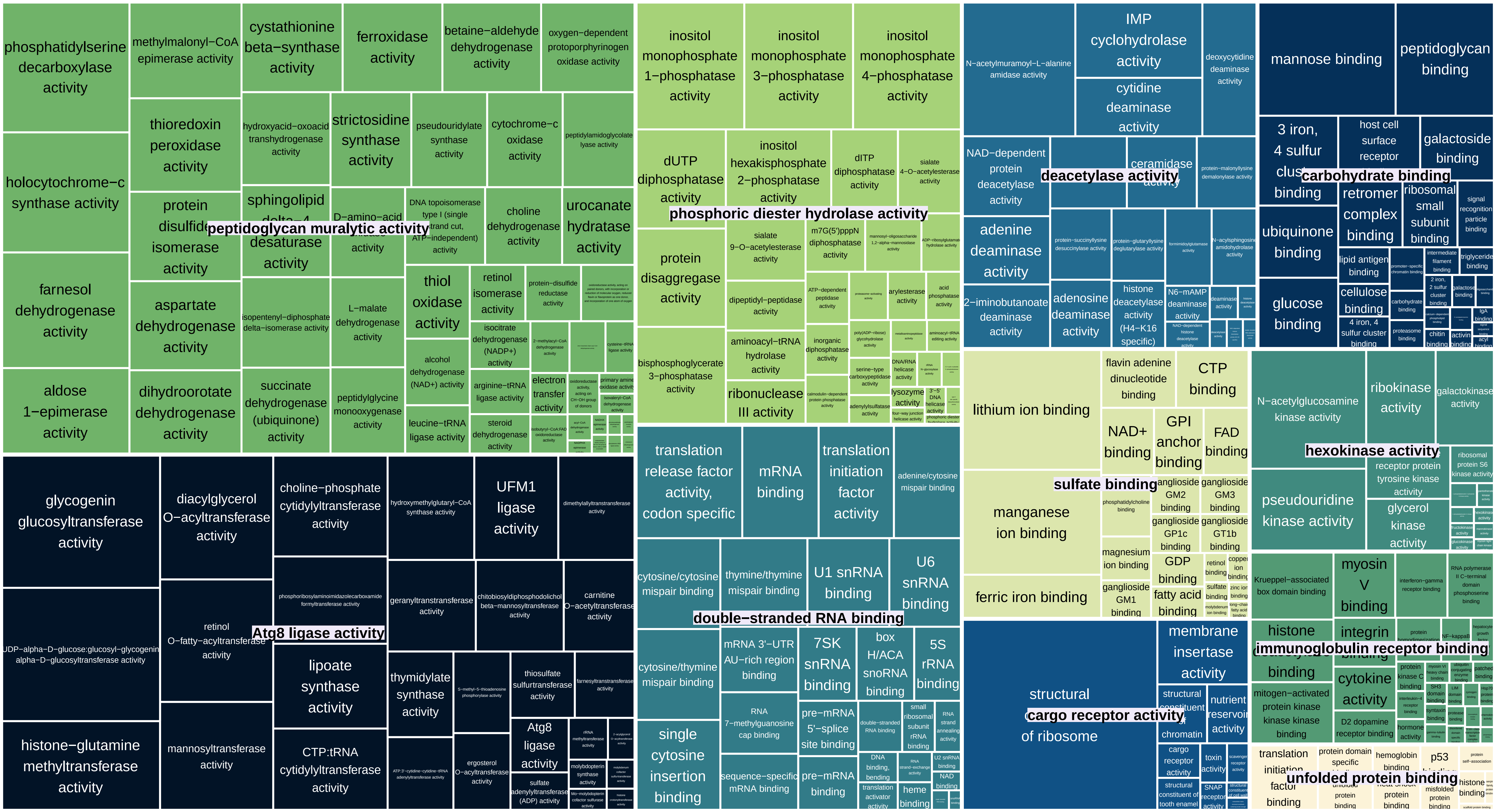

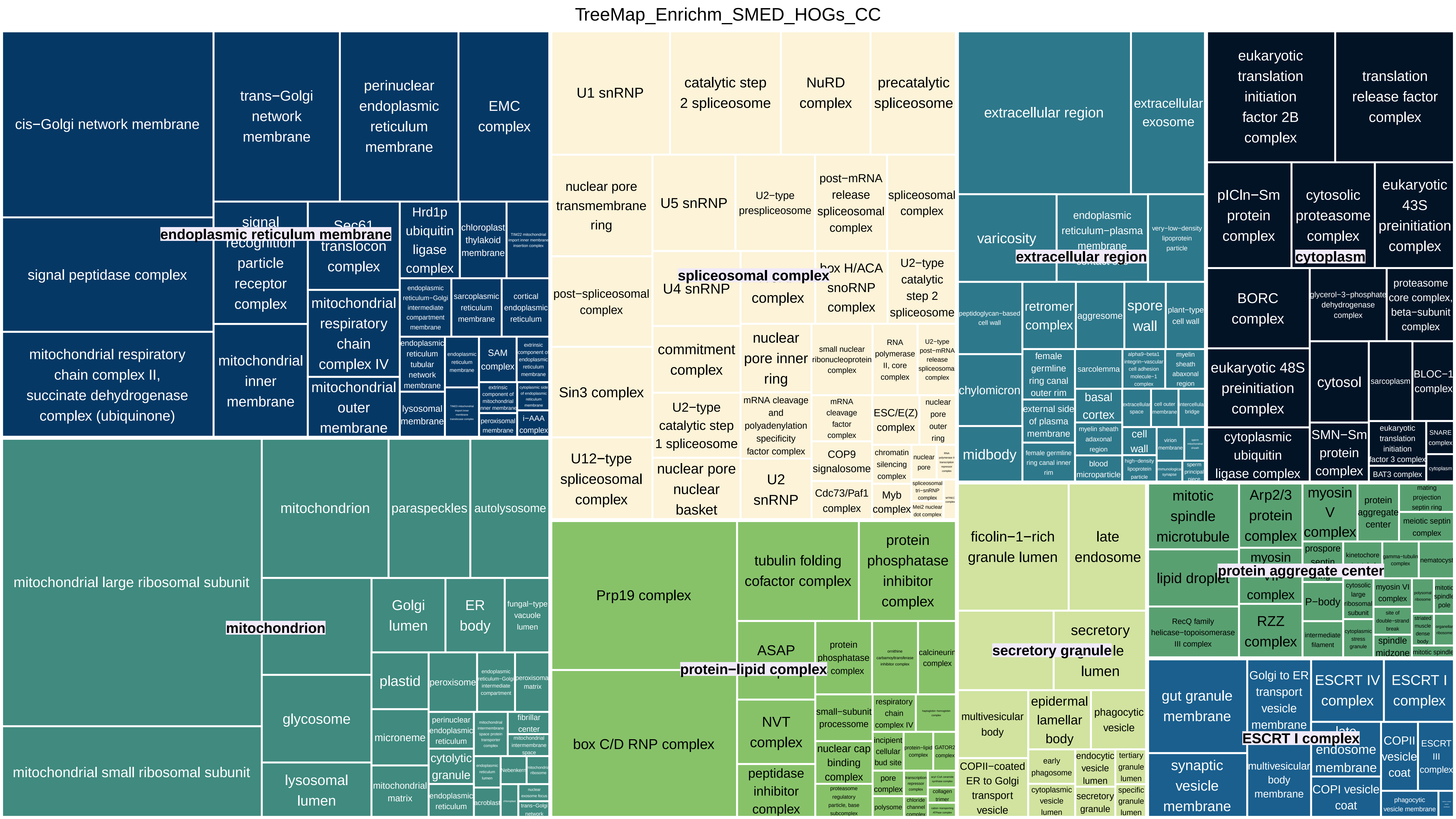
TreeMap_Enrichm_SMED_HOGs_CC
endoplasmic reticulum membrane
cis−Golgi network membrane
trans−Golgi network membrane
perinuclear endoplasmic reticulum membrane
EMC complex
signal peptidase complex
signal recognition particle receptor complex
Sec61 translocon complex
Hrd1p ubiquitin ligase complex
chloroplast thylakoid membrane
mitochondrial respiratory chain complex II, succinate dehydrogenase complex (ubiquinone)
mitochondrial inner membrane
mitochondrial respiratory chain complex IV
mitochondrial outer membrane
lysosomal membrane
SAM complex
i−AAA complex
mitochondrion
mitochondrial large ribosomal subunit
mitochondrial small ribosomal subunit
paraspeckles
autolysosome
Golgi lumen
ER body
fungal−type vacuole lumen
glycosome
plastid
peroxisome
microneme
lysosomal lumen
cytolytic granule
mitochondrial matrix
spliceosomal complex
U1 snRNP
catalytic step 2 spliceosome
NuRD complex
precatalytic spliceosome
nuclear pore transmembrane ring
U5 snRNP
U2−type prespliceosome
post−mRNA release spliceosomal complex
post−spliceosomal complex
U4 snRNP
box H/ACA snoRNP complex
U2−type catalytic step 2 spliceosome
Sin3 complex
commitment complex
nuclear pore inner ring
U12−type spliceosomal complex
U2−type catalytic step 1 spliceosome
nuclear pore nuclear basket
U2 snRNP
COP9 signalosome
Cdc73/Paf1 complex
ESC/E(Z) complex
Myb complex
protein−lipid complex
Prp19 complex
tubulin folding cofactor complex
protein phosphatase inhibitor complex
ASAP complex
box C/D RNP complex
NVT complex
peptidase inhibitor complex
nuclear cap binding complex
extracellular region
extracellular exosome
varicosity
retromer complex
aggresome
spore wall
chylomicron
midbody
basal cortex
sarcolemma
cell wall
cytoplasm
eukaryotic translation initiation factor 2B complex
translation release factor complex
pICln−Sm protein complex
cytosolic proteasome complex
eukaryotic 43S preinitiation complex
BORC complex
eukaryotic 48S preinitiation complex
cytosol
cytoplasmic ubiquitin ligase complex
SMN−Sm protein complex
protein aggregate center
ficolin−1−rich granule lumen
late endosome
mitotic spindle microtubule
Arp2/3 protein complex
myosin V complex
lipid droplet
RZZ complex
secretory granule
secretory granule lumen
multivesicular body
epidermal lamellar body
phagocytic vesicle
COPII−coated ER to Golgi transport vesicle
ESCRT I complex
gut granule membrane
Golgi to ER transport vesicle membrane
ESCRT IV complex
synaptic vesicle membrane
multivesicular body membrane
COPI vesicle coat
COPII vesicle coat
ESCRT III complex

TreeMap_Enrichm_SHARED_HOGs_BP

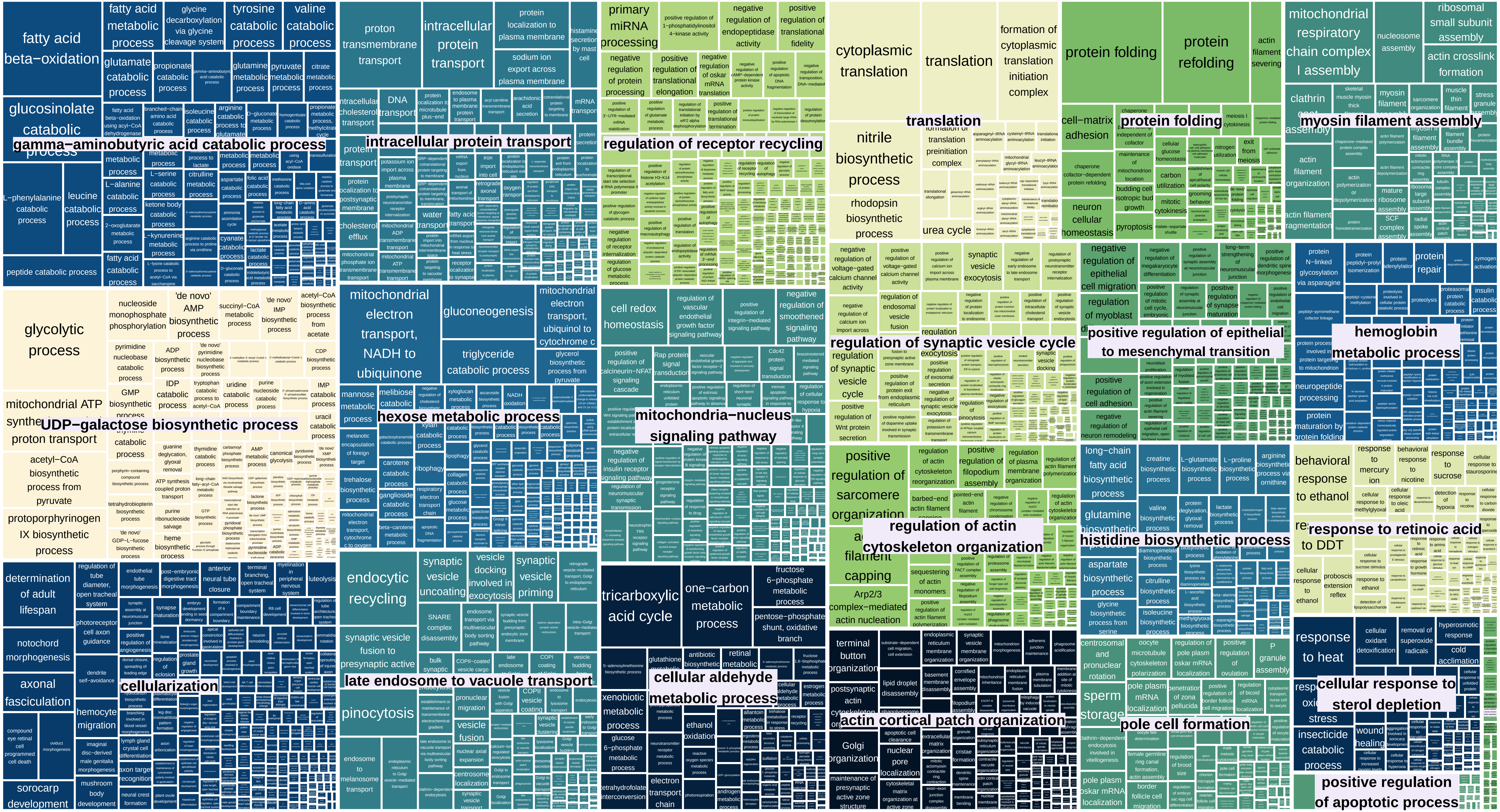

TreeMap_Enrichm_SHARED_HOGs_MF

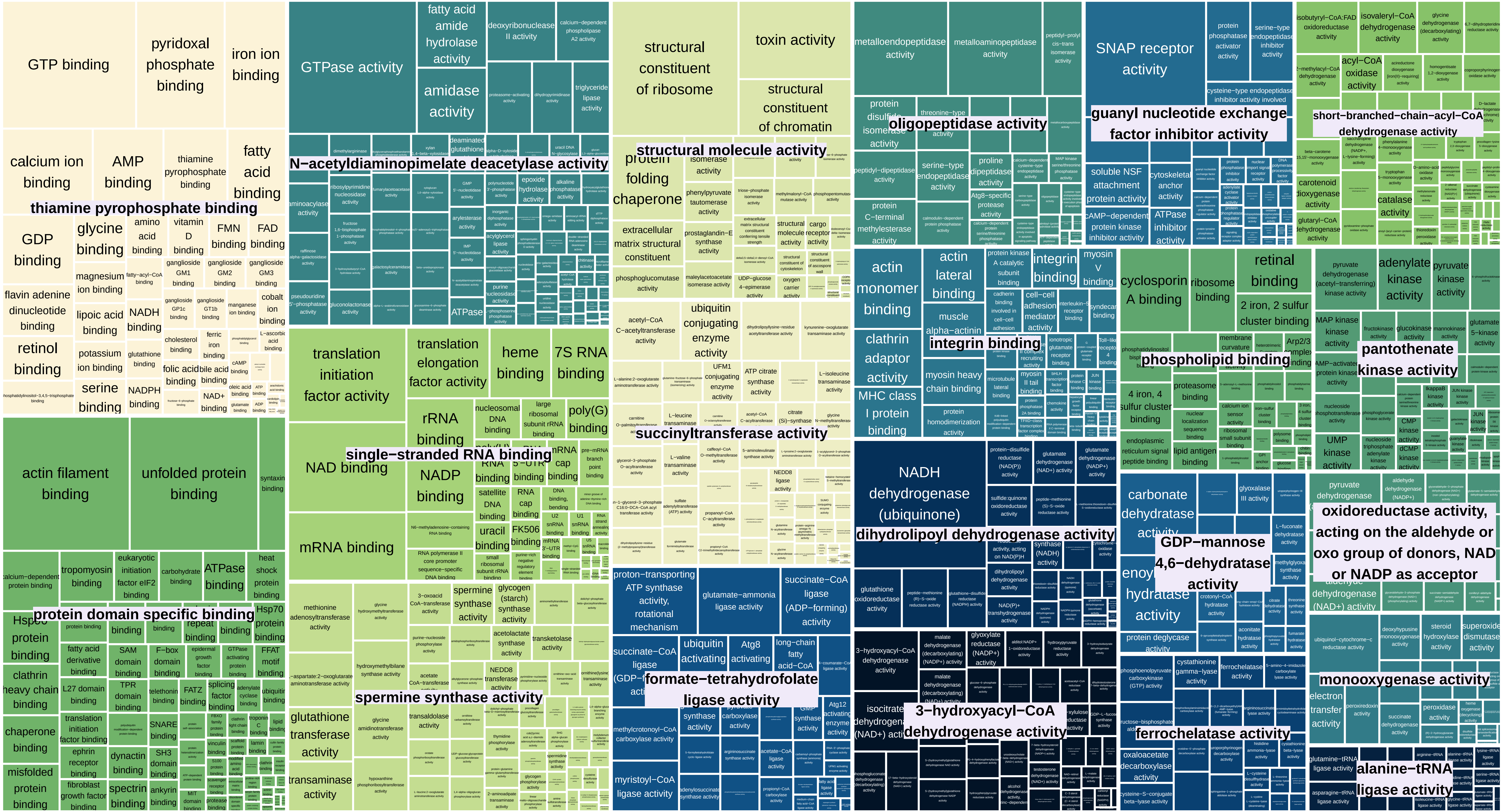

TreeMap_Enrichm_SHARED_HOGs_CC

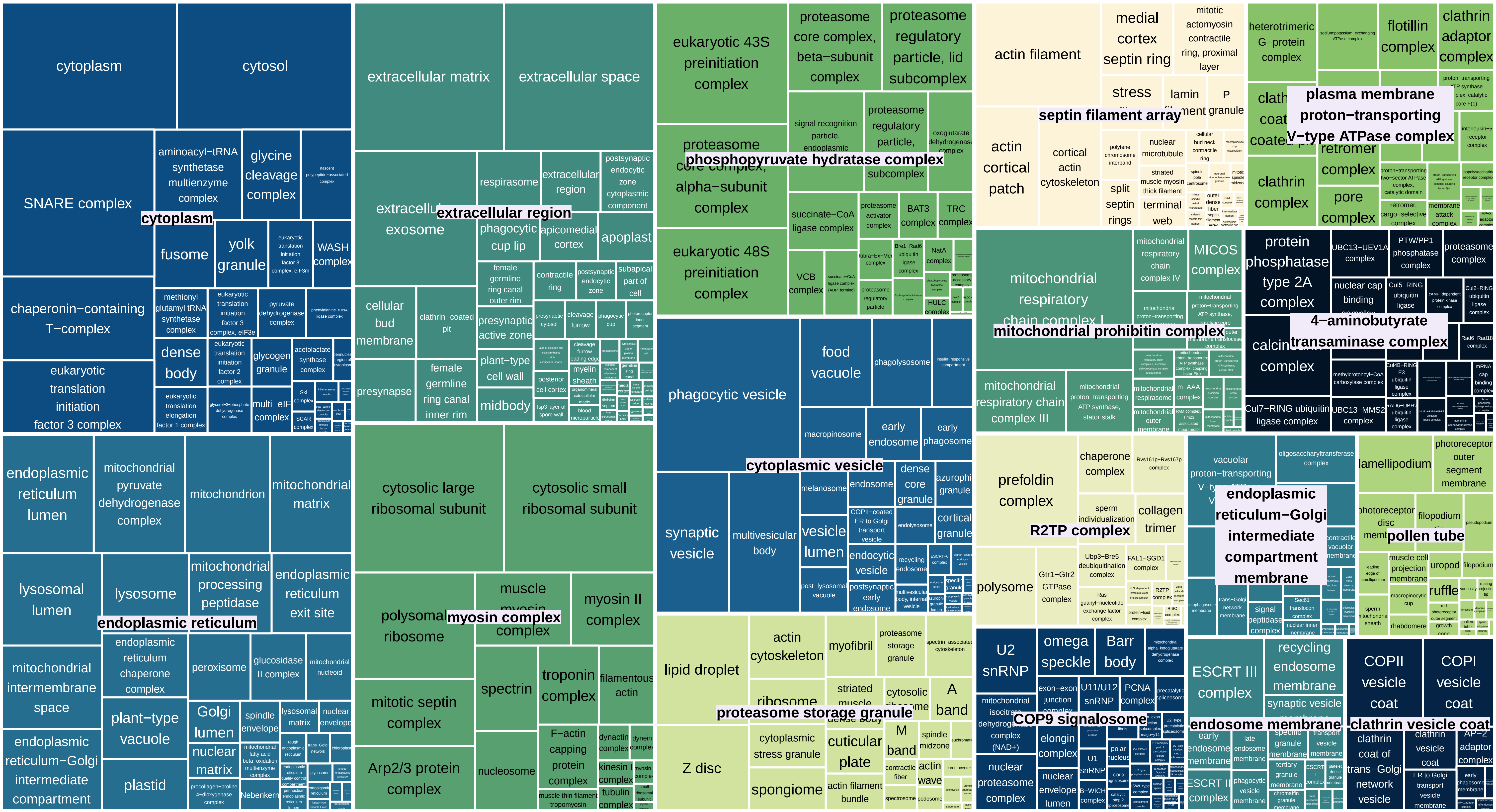
