## Supplementary Material 3 for "Genomic exaptation and regulatory landscape shifts as key mechanisms enabling flatworm terrestrialization"

**Supplementary Material 4.** Enrichments Differentially Expressed Genes (DGEs). Treemaps showing enriched GO terms (BP: Biological Processes, MF: Molecular Functions, CC: Cellular Components) in HOGs including DGEs detected in ONUN, DGEs detected in SMED, and DGEs detencted in both species (shared HOGs)

TreeMap_Enrichm_ONUN_HOGs_BP

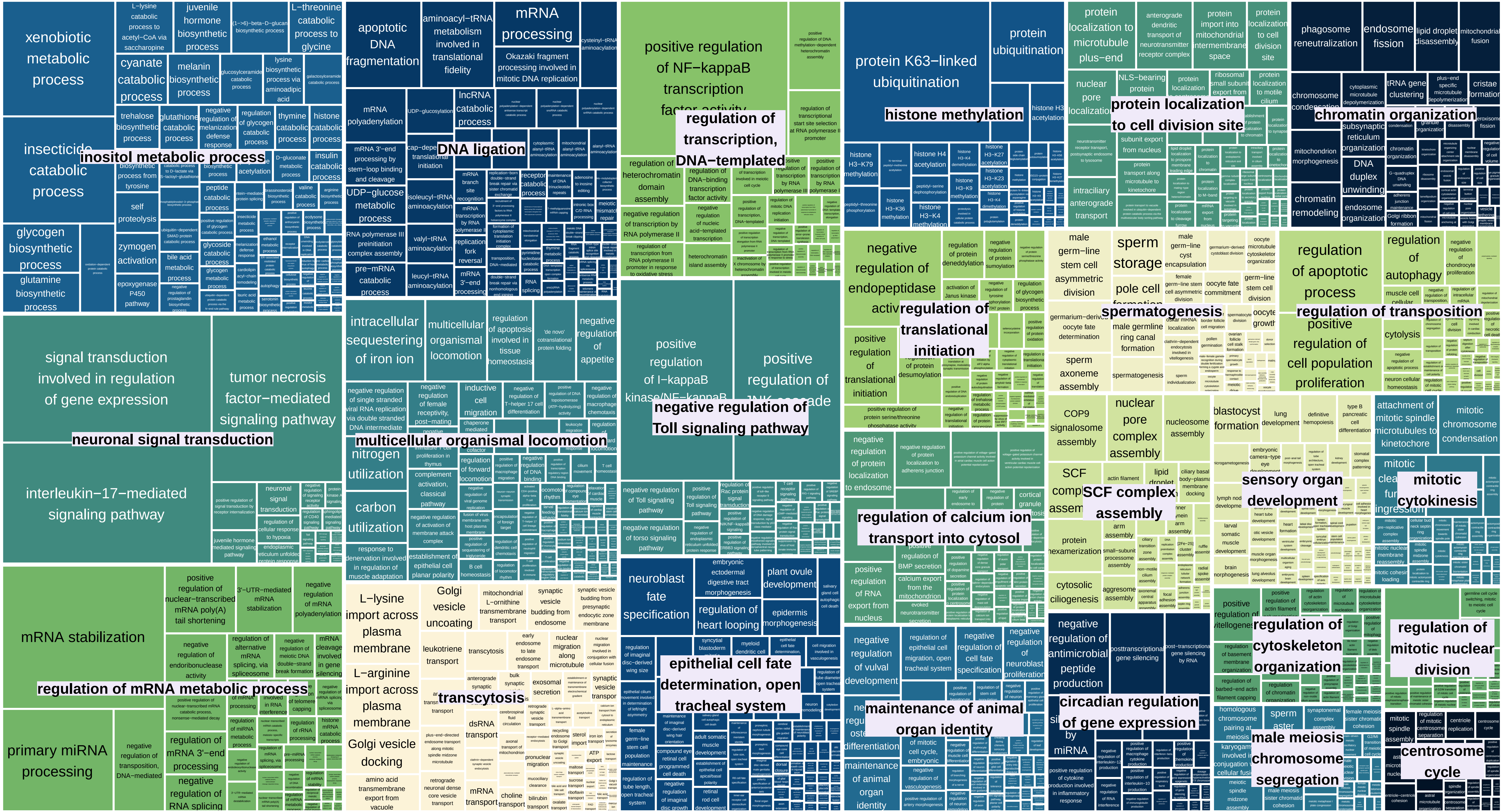

TreeMap_Enrichm_ONUN_HOGs_MF

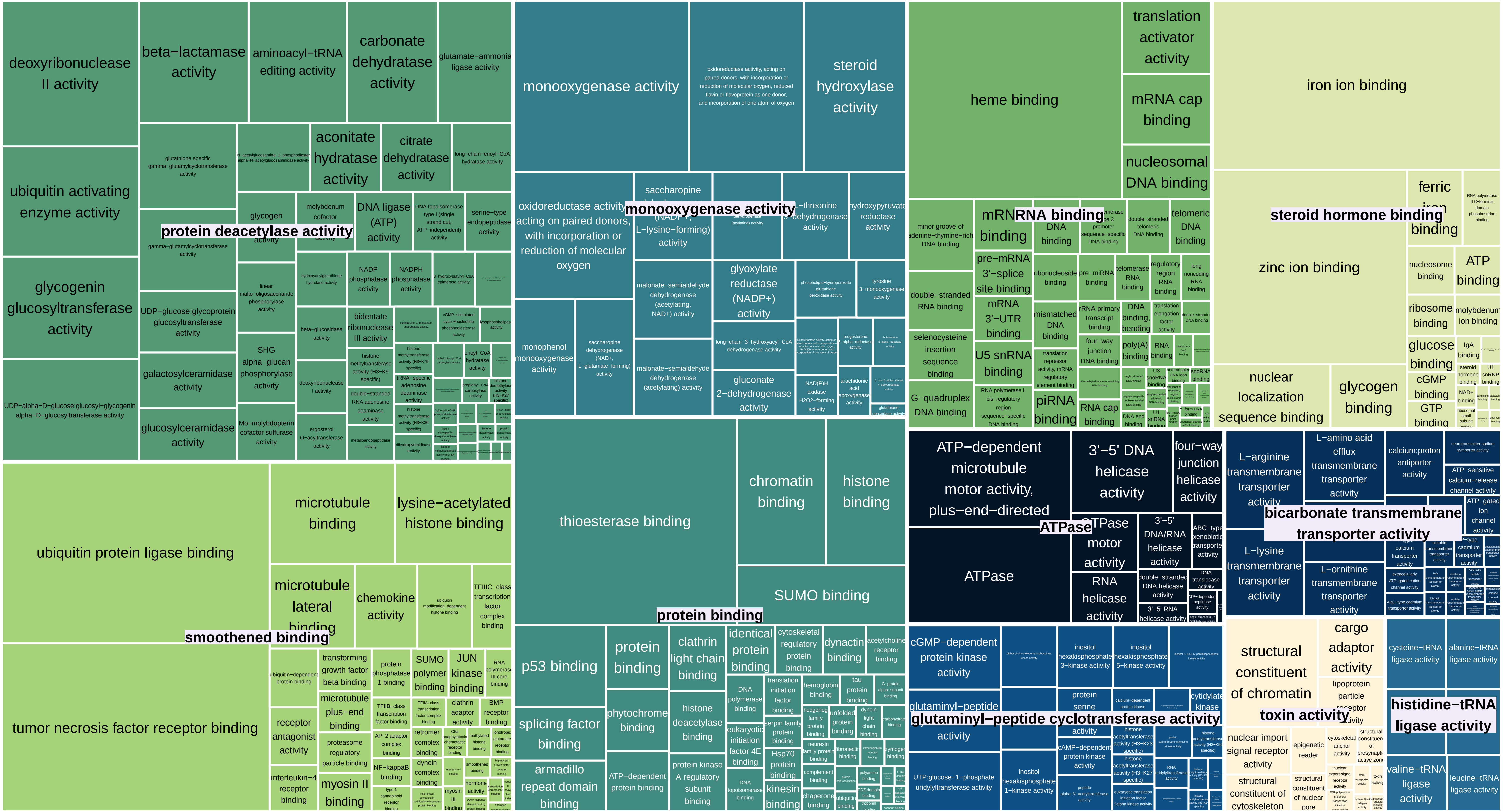

TreeMap_Enrichm_ONUN_HOGs_CC

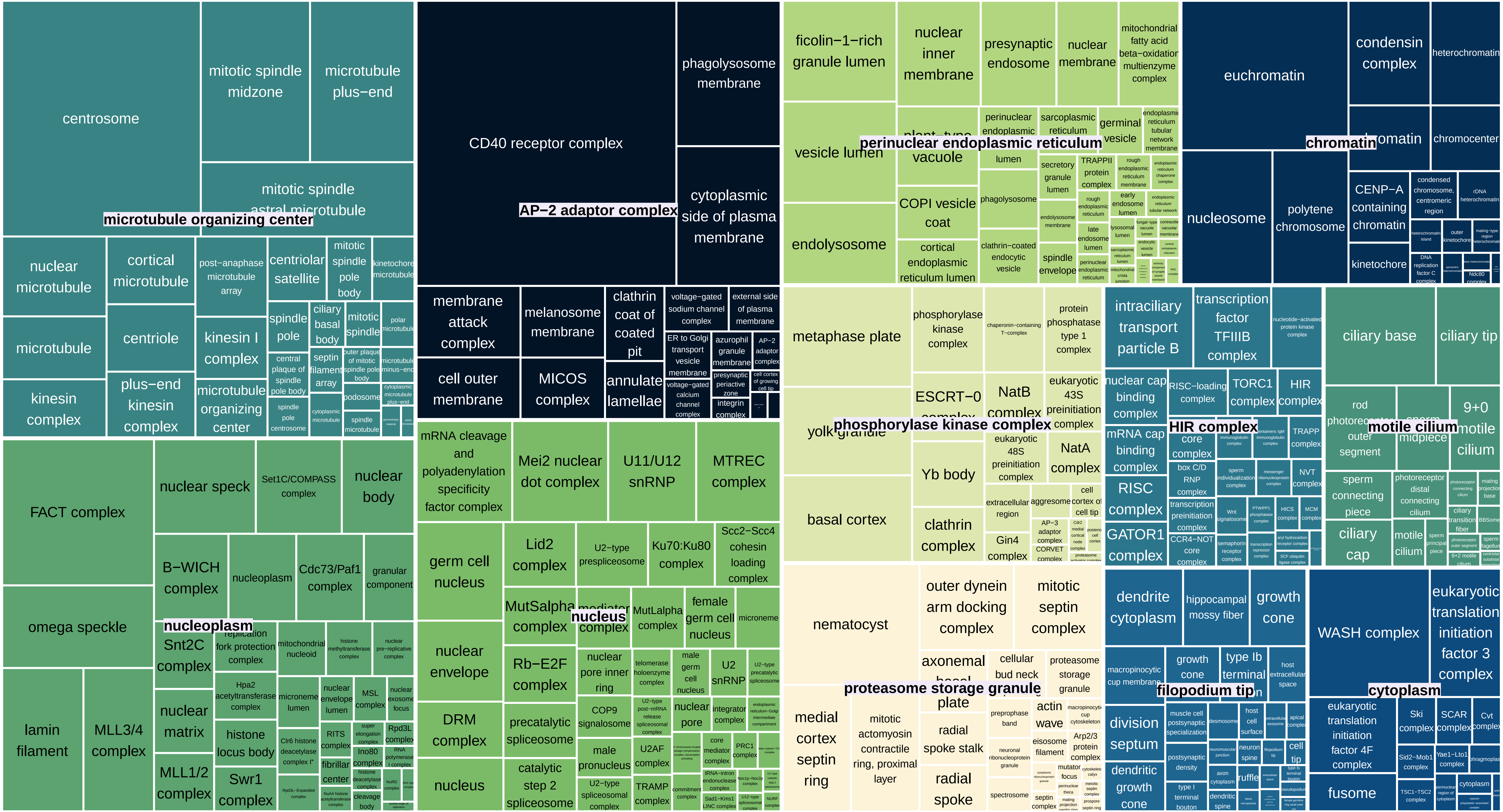

TreeMap_Enrichm_SMED_HOGs_BP

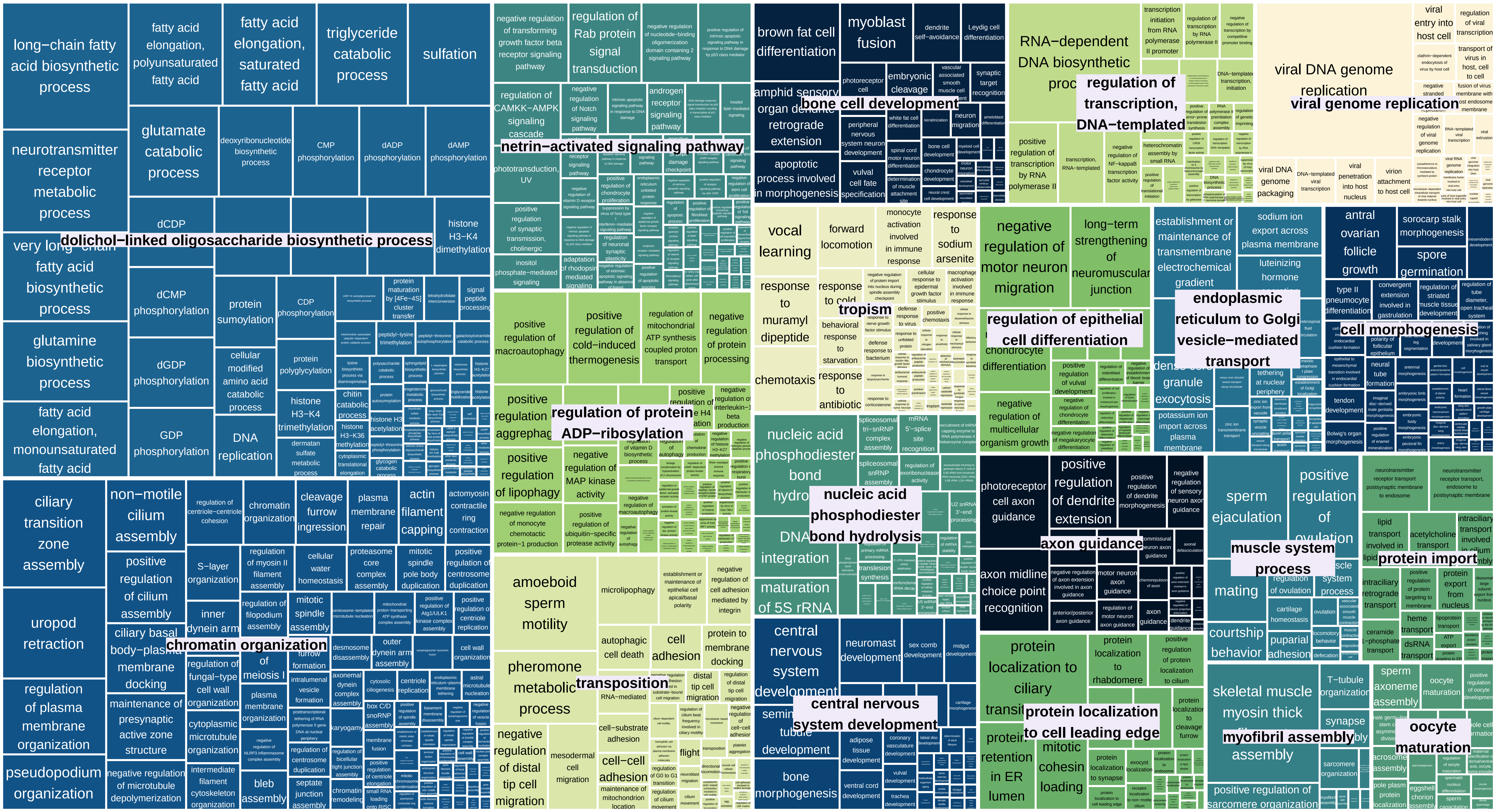

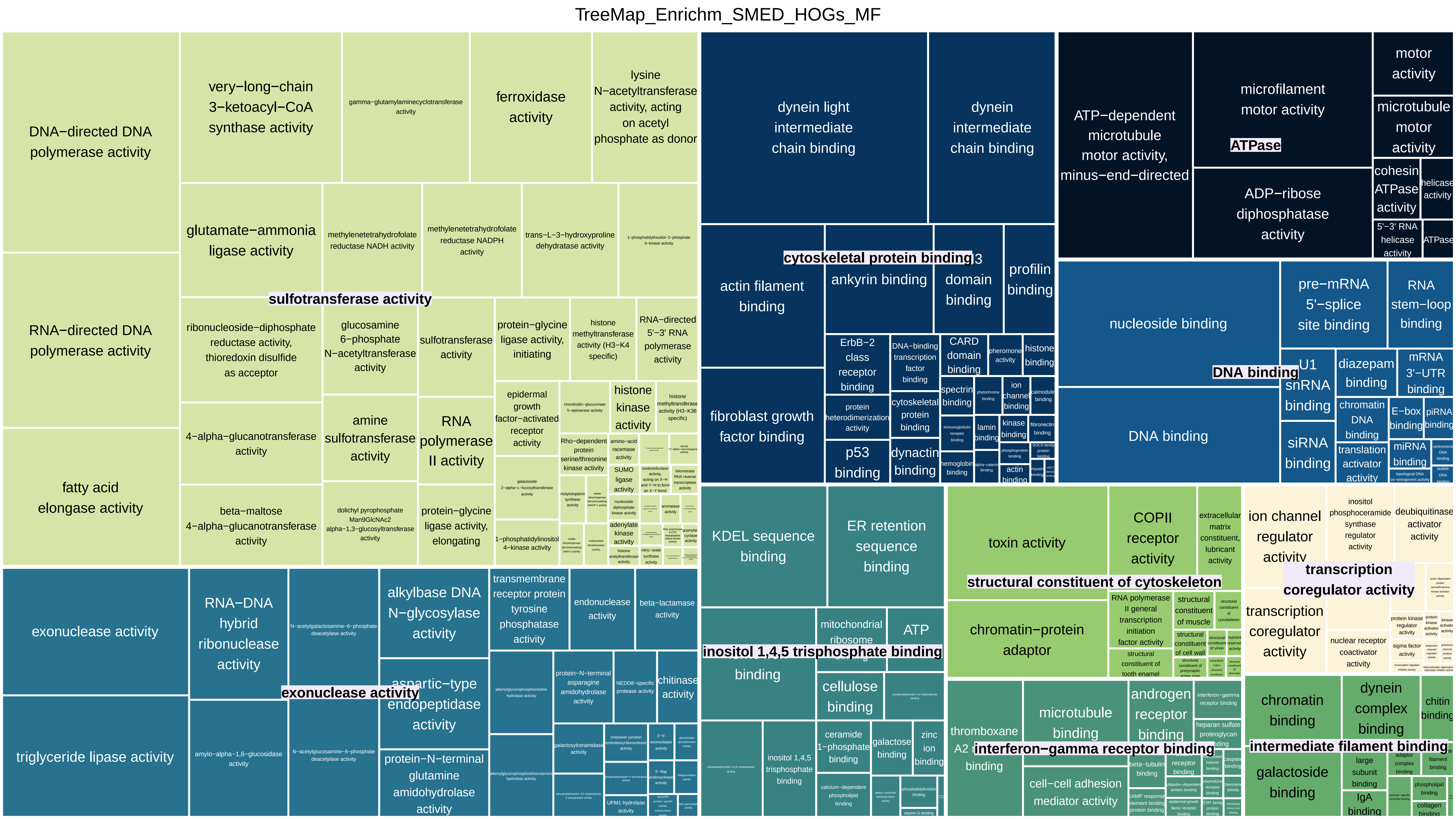
TreeMap_Enrichm_SMED_HOGs_MF
DNA−directed DNA polymerase activity
RNA−directed DNA polymerase activity
fatty acid elongase activity
very−long−chain 3−ketoacyl−CoA synthase activity
gamma−glutamylaminecyclotransferase activity
ferroxidase activity
lysine N−acetyltransferase activity, acting on acetyl phosphate as donor
glutamate−ammonia ligase activity
methylenetetrahydrofolate reductase NADH activity
methylenetetrahydrofolate reductase NADPH activity
trans−L−3−hydroxyproline dehydratase activity
1−phosphatidylinositol−3−phosphate 5−kinase activity
sulfotransferase activity
ribonucleoside−diphosphate reductase activity, thioredoxin disulfide as acceptor
glucosamine 6−phosphate N−acetyltransferase activity
sulfotransferase activity
protein−glycine ligase activity, initiating
histone methyltransferase activity (H3−K4 specific)
RNA−directed 5'−3' RNA polymerase activity
4−alpha−glucanotransferase activity
amine sulfotransferase activity
RNA polymerase II activity
epidermal growth factor−activated receptor activity
histone kinase activity
beta−maltose 4−alpha−glucanotransferase activity
dolichyl pyrophosphate Man9GlcNAc2 alpha−1,3−glucosyltransferase activity
protein−glycine ligase activity, elongating
1−phosphatidylinositol 4−kinase activity
exonuclease activity
triglyceride lipase activity
RNA−DNA hybrid ribonuclease activity
amylo−alpha−1,6−glucosidase activity
N−acetylgalactosamine−6−phosphate deacetylase activity
N−acetylglucosamine−6−phosphate deacetylase activity
alkylbase DNA N−glycosylase activity
aspartic−type endopeptidase activity
protein−N−terminal glutamine amidohydrolase activity
transmembrane receptor protein tyrosine phosphatase activity
endonuclease activity
beta−lactamase activity
protein−N−terminal asparagine amidohydrolase activity
chitinase activity
dynein light intermediate chain binding
dynein intermediate chain binding
cytoskeletal protein binding
actin filament binding
ankyrin binding
SH3 domain binding
profilin binding
fibroblast growth factor binding
ErbB−2 class receptor binding
DNA−binding transcription factor binding
CARD domain binding
histone binding
protein heterodimerization activity
cytoskeletal protein binding
p53 binding
dynactin binding
spectrin binding
ATP−dependent microtubule motor activity, minus−end−directed
microfilament motor activity
ATPase
ADP−ribose diphosphatase activity
motor activity
microtubule motor activity
cohesin ATPase activity
helicase activity
5'−3' RNA helicase activity
ATPase
nucleoside binding
DNA binding
DNA binding
pre−mRNA 5'−splice site binding
RNA stem−loop binding
U1 snRNA binding
diazepam binding
mRNA 3'−UTR binding
chromatin DNA binding
E−box binding
piRNA binding
siRNA binding
translation activator activity
miRNA binding
KDEL sequence binding
ER retention sequence binding
mitochondrial ribosome binding
ATP binding
inositol 1,4,5 trisphosphate binding
cellulose binding
inositol 1,4,5 trisphosphate binding
ceramide 1−phosphate binding
galactose binding
zinc ion binding
toxin activity
COPII receptor activity
extracellular matrix constituent, lubricant activity
structural constituent of cytoskeleton
chromatin−protein adaptor
RNA polymerase II general transcription initiation factor activity
structural constituent of muscle
structural constituent of tooth enamel
thromboxane A2 receptor binding
microtubule binding
androgen receptor binding
interferon−gamma receptor binding
cell−cell adhesion mediator activity
ion channel regulator activity
inositol phosphoceramide synthase regulator activity
deubiquitinase activator activity
transcription coregulator activity
transcription coregulator activity
nuclear receptor coactivator activity
chromatin binding
dynein complex binding
chitin binding
intermediate filament binding
galactoside binding
large subunit binding
IgA binding

TreeMap_Enrichm_SMED_HOGs_CC

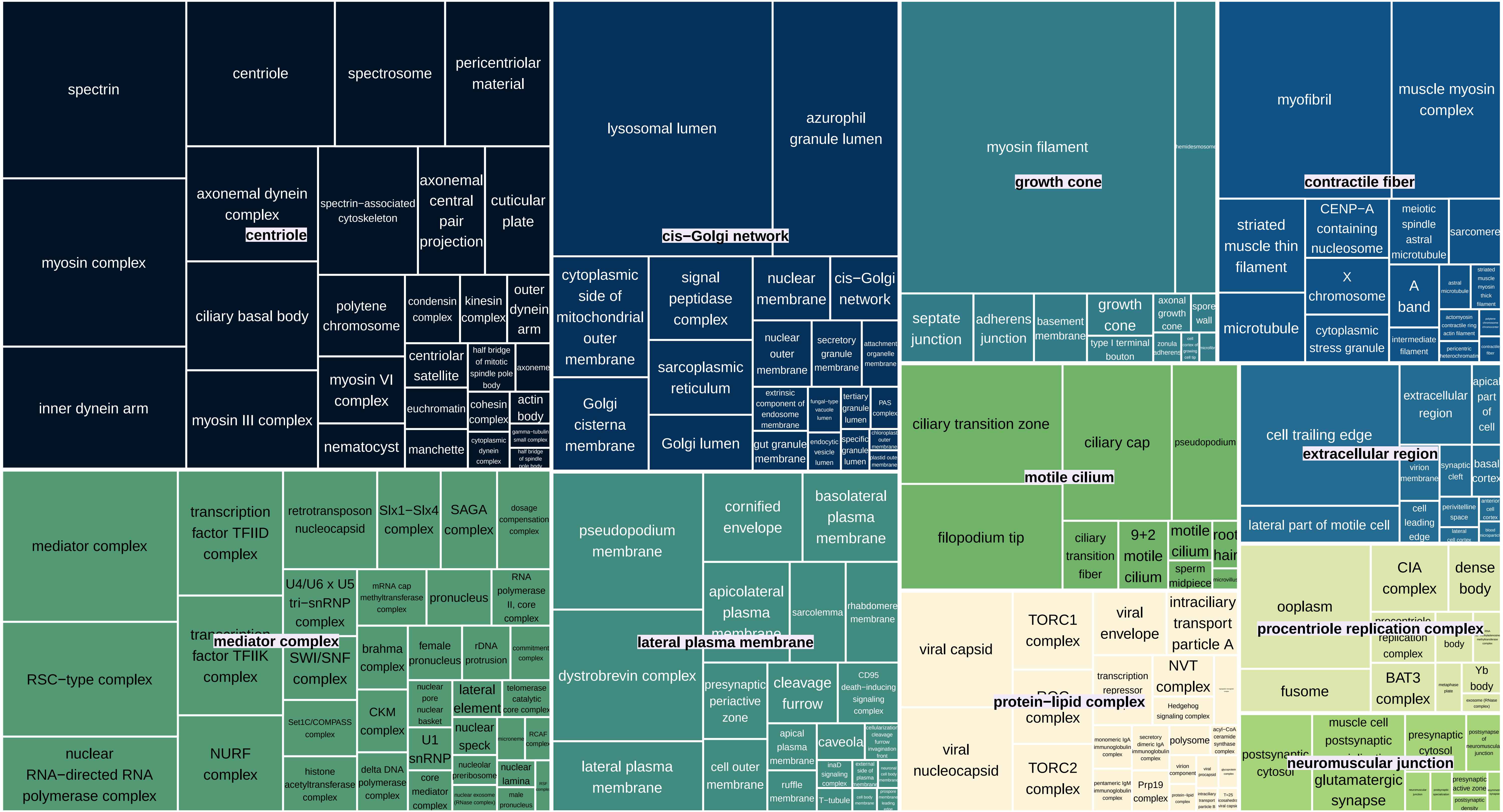

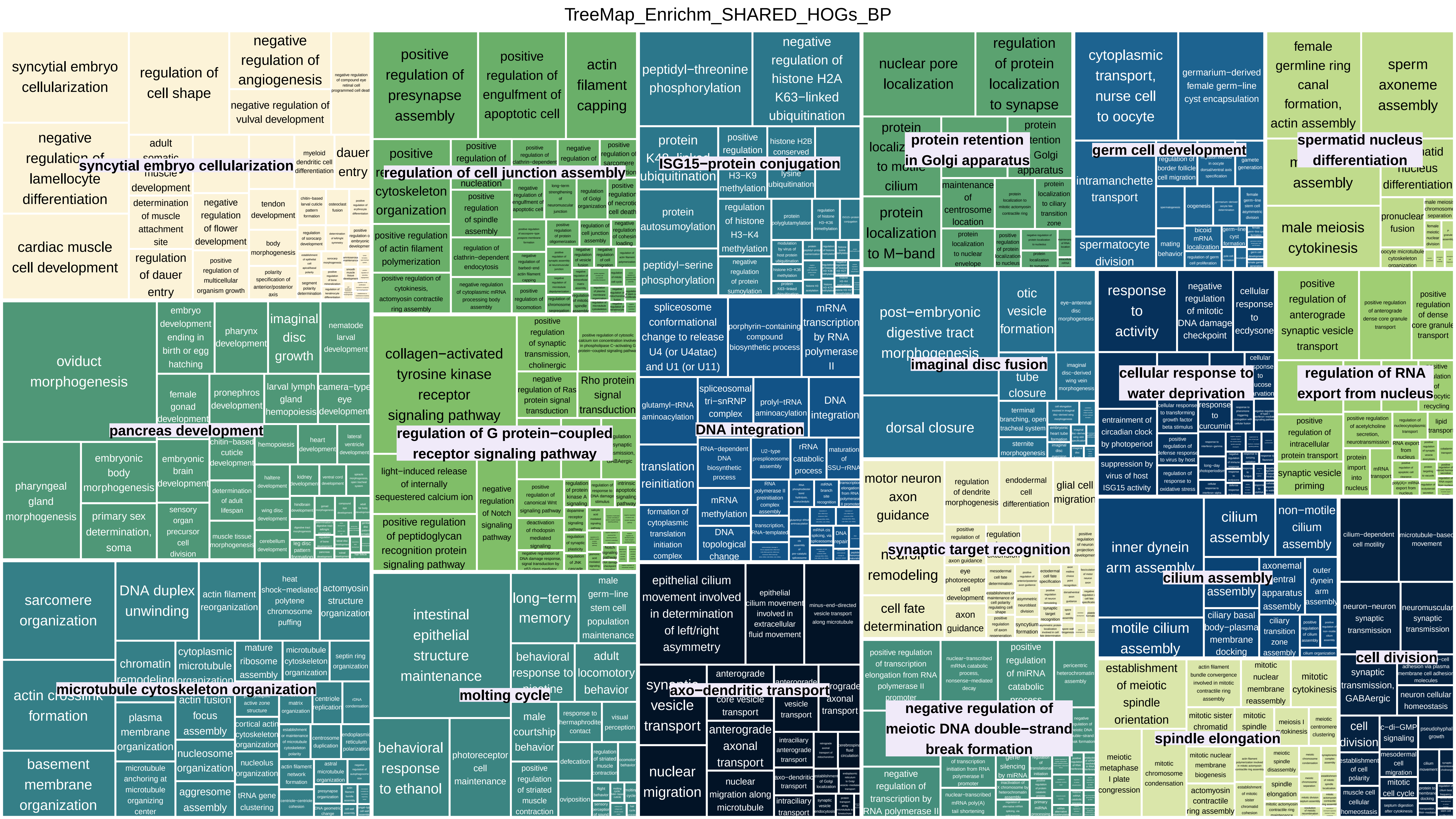

TreeMap_Enrichm_SHARED_HOGs_BP
syncytial embryo cellularization
regulation of cell junction assembly
ISG15−protein conjugation
protein retention in Golgi apparatus
germ cell development
spermatid nucleus differentiation
pancreas development
regulation of G protein−coupled receptor signaling pathway
DNA integration
imaginal disc fusion
cellular response to water deprivation
regulation of RNA export from nucleus
synaptic target recognition
cilium assembly
microtubule cytoskeleton organization
molting cycle
axo−dendritic transport
negative regulation of meiotic DNA double−strand break formation
spindle elongation
cell division

TreeMap_Enrichm_SHARED_HOGs_MF

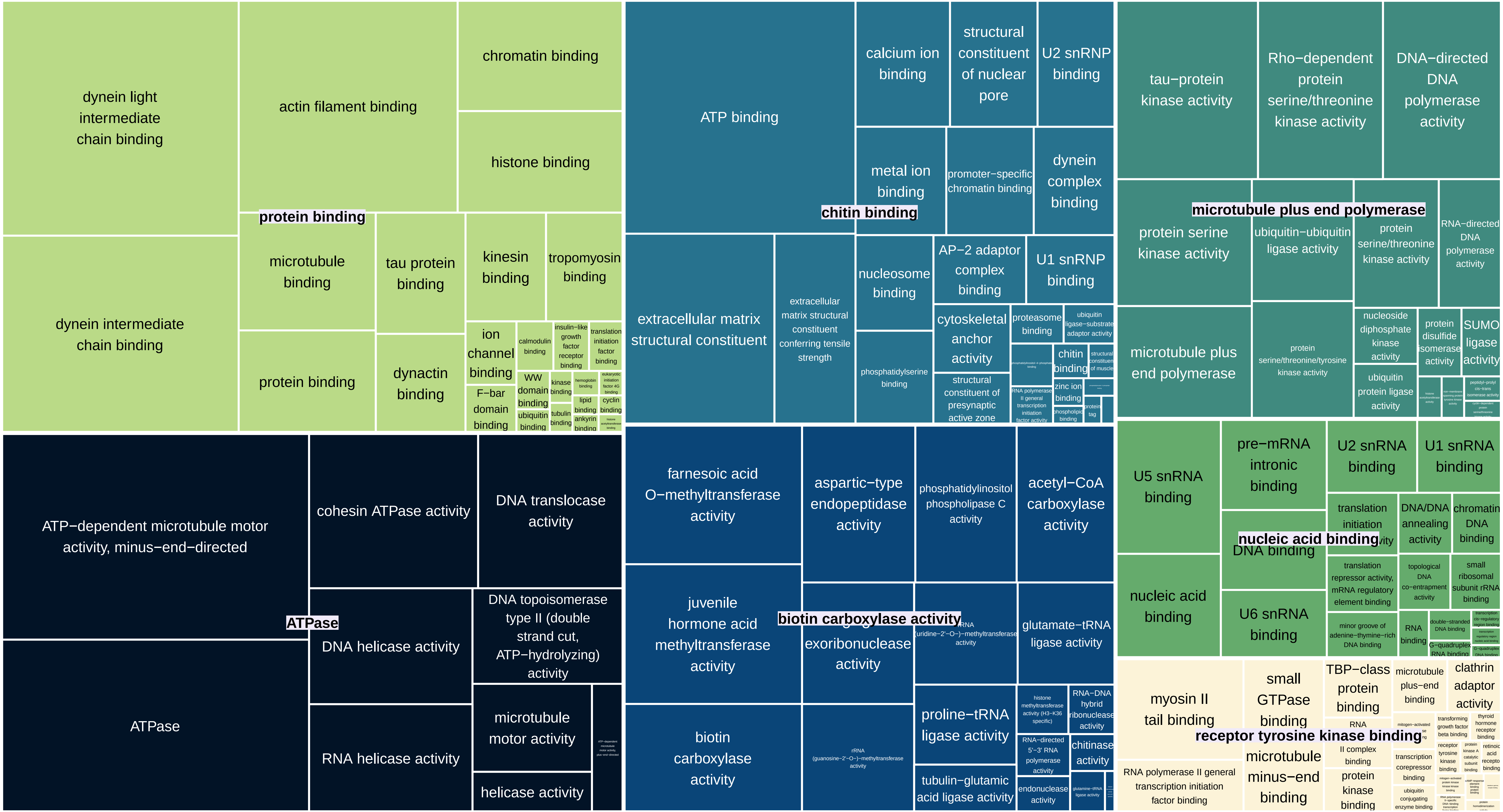

TreeMap_Enrichm_SHARED_HOGs_CC

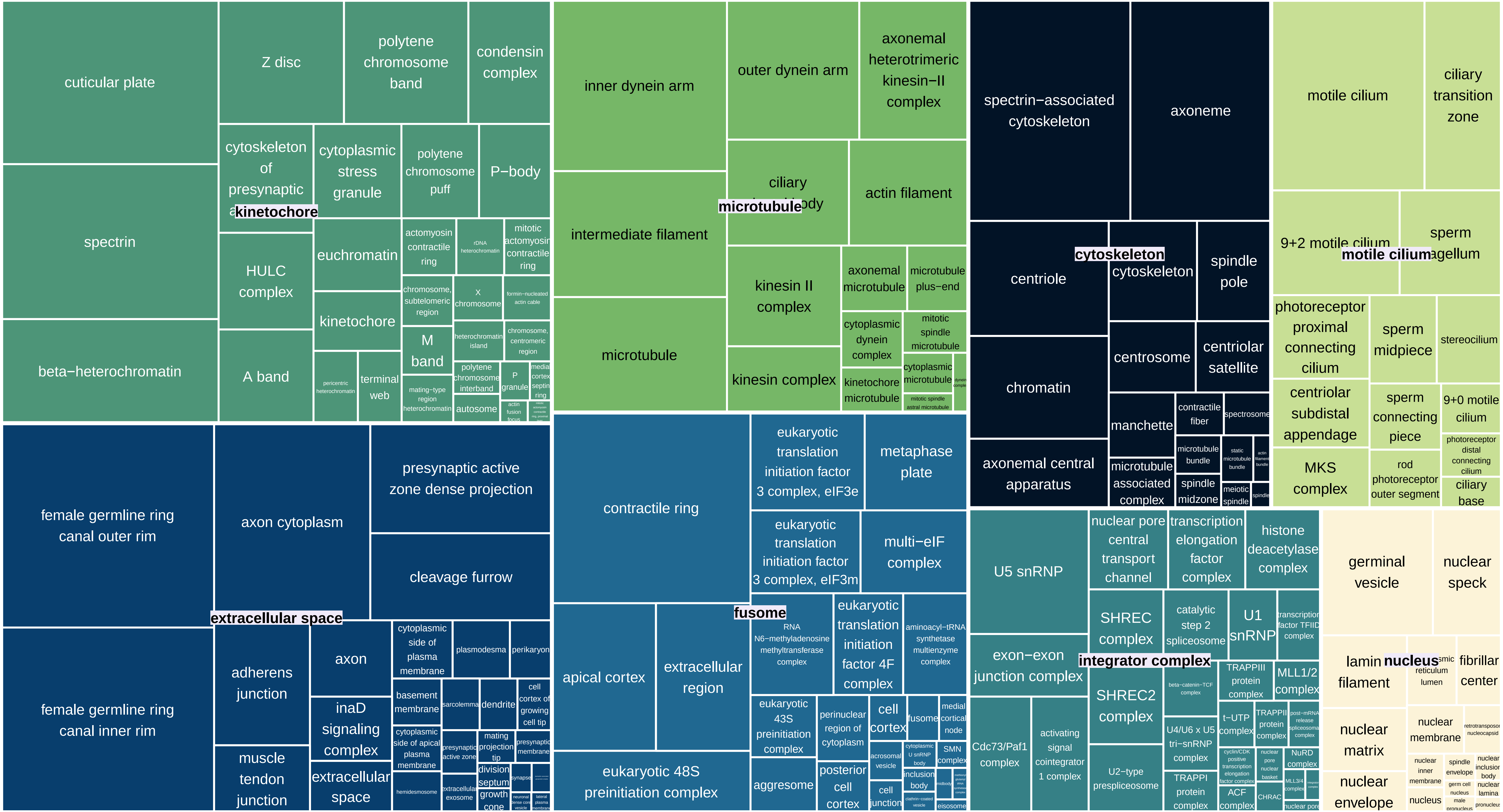
