## Supplementary Material 4 for "Genomic exaptation and regulatory landscape shifts as key mechanisms enabling flatworm terrestrialization"

**Supplementary Material 5.** Functional characterization of HOGs including differentially expressed genes from *O.nungara* (ONUN) and *S. mediterranea* (SMED) shared between different experiments. A) Comparisons between experiments and between species in different experiments. Blue: comparison between species in the same experiment and the same regulation do not share HOGs. expA has been used to represent all experimental conditions. Green: comparisons sharing a high number of HOGs. Pink: comparisons sharing a low number of HOGs. B) Functional characterization of shared HOGs. Up: a simplified treemap of the enriched functions in all shared HOGs. Only the most general GOs in the squares are indicated. Some enriched GOs that can be related to adaptation to life on land are indicated out of the treemap. Down (left): Identified pathways were these genes are involved. Down (rigth): General functions of HOGs that can act as regulatory elements

**A**

| | |
|---|---|
| expA_UP_ONUN-expA_UP_SMED | 0 |
| expA_Down_ONUN-expA_Down_SMED | 0 |

| | |
|---|---|
| ONUN_Hyperoxia-**SMED_Osmo** | 122 |
| ONUN_Hypoxia-**SMED_Osmo** | 102 |
| ONUN_VL-**SMED_Osmo** | 80 |
| ONUN_Osmo-**SMED_Osmo** | 13 |

| | |
|---|---|
| ONUN_Hyperoxia-SMED_chemoDeath | 4 |
| ONUN_Hyperoxia-SMED_chemoFood | 3 |
| ONUN_Hyperoxia-SMED_UV24D | 3 |
| ONUN_Hypoxia-SMED_chemoDeath | 3 |
| ONUN_Hypoxia-SMED_UV24D | 3 |
| ONUN_chemoFood-**SMED_Osmo** | 2 |
| ONUN_Hyperoxia-SMED_Hypoxia | 2 |
| ONUN_Hyperoxia-SMED_VL | 2 |
| ONUN_chemoDeath-**SMED_Osmo** | 1 |
| ONUN_Hypoxia-SMED_chemoFood | 1 |

**B**

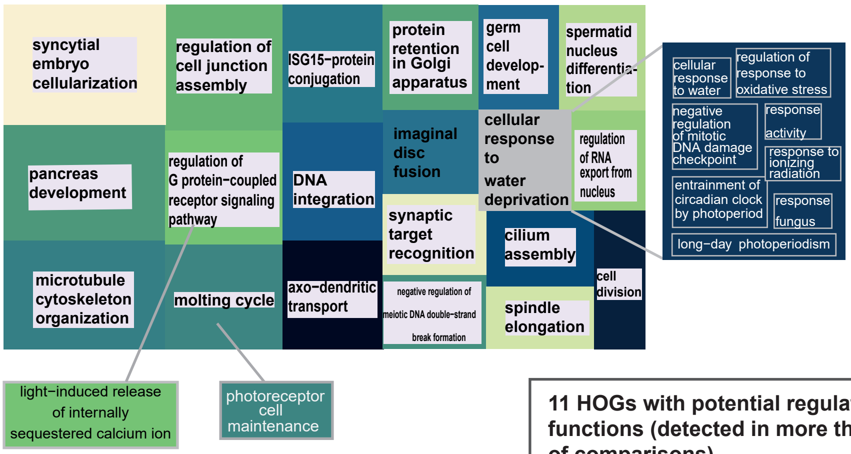


**125 genes involved in pathways related to:**

Metabolism of RNA
Signal Transduction
Metabolism
Cell Cycle
Organelle biogenesis and maintenance
Disease
Gene expression (Transcription)

**11 HOGs with potential regulatory functions (detected in more than 10% of comparisons)**

protein modifications
mRNA splicing
regulation of apoptosis
cellular response to water deprivation
methylation
signal transduction
DNA and RNA processes
