## Supplementary Material 5 for "Genomic exaptation and regulatory landscape shifts as key mechanisms enabling flatworm terrestrialization"

**Supplementary Material 7.** Functional characterization of Differentially Expressed Genes detected by proteomics (DGEs-prot) in *Obama nungara* (ONUN) and *Schmidtea mediterranea* (SMED). Treemaps showing enriched GO terms related to Biological Processes (BP). Extract of Reactome report for genes annotated with eggnog-mapper. The full Reactome reports and the Blast output are available at (see links below)

TreeMap_Enrichm_Validated_DGE_prot_ONUN_BP

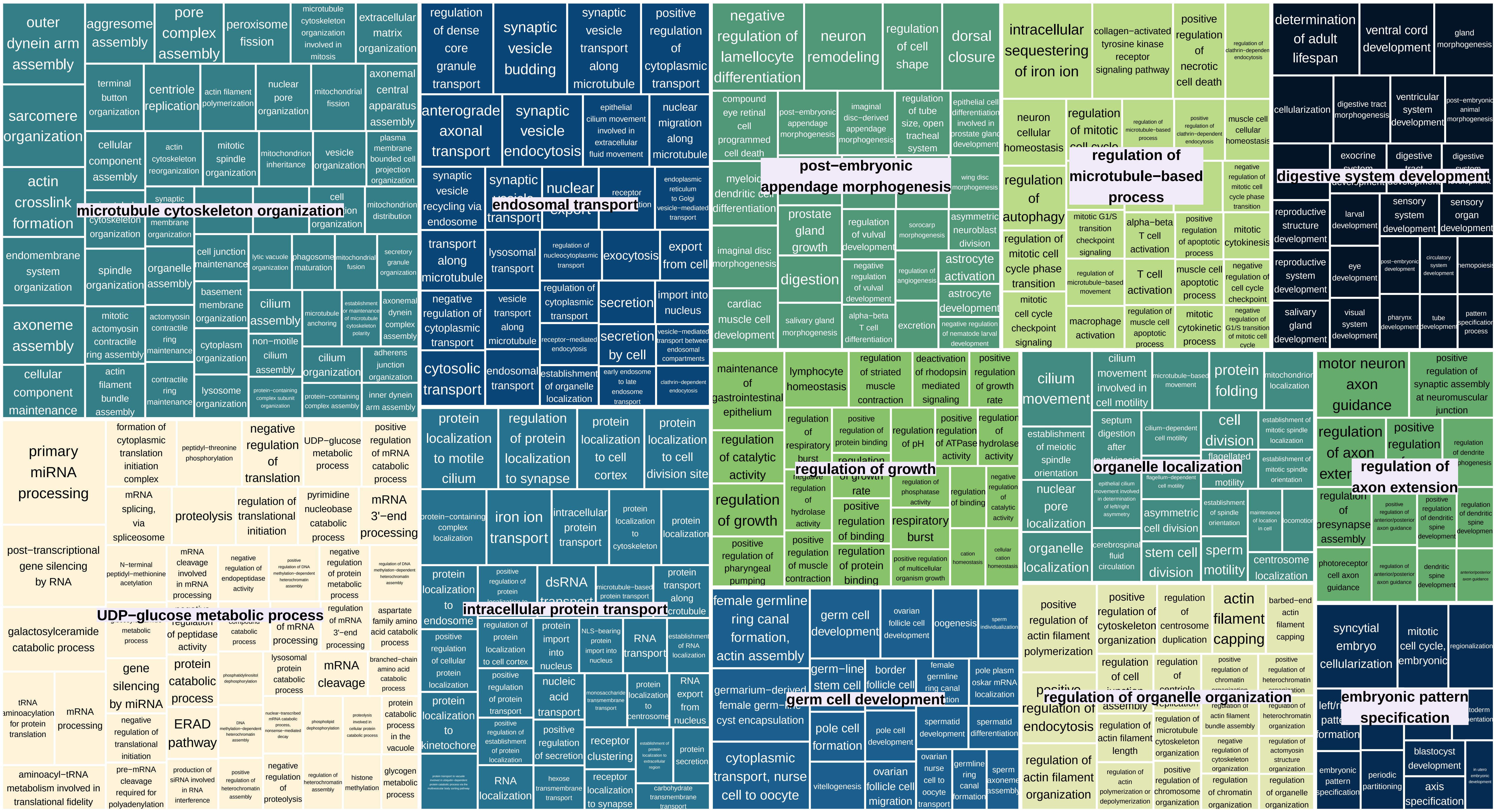

### Most significant pathways DGEs-prot ONUN

link to full report: poner link en github

The following table shows the 25 most relevant pathways sorted by p-value.

| Pathway name | Entities | | | | Reactions | |
|---|---|---|---|---|---|---|
| | found | ratio | p-value | FDR* | found | ratio |
| MHC class II antigen presentation | 19 / 199 | 0.009 | 1.50e-06 | 0.002 | 15 / 26 | 0.002 |
| Cytosolic tRNA aminoacylation | 7 / 63 | 0.003 | 9.58e-05 | 0.071 | 13 / 21 | 0.001 |
| mRNA Splicing - Minor Pathway | 10 / 139 | 0.006 | 5.59e-04 | 0.275 | 5 / 5 | 3.36e-04 |
| RHO GTPases activate KTN1 | 3 / 12 | 5.22e-04 | 0.001 | 0.398 | 1 / 2 | 1.34e-04 |
| Clathrin-mediated endocytosis | 13 / 234 | 0.01 | 0.002 | 0.562 | 27 / 35 | 0.002 |
| Golgi-to-ER retrograde transport | 11 / 248 | 0.011 | 0.003 | 0.712 | 9 / 18 | 0.001 |
| Formation of annular gap junctions | 3 / 18 | 7.83e-04 | 0.003 | 0.712 | 2 / 2 | 1.34e-04 |
| Scavenging by Class A Receptors | 4 / 49 | 0.002 | 0.009 | 0.875 | 6 / 10 | 6.72e-04 |
| Gap junction degradation | 3 / 26 | 0.001 | 0.009 | 0.875 | 4 / 4 | 2.69e-04 |
| Retrograde neurotrophin signalling | 6 / 50 | 0.002 | 0.01 | 0.875 | 3 / 3 | 2.02e-04 |
| COPI-mediated anterograde transport | 9 / 215 | 0.009 | 0.01 | 0.875 | 10 / 12 | 8.06e-04 |
| Formation of a pool of free 40S subunits | 6 / 111 | 0.005 | 0.011 | 0.875 | 2 / 2 | 1.34e-04 |
| Prefoldin mediated transfer of substrate to CCT/TriC | 3 / 29 | 0.001 | 0.012 | 0.875 | 1 / 2 | 1.34e-04 |
| Formation of the ternary complex, and subsequently, the 43S complex | 4 / 54 | 0.002 | 0.013 | 0.875 | 1 / 3 | 2.02e-04 |
| Neutrophil degranulation | 15 / 478 | 0.021 | 0.013 | 0.875 | 6 / 10 | 6.72e-04 |
| Formation of tubulin folding intermediates by CCT/TriC | 3 / 30 | 0.001 | 0.014 | 0.875 | 2 / 2 | 1.34e-04 |
| Defective SFTPA2 causes IPF | 1 / 1 | 4.35e-05 | 0.016 | 0.875 | 1 / 1 | 6.72e-05 |
| ALK mutants bind TKIs | 2 / 12 | 5.22e-04 | 0.017 | 0.875 | 1 / 1 | 6.72e-05 |
| Interaction between L1 and Ankyrins | 3 / 33 | 0.001 | 0.017 | 0.875 | 4 / 4 | 2.69e-04 |
| L13a-mediated translational silencing of Ceruloplasmin expression | 6 / 126 | 0.005 | 0.019 | 0.875 | 2 / 3 | 2.02e-04 |
| Rev-mediated nuclear export of HIV RNA | 7 / 92 | 0.004 | 0.019 | 0.875 | 9 / 10 | 6.72e-04 |
| Glycogen metabolism | 4 / 61 | 0.003 | 0.019 | 0.875 | 10 / 37 | 0.002 |
| WNT5A-dependent internalization of FZD2, FZD5 and ROR2 | 2 / 13 | 5.65e-04 | 0.02 | 0.875 | 1 / 2 | 1.34e-04 |
| Defective TPR may confer susceptibility towards thyroid papillary carcinoma (TPC) | 3 / 36 | 0.002 | 0.022 | 0.875 | 1 / 1 | 6.72e-05 |

| Pathway name | Entities | | | | Reactions | |
|---|---|---|---|---|---|---|
| | found | ratio | p-value | FDR* | found | ratio |
| Depolymerization of the Nuclear Lamina | 3 / 36 | 0.002 | 0.022 | 0.875 | 3 / 6 | 4.03e-04 |

* False Discovery Rate

TreeMap_Enrichm_Validated_DGE_prot_SMED_BP

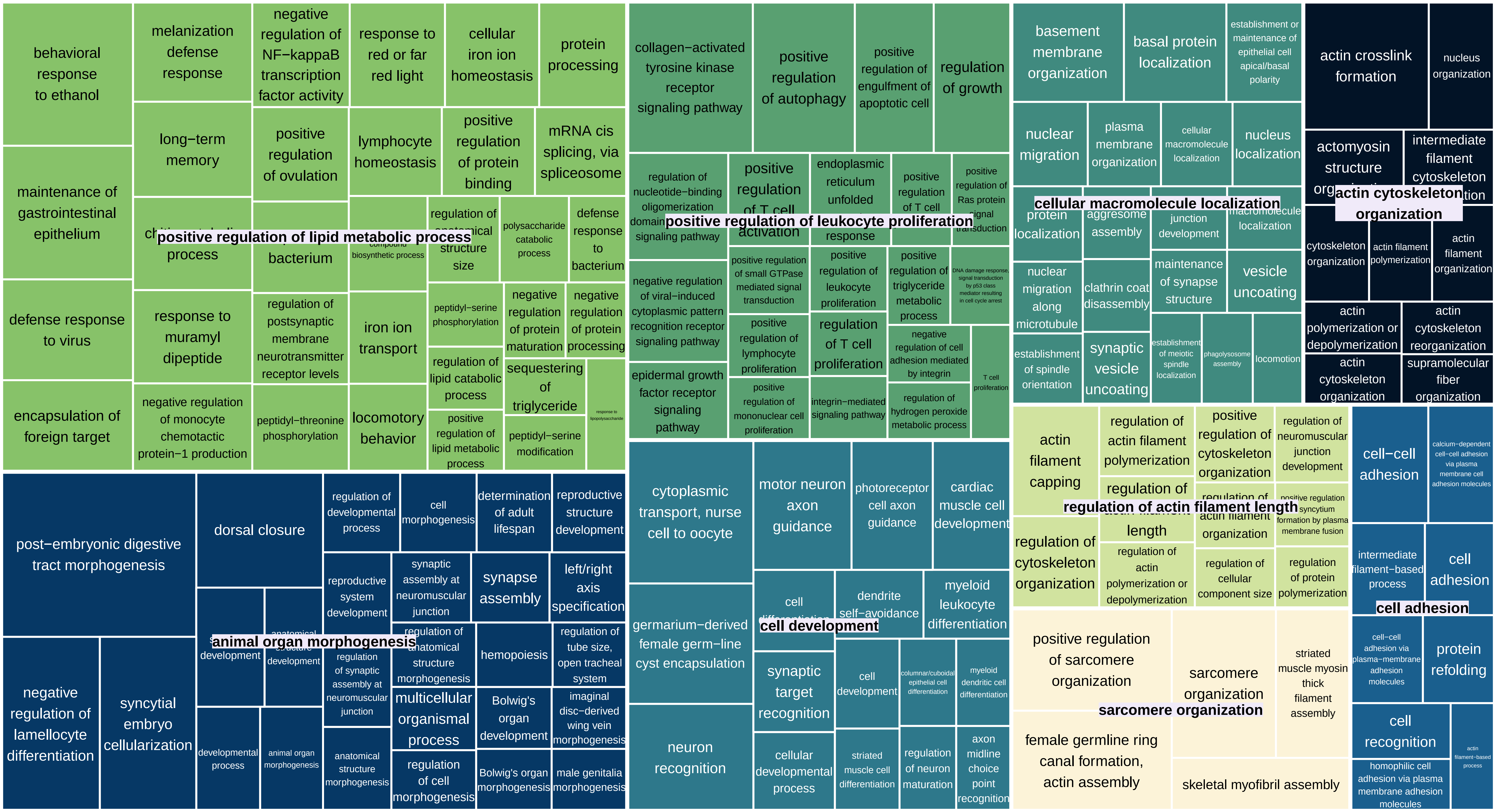

### Most significant pathways DGEs-prot SMED

link to full report: poner link en github

The following table shows the 25 most relevant pathways sorted by p-value.

| Pathway name | Entities | | | | Reactions | |
|---|---|---|---|---|---|---|
| | found | ratio | p-value | FDR* | found | ratio |
| Surfactant metabolism | 6 / 95 | 0.004 | 4.50e-06 | 0.004 | 11 / 29 | 0.002 |
| Formation of a pool of free 40S subunits | 5 / 111 | 0.005 | 1.41e-04 | 0.059 | 2 / 2 | 1.34e-04 |
| SRP-dependent cotranslational protein targeting to membrane | 5 / 119 | 0.005 | 1.95e-04 | 0.059 | 5 / 5 | 3.36e-04 |
| L13a-mediated translational silencing of Ceruloplasmin expression | 5 / 126 | 0.005 | 2.53e-04 | 0.059 | 2 / 3 | 2.02e-04 |
| Interaction between L1 and Ankyrins | 3 / 33 | 0.001 | 4.43e-04 | 0.072 | 4 / 4 | 2.69e-04 |
| FLT3 signaling by CBL mutants | 2 / 7 | 3.04e-04 | 4.59e-04 | 0.072 | 1 / 1 | 6.72e-05 |
| Modulation by Mtb of host immune system | 3 / 36 | 0.002 | 5.70e-04 | 0.076 | 2 / 6 | 4.03e-04 |
| Defective CSF2RB causes SMDP5 | 3 / 40 | 0.002 | 7.72e-04 | 0.08 | 1 / 1 | 6.72e-05 |
| Defective CSF2RA causes SMDP4 | 3 / 40 | 0.002 | 7.72e-04 | 0.08 | 1 / 1 | 6.72e-05 |
| GTP hydrolysis and joining of the 60S ribosomal subunit | 7 / 170 | 0.007 | 9.71e-04 | 0.083 | 3 / 3 | 2.02e-04 |
| Signal regulatory protein family interactions | 3 / 45 | 0.002 | 0.001 | 0.083 | 1 / 10 | 6.72e-04 |
| Myoclonic epilepsy of Lafora | 2 / 11 | 4.78e-04 | 0.001 | 0.083 | 1 / 2 | 1.34e-04 |
| Peptide chain elongation | 4 / 105 | 0.005 | 0.001 | 0.083 | 4 / 5 | 3.36e-04 |
| Diseases associated with surfactant metabolism | 3 / 48 | 0.002 | 0.001 | 0.083 | 3 / 7 | 4.70e-04 |
| Scavenging by Class A Receptors | 3 / 49 | 0.002 | 0.001 | 0.083 | 6 / 10 | 6.72e-04 |
| Eukaryotic Translation Termination | 4 / 109 | 0.005 | 0.001 | 0.083 | 3 / 5 | 3.36e-04 |
| Selenoamino acid metabolism | 5 / 191 | 0.008 | 0.002 | 0.089 | 3 / 33 | 0.002 |
| Selenocysteine synthesis | 4 / 115 | 0.005 | 0.002 | 0.09 | 2 / 7 | 4.70e-04 |
| Nonsense Mediated Decay (NMD) independent of the Exon Junction Complex (EJC) | 4 / 123 | 0.005 | 0.002 | 0.108 | 1 / 1 | 6.72e-05 |
| Maturation of protein E | 2 / 18 | 7.83e-04 | 0.003 | 0.131 | 1 / 5 | 3.36e-04 |
| Regulation of TLR by endogenous ligand | 3 / 65 | 0.003 | 0.003 | 0.131 | 2 / 13 | 8.73e-04 |
| Viral mRNA Translation | 4 / 136 | 0.006 | 0.003 | 0.131 | 2 / 2 | 1.34e-04 |
| Cap-dependent Translation Initiation | 7 / 225 | 0.01 | 0.003 | 0.131 | 12 / 18 | 0.001 |
| Eukaryotic Translation Initiation | 7 / 228 | 0.01 | 0.003 | 0.135 | 14 / 21 | 0.001 |

| Pathway name | Entities | | | | Reactions | |
|---|---|---|---|---|---|---|
| | found | ratio | p-value | FDR* | found | ratio |
| Translesion synthesis by POLI | 2 / 22 | 9.57e-04 | 0.004 | 0.149 | 3 / 3 | 2.02e-04 |

* False Discovery Rate

link to Blast results ONUN: poner link en github

link to Blast results SMED: poner link en github
