## Supplementary Material 6 for "Genomic exaptation and regulatory landscape shifts as key mechanisms enabling flatworm terrestrialization"

**Supplementary Material 6.** Enrichments Differentially Expressed Genes (DEGs) under selection. Treemaps showing enriched GO terms (BP: Biological Processes, MF: Molecular Functions, CC: Cellular Components) in DGEs arisen in Tricladida (node 65), Continenticola (node64), Geoplanoidea (node 63), and Geoplaniidae (node 62) with shift selection signal

TreeMap_Enrichm_DGEs_ONUN_Sel_node65_BP

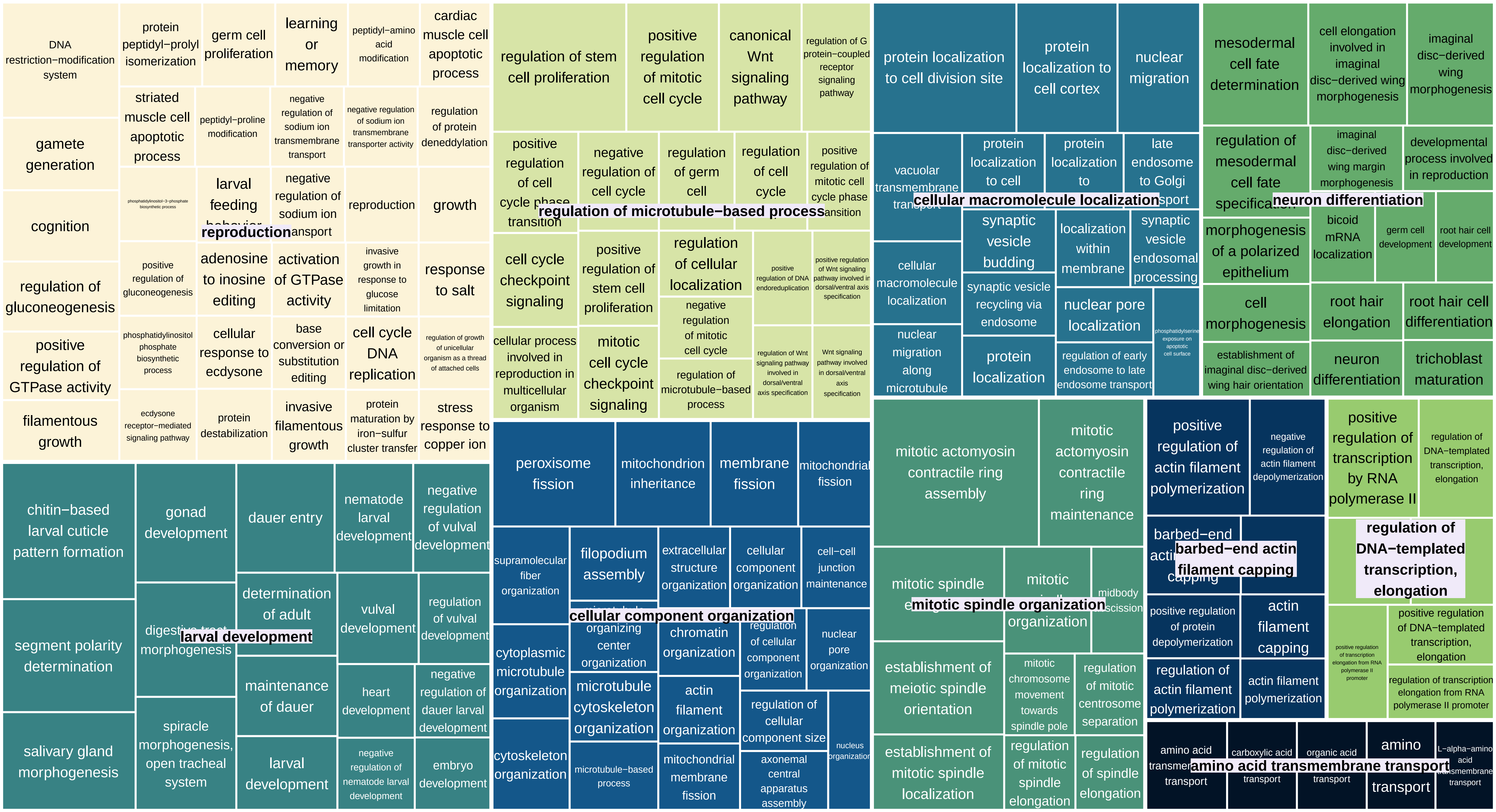

TreeMap_Enrichm_DGEs_ONUN_Sel_node65_MF

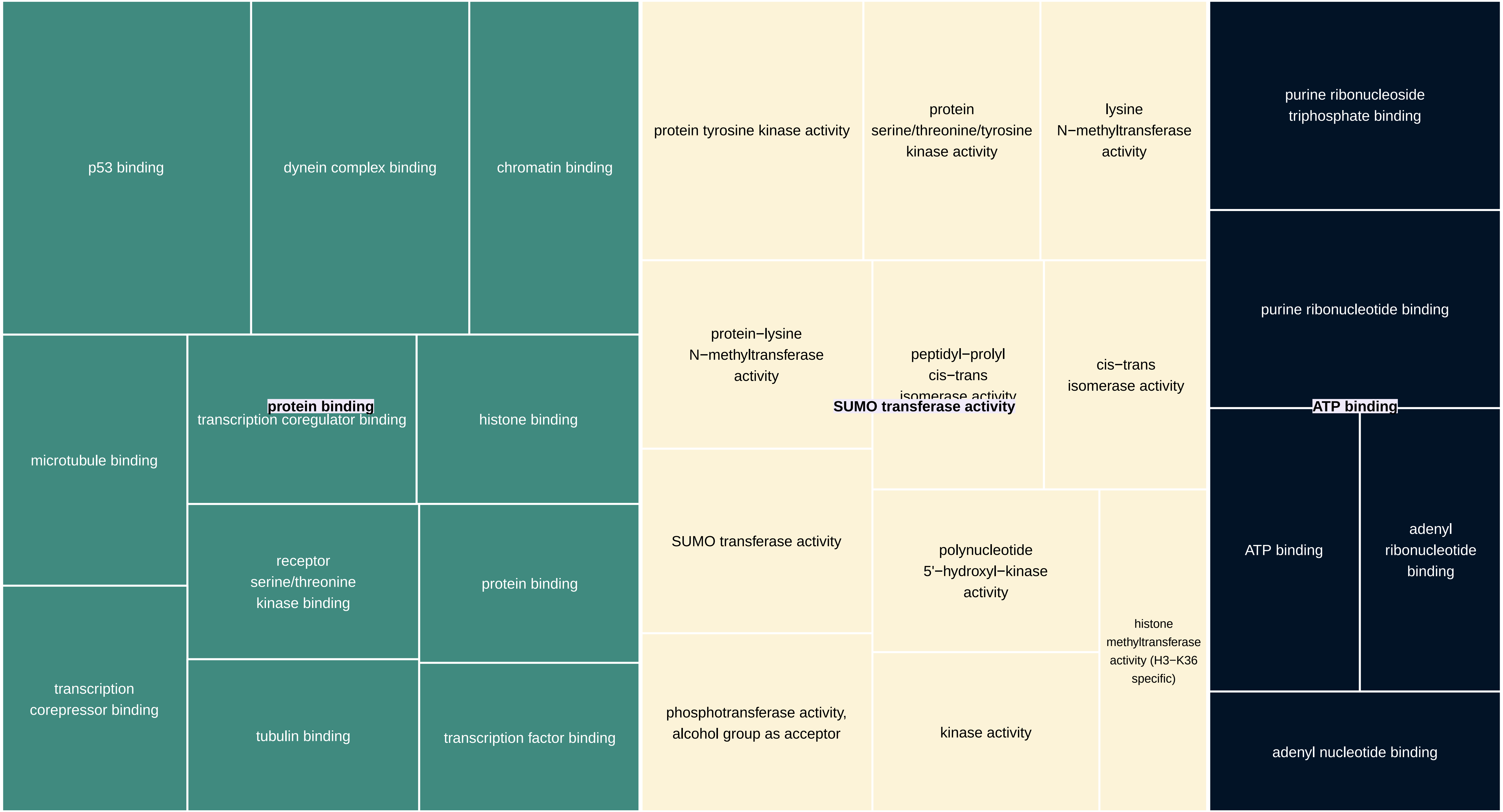

TreeMap_Enrichm_DGEs_ONUN_Sel_node65_CC

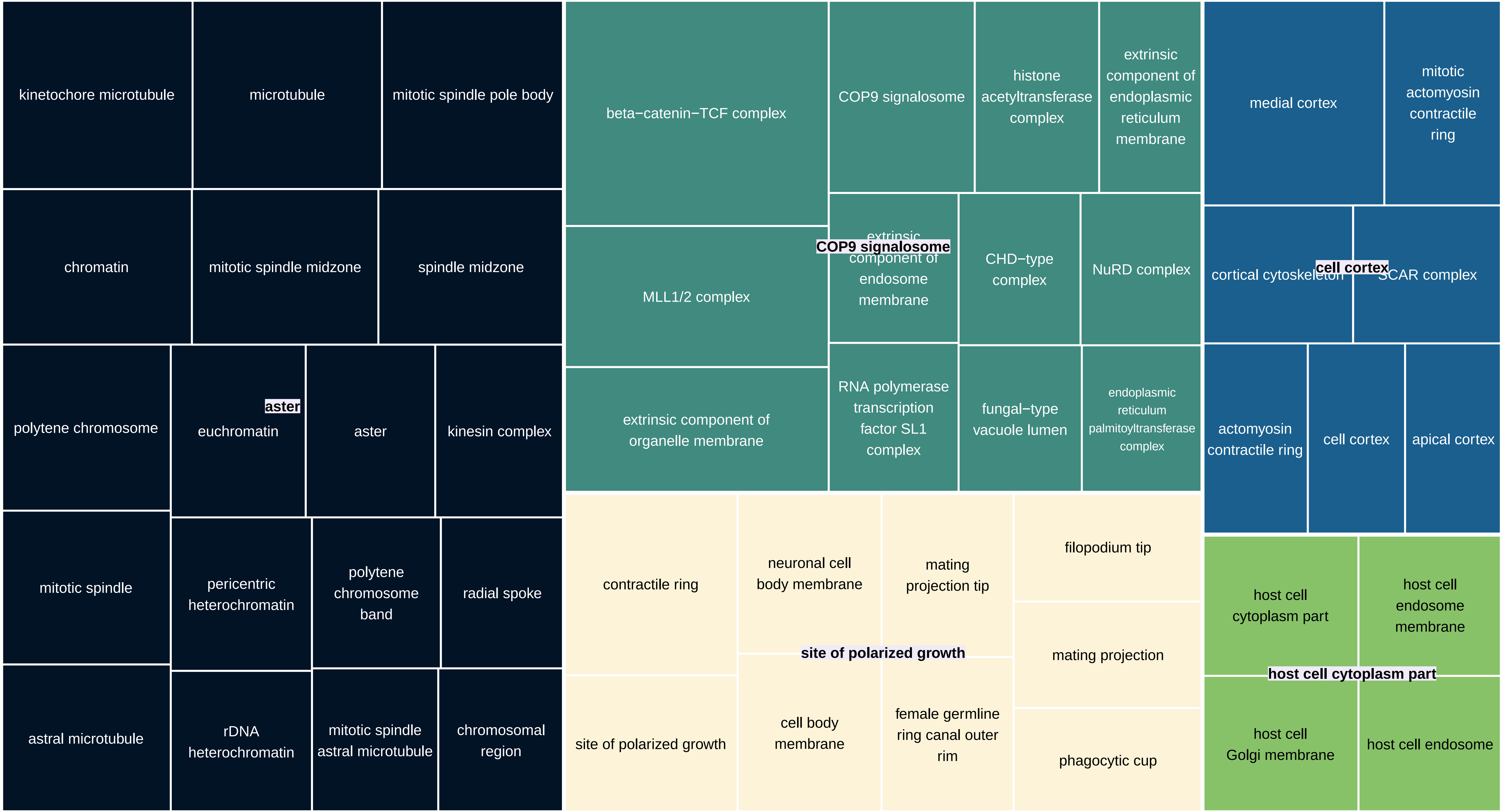

TreeMap_Enrichm_DGEs_ONUN_Sel_node64_BP

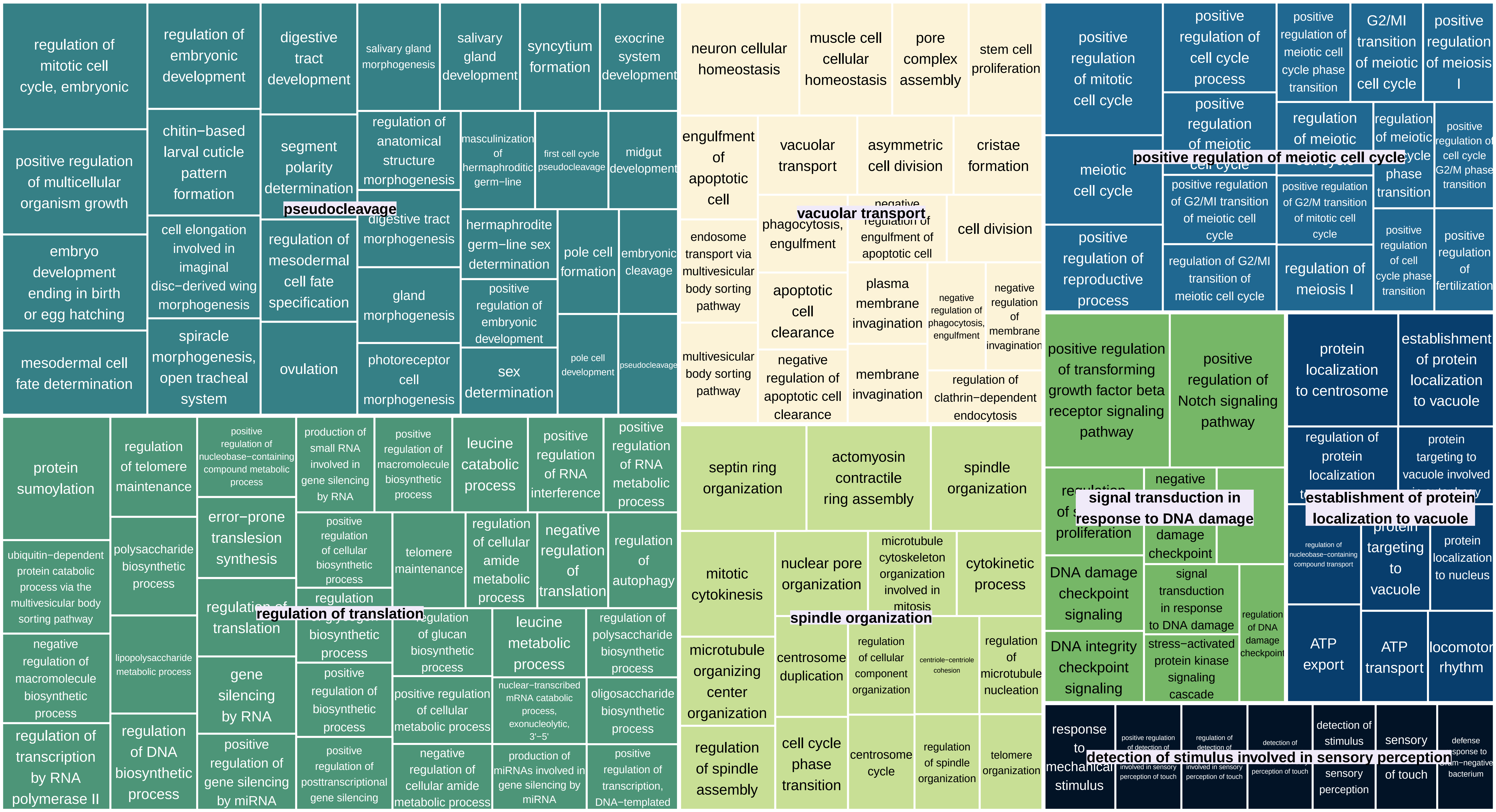

TreeMap_Enrichm_DGEs_ONUN_Sel_node64_MF

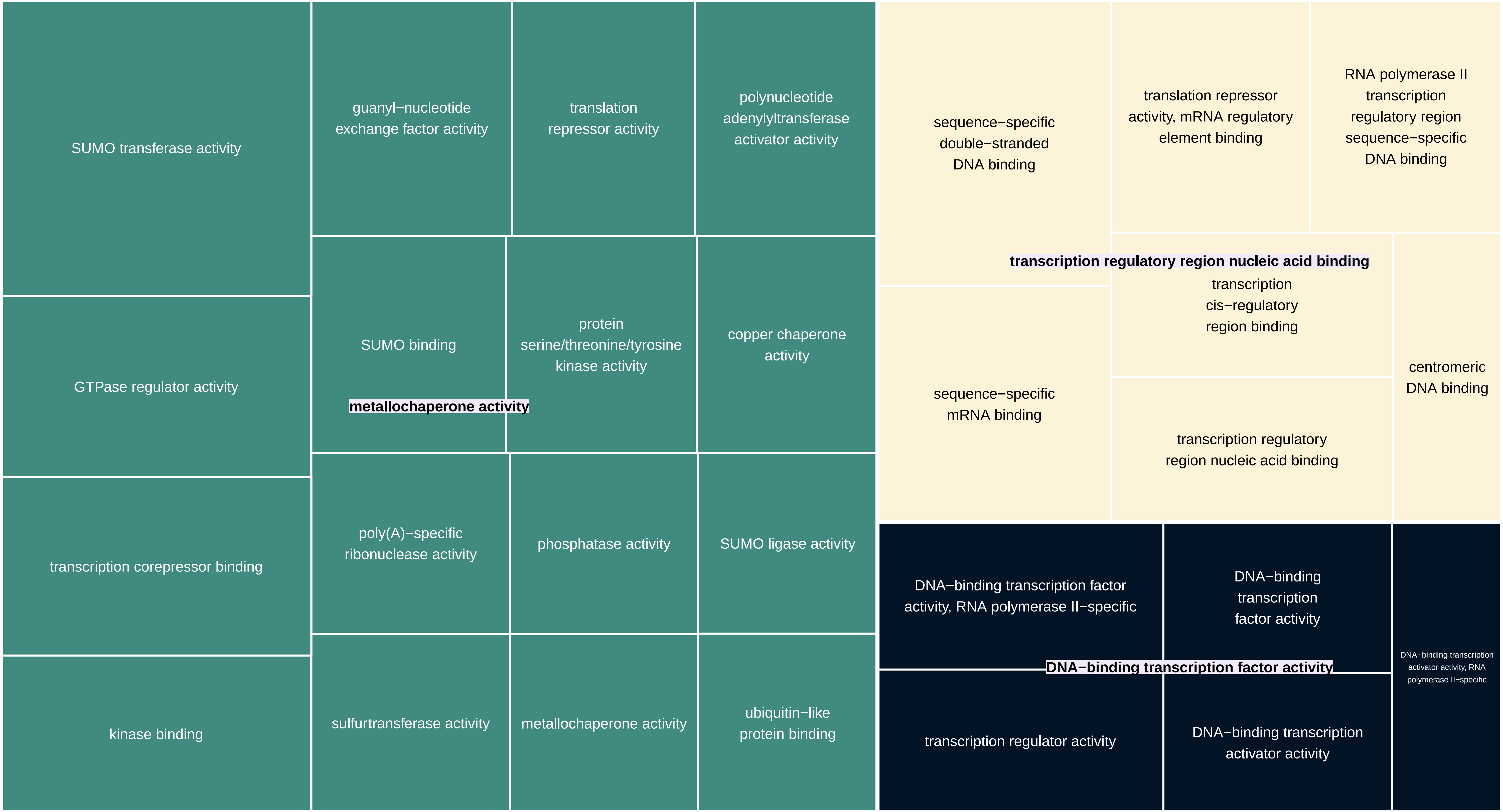

TreeMap_Enrichm_DGEs_ONUN_Sel_node64_CC

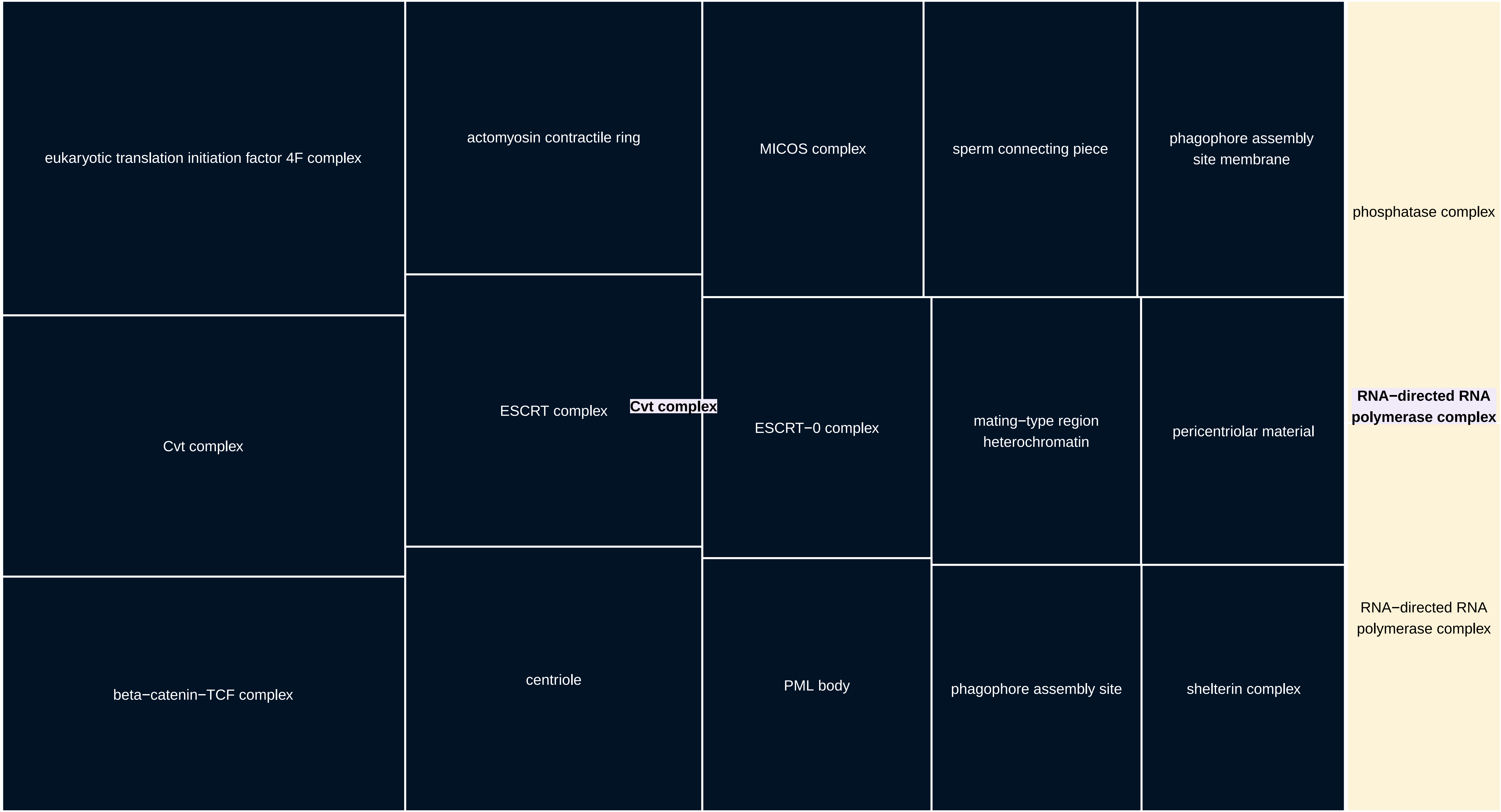

TreeMap_Enrichm_DGEs_ONUN_Sel_node63_BP

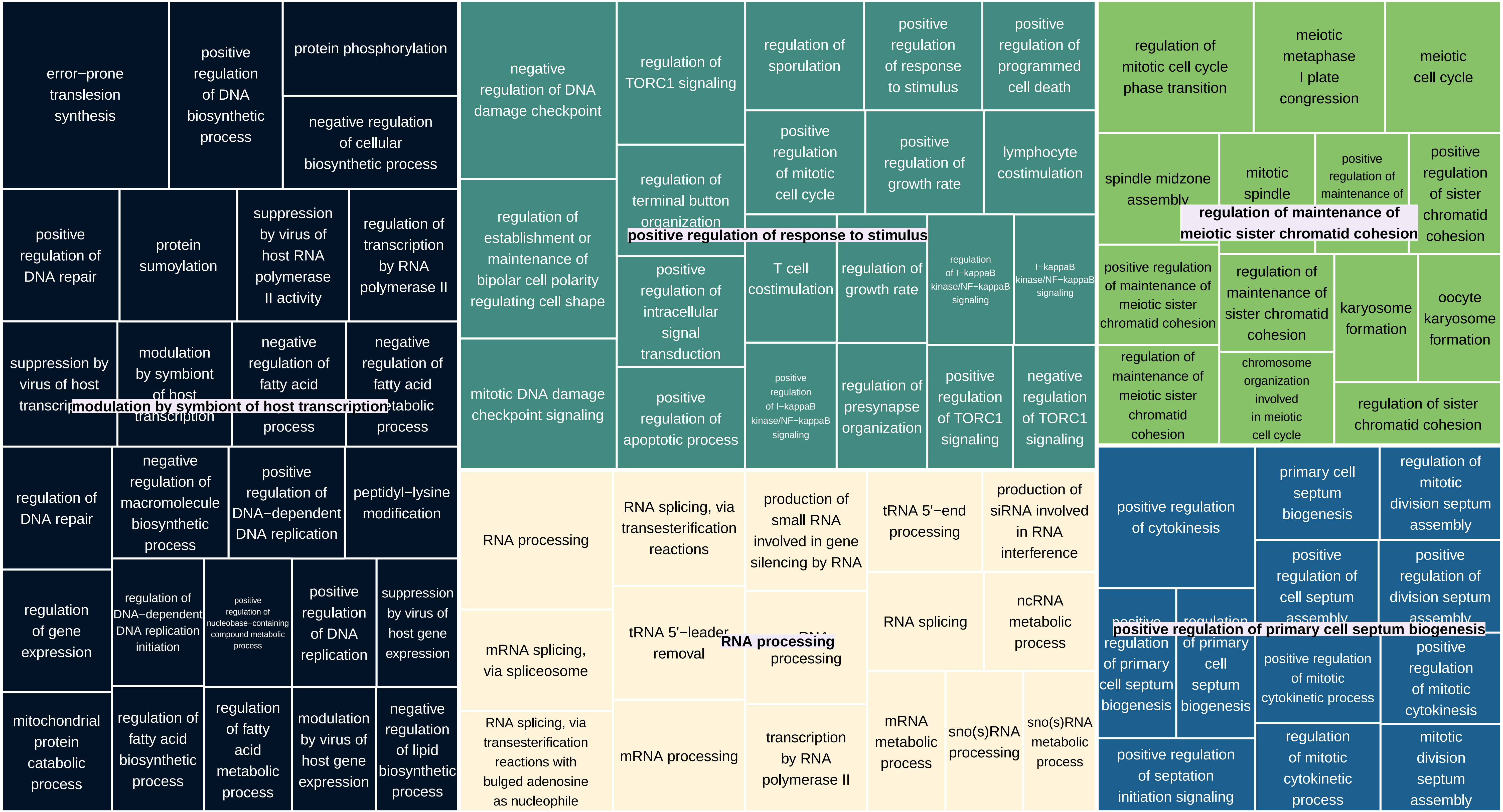

TreeMap_Enrichm_DGEs_ONUN_Sel_node63_MF

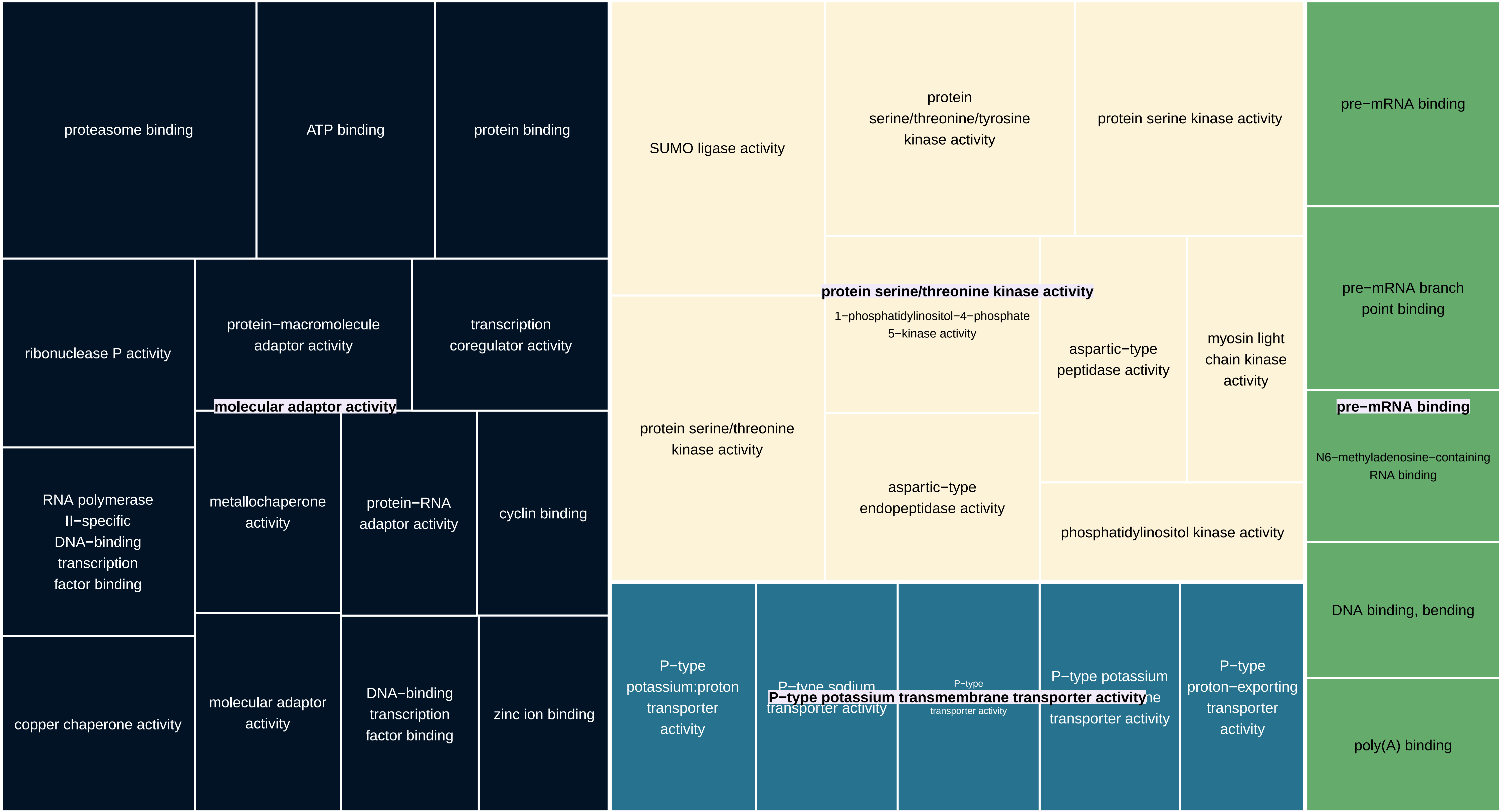

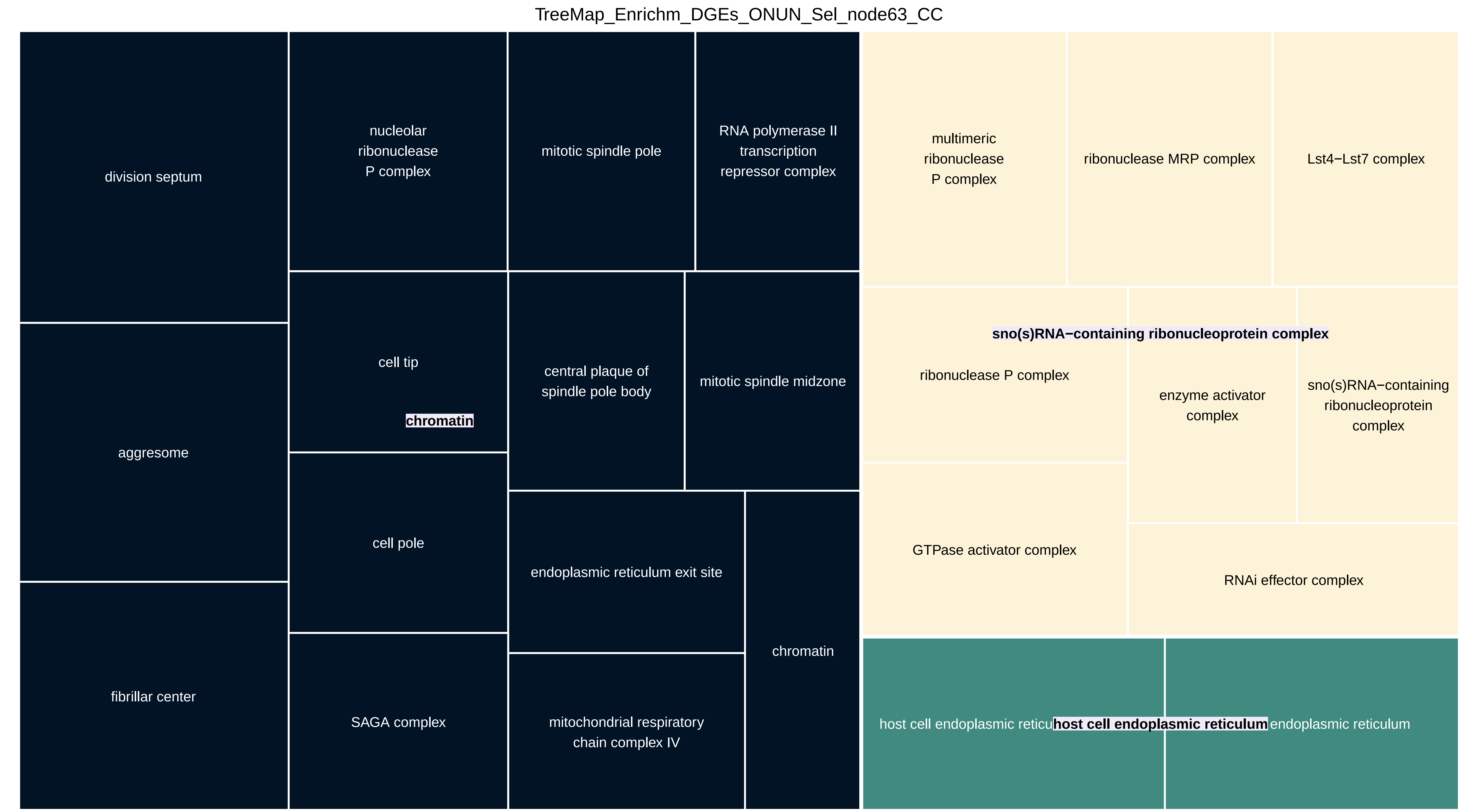
TreeMap_Enrichm_DGEs_ONUN_Sel_node63_CC
division septum
nucleolar ribonuclease P complex
mitotic spindle pole
RNA polymerase II transcription repressor complex
multimeric ribonuclease P complex
ribonuclease MRP complex
Lst4−Lst7 complex
sno(s)RNA−containing ribonucleoprotein complex
aggresome
cell tip
chromatin
central plaque of spindle pole body
mitotic spindle midzone
ribonuclease P complex
enzyme activator complex
sno(s)RNA−containing ribonucleoprotein complex
cell pole
endoplasmic reticulum exit site
GTPase activator complex
RNAi effector complex
fibrillar center
SAGA complex
mitochondrial respiratory chain complex IV
chromatin
host cell endoplasmic reticu
host cell endoplasmic reticulum
endoplasmic reticulum
