## Supplementary Material 7 for "Genomic exaptation and regulatory landscape shifts as key mechanisms enabling flatworm terrestrialization"

TreeMap_Enrichm_DGEs_ONUN_Oxygen_VL_exp_node65_BP

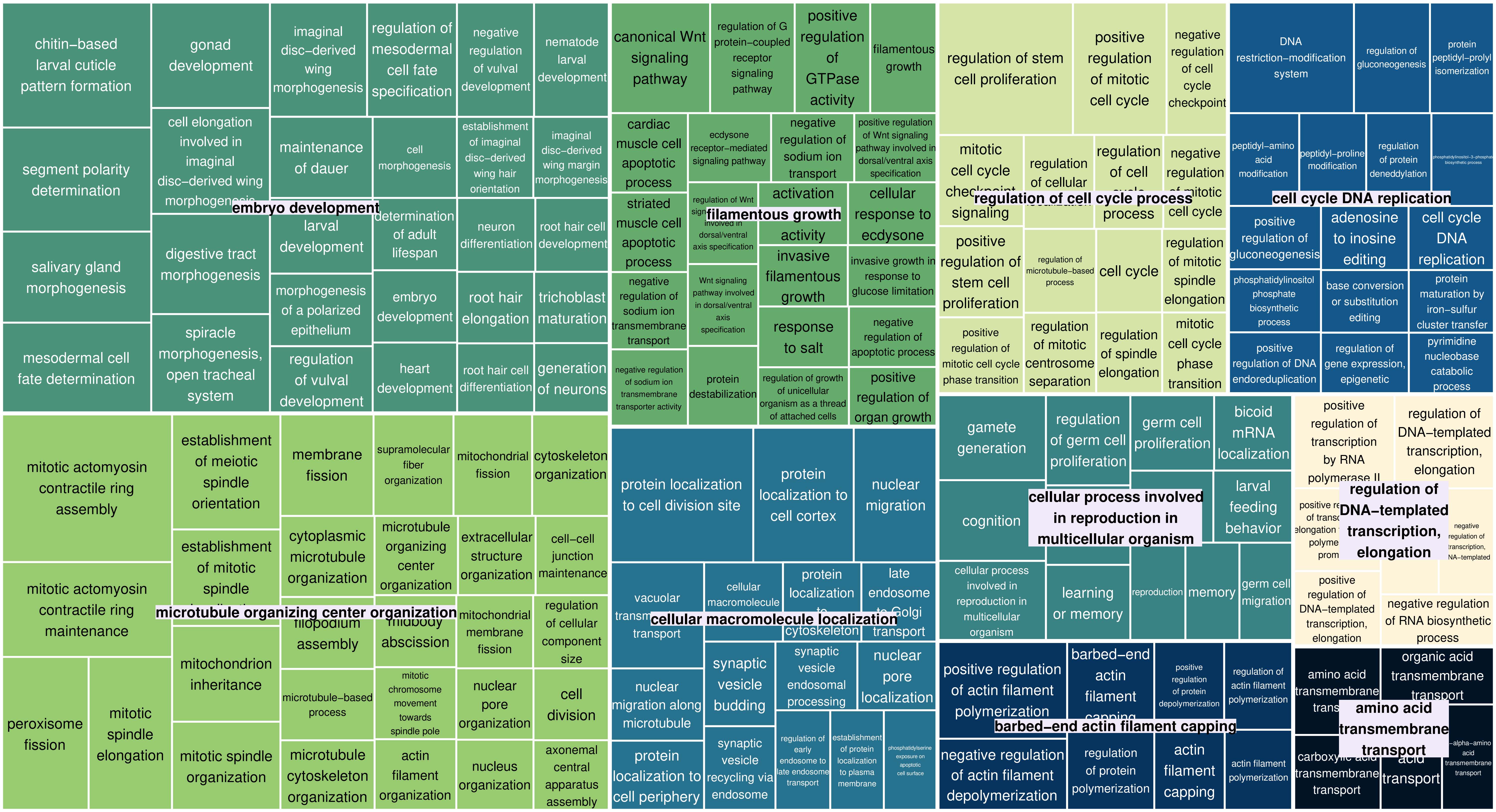

TreeMap_Enrichm_DGEs_ONUN_Oxygen_VL_exp_node64_BP

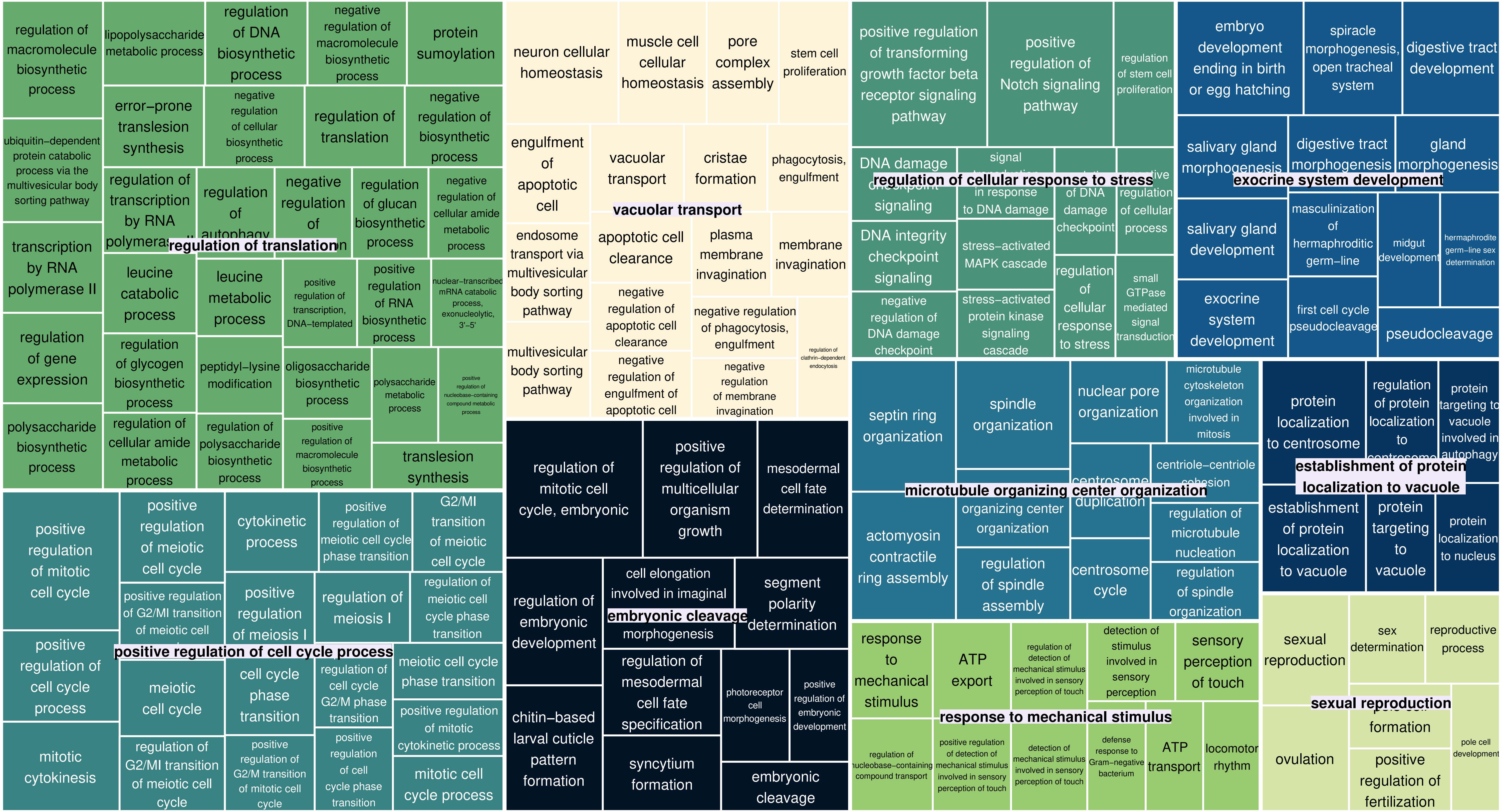

TreeMap_Enrichm_DGEs_ONUN_Oxygen_VL_exp_node63_BP

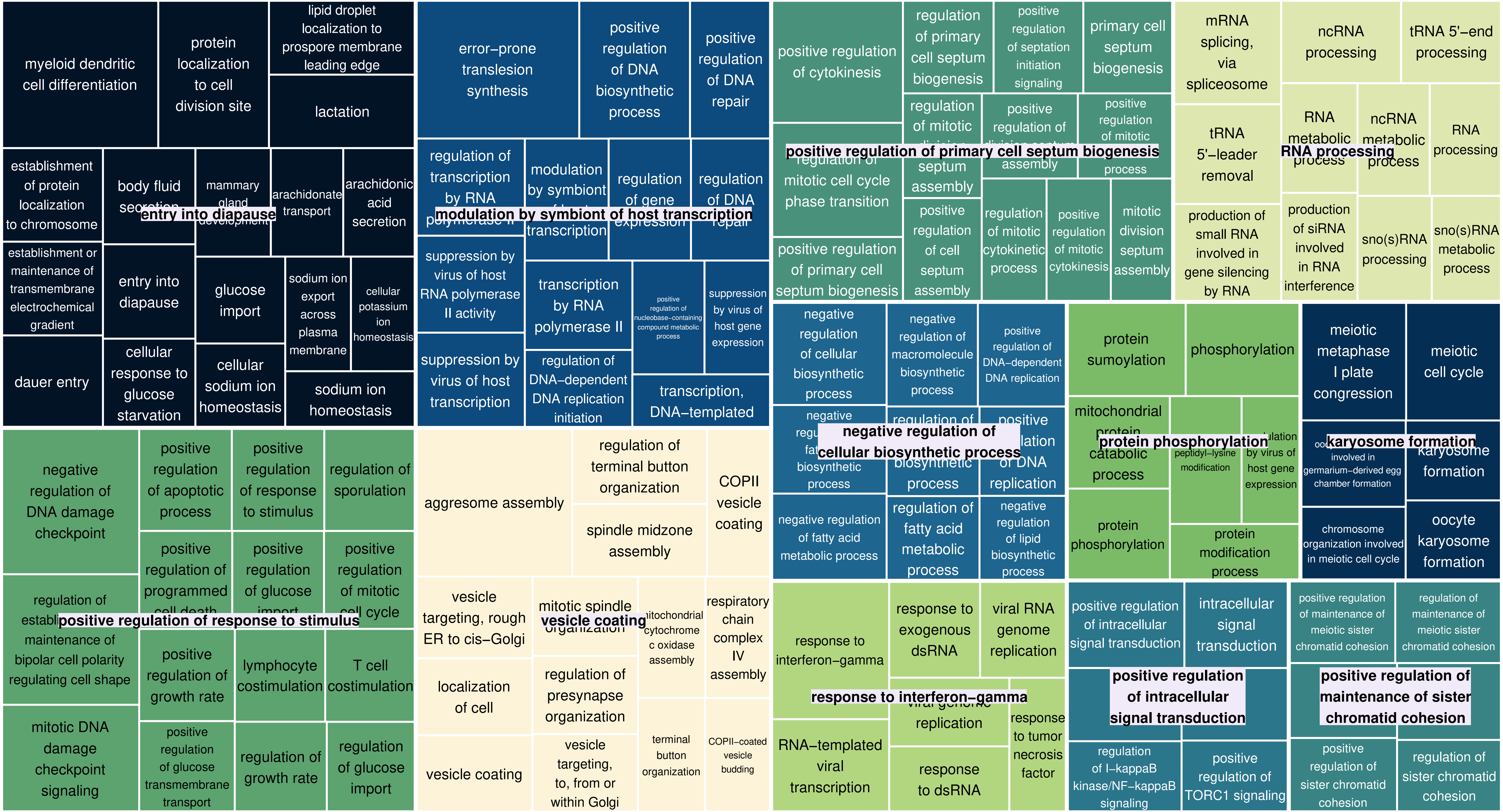
